# Novel Comparison of Evaluation Metrics for Gene Ontology Classifiers Reveals Drastic Performance Differences

**DOI:** 10.1101/427096

**Authors:** Ilya Plyusnin, Liisa Holm, Petri Törönen

## Abstract

Automated protein annotation using the Gene Ontology (GO) plays an important role in the biosciences. Evaluation has always been considered central to developing novel annotation methods, but little attention has been paid to the evaluation metrics themselves. Evaluation metrics define how well an annotation method performs and allows for them to be ranked against one another. Unfortunately, most of these metrics were adopted from the machine learning literature without establishing whether they were appropriate for GO annotations.

We propose a novel approach for comparing GO evaluation metrics called *Artificial Dilution Series* (ADS). Our approach uses existing annotation data to generate a series of annotation sets with different levels of correctness (referred to as their signal level). We calculate the evaluation metric being tested for each annotation set in the series, allowing us to identify whether it can separate different signal levels. Finally, we contrast these results with several *false positive annotation sets*, which are designed to expose systematic weaknesses in GO assessment.

We compared 37 evaluation metrics for GO annotation using ADS and identified drastic differences between metrics. We show that some metrics struggle to differentiate between different signal levels, while others give erroneously high scores to the false positive data sets. Based on our findings, we provide guidelines on which evaluation metrics perform well with the Gene Ontology and propose improvements to several well-known evaluation metrics. In general, we argue that evaluation metrics should be tested for their performance and we provide software for this purpose (https://bitbucket.org/plyusnin/ads/). ADS is applicable to other areas of science where the evaluation of prediction results is non-trivial.

**Author Summary:** In the biosciences, predictive methods are becoming increasingly necessary as novel sequences are generated at an ever-increasing rate. The volume of sequence data necessitates Automated Function Prediction (AFP) as manual curation is often impossible. Unfortunately, selecting the best AFP method is complicated by researchers using different evaluation metrics. Furthermore, many commonly-used metrics can give misleading results. We argue that the use of poor metrics in AFP evaluation is a result of the lack of methods to benchmark the metrics themselves. We propose an approach called Artificial Dilution Series (ADS). ADS uses existing data sets to generate multiple artificial AFP results, where each result has a controlled error rate. We use ADS to understand whether different metrics can distinguish between results with known quantities of error. Our results highlight dramatic differences in performance between evaluation metrics.

## Introduction

The biosciences generate sequences at a faster rate than their structure, function or interactions can be experimentally determined. This has created a demand for computational methods that can automatically link novel sequences to their biological function [1], resulting in the development of many Automated Function Prediction (AFP) methods [2, 3]. The comparison of different methods is therefore an important task for bioscientists who need to use AFP methods in sequencing projects, for bioinformaticians who develop AFP methods and for reviewers who need to peer-review AFP-related publications.

Evaluating AFP tools is challenging. Articles use different data sets and report different evaluation metrics in their results. This makes it unclear how different methods perform and sets a bad example for method developers. Standard data sets, such as the MouseFunc project [4] and CAFA competitions [2, 3, 5], are starting to emerge. In our own work, we have additionally used filtered sets of well-annotated sequences [6, 7]. Unfortunately, there are no clear standards for which Evaluation Metrics (EvM) to use. An EvM is a function that quantifies the performance of a method being tested [3, 8–10] and, as Table 1 in supplementary S2 Text demonstrates, there is no consensus in the literature for which are best. Furthermore, although the CAFA competitions [2, 3, 5] have led to defacto standards, these have been criticised. Gillis and Pavlidis [11], for example, observed that *“the primary performance metric* … *for CAFA is unsatisfactory*. … *by this measure, a null ‘prediction method’ outperforms most methods*.*”* Kahanda et al. [12] pointed out that the ranking of methods in CAFA varied considerably between different EvMs. Even the CAFA authors themselves concede that finding appropriate EvMs for GO annotation is an open problem [2, 3]. This shows that there is a clear need for research on AFP-related EvMs.

**Table 1.**
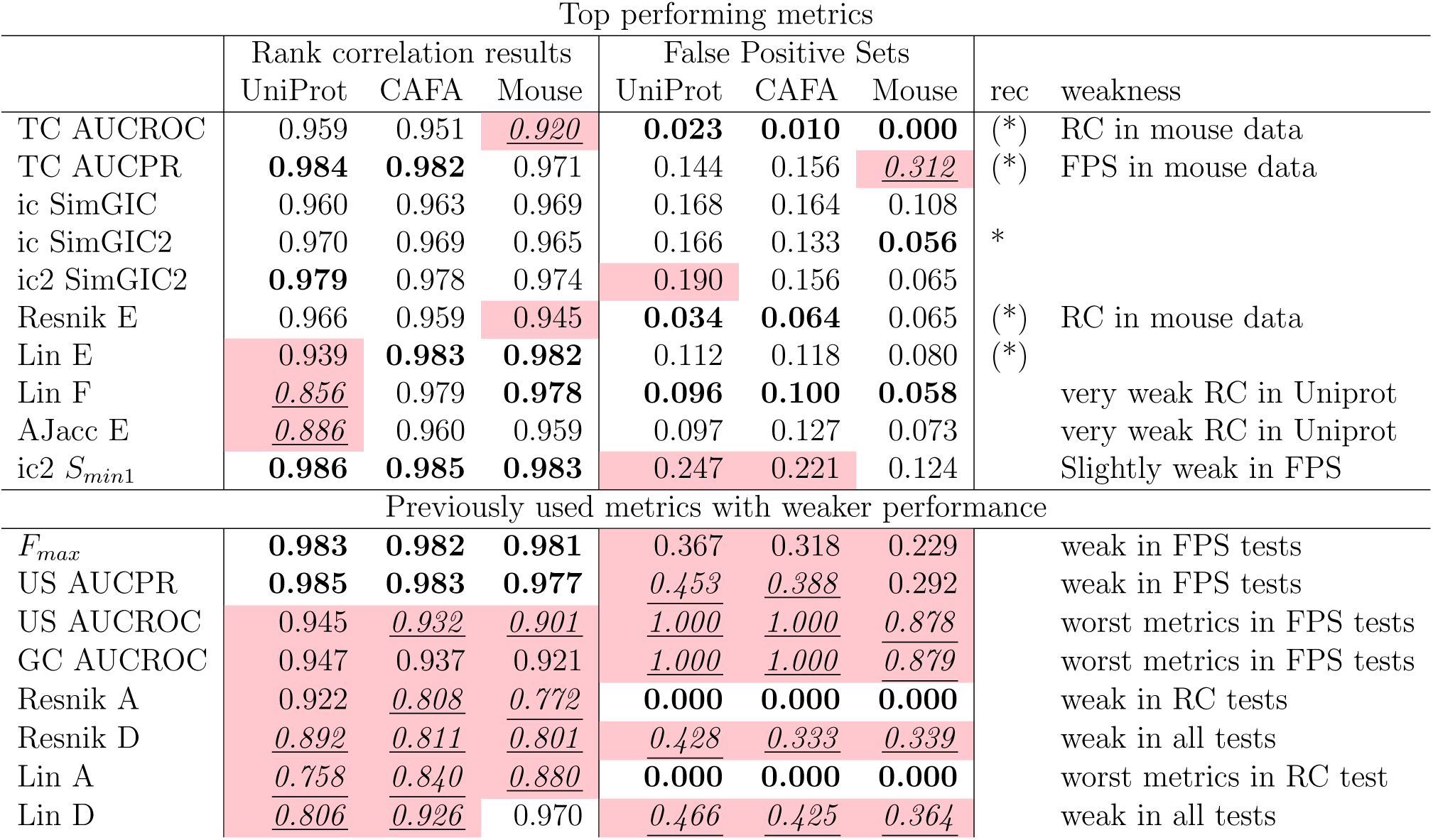
Summary of results for best performing and widely-used metrics. Here we show RC (Rank Correlation) and FP (False Positive) results for the best performing methods. We also show same results for some widely-used metrics. Good metrics should have a high RC score and low FP scores. Rec column shows our selected recommendations (See text for details). The five best results in each column are shown in bold. The five weakest results in each column are shown with underlined italics. Metrics that fail a given test are highlighted in red (see text for details). Note how methods in lower block show consistent weak performance either in RC or FP tests.

The challenges that an EvM must overcome are related to the structure of the Gene Ontology [13] itself:

- The GO structure has a large number of classes and genes can belong to many of them. This creates a multilabel and multiclass classification task.
- The hierarchy of the GO structure causes strong and complex correlations between classes.
- Class sizes vary widely, with most being very small. This causes strong class size imbalance. and the nature of biological data:
- The set of genes used for evaluation can be small compared to the number of classes [2], making the table of correct annotations sparse.
- The meaning and definition of true negative GO annotations is ambiguous [14, 15].
- Each gene has a varying number of correct annotations.

These challenges point to the selection of EvM as clearly nontrivial. Furthermore, the EvM selection problem is not limited to AFP comparison, but is a problem in many other fields [8, 9], such as in speech recognition [16] and with the classification of cognitive activities [17]. Finally, EvMs often forms the core of the results section in bioinformatics articles and can be selected to favour one’s own method [18].

We propose a novel approach, **Artificial Dilution Series (ADS)**, to address these challenges. ADS checks the performance and stability of any EvM using real-life GO annotations. ADS creates artificial classifier results by taking a set of correct GO predictions and replacing a controlled percentage of correct annotations with errors (creating type 1 and type 2 errors, collectively referred to as “noise”). The noise percentage is then increased in a step-wise fashion, creating a set of separate result data sets with a controlled level of noise at each step. This process creates a “dilution series” of the original signal in a collection of altered data sets. This series is then used to test different EvMs to see how well they separate data sets with different signal levels.

Additionally, we included a secondary test using several **False Positive (FP) data sets**. We run each EvM with each FP data set, and compare them with the ADS results. A good EvM would place FP sets close to sets with zero signal. FP sets allow us to monitor how each EvM ranks, for example, the naïve predictor [2, 3], which is a known issue for some EvMs [11]. Our supplementary text S2 Text compares our approach to related research.

We tested many EvMs previously applied to GO annotations using three datasets: CAFA1 [2], MouseFunc [4] and 1000 randomly selected GO annotated sequences from the UniProt database. We also tested small variations of the existing EvMs. Our results show that *1*) many popular EvMs often fail the FP tests, *2*) semantic scores [19] are heavily affected by what summation method is used, with methods failing either in ADS or FP tests, *3*) The best, most consistent performance across all the data sets was obtained with modified SimGIC functions (a weighted Jaccard correlation [20]) and *4*) term centric analysis or Information Content (IC) weights are usually required for good performance. IC weights are discussed in methods and in S2 Text.

Some EvMs showed better performance than SimGIC on one or two data sets. This suggests that different GO data sets have different challenges, each requiring separate evaluation. The ADS software, all tested EvMs and our analysis scripts are freely available online^1,2^.

## Materials and Methods

### Generating Artificial Dilution Series

#### Motivation and Requirements

We assume that we have a set of genes, each with a varying number of GO annotations. This set of annotated genes will be referred to as the truth set, *T*. *T* is expected to be a table that contains gene names in the first column and the associated GO term identifiers in the second column. We assume the existence of an ontology, such as GO. The ontology links smaller, more precise child terms to more broadly defined ancestral terms. Here, if a term *t*_*child*_, is a child term for *t*_*par*_, then all genes that belong to *t*_*child*_ will also belong to *t*_*par*_ (i.e. *t*_*child*_ ⊆ *t*_*par*_). These child–ancestor connections form a directed acyclic graph, where every GO term, with the exception of the root term, has a varying number of parent terms [13]. Genes can be mapped to GO terms at varying levels in the ontology, depending on how detailed our level of knowledge is.

Artificial Dilution Series creates several data sets, each representing functional classifications for genes in *T*. These data sets, referred to here as *Artificial Prediction Sets* (or AP sets), imitate a GO prediction set from a GO classifier. They are comprised of a list of triples, (gene, GO, *sc*), where *sc* is the artificial prediction score. Each AP set is created artificially, with a defined proportion of correct and erroneous annotations. AP set creation does not require a real classifier, nor the actual gene sequences. Furthermore, each AP set can also be defined as a table, where rows represents different annotation entities and columns represent genes, GO terms and classifier scores.

The workflow for the ADS pipeline is shown in Fig. 1 and outlined in pseudocode in S1 Text. The first step is to create two GO prediction sets, the *Positive Prediction set, P*_*pos*_, and the *Negative Prediction set, P*_*neg*_. These steps use a *signal model* to perturb the ground truth GO annotations and a *noise model* to generate erroneous annotation predictions. Note that both *P*_*pos*_ and *P*_*neg*_ have a table structure, similar to the AP set.

**Fig 1.**
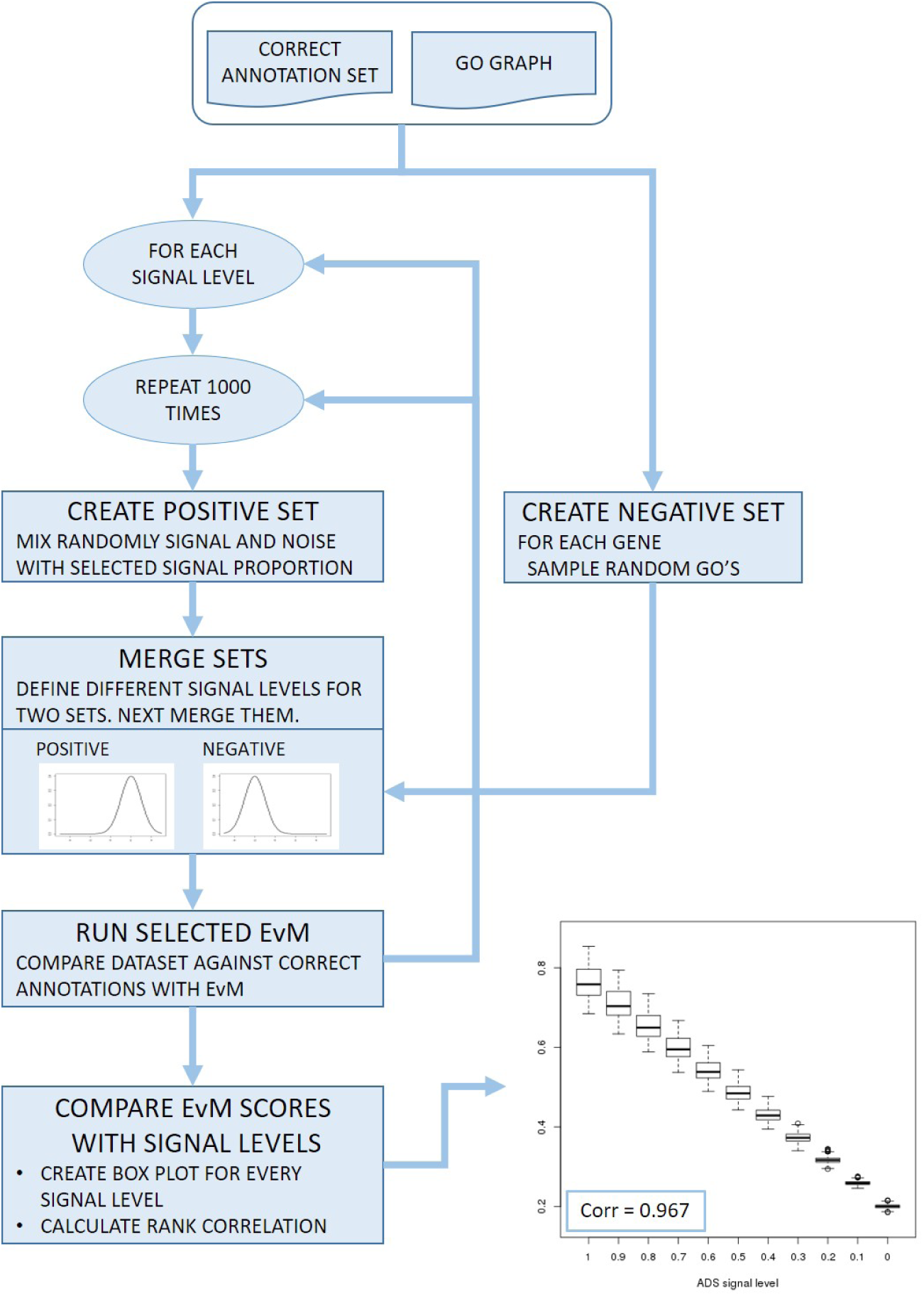
ADS workflow. The ADS pipeline iterates through specified signal levels (e.g. from 100% to 0%) at each level creating a collection of artificial prediction sets (AP sets). Each AP set is a union of a set containing only false annotations (negative set) and a set containing a controlled fraction of true annotations (positive set). AP sets are compared against the correct annotations using an Evaluation Metric (EvM). The results can be shown graphically by plotting EvM scores for AP sets at each signal level as boxplots. These reveal how stable an EvM is against random variation introduced at different signal levels and whether it can measure the amount of signal retained in different AP sets. Finally, EvM performance is quantified with rank correlation.

#### Step 1: Create the Positive Set

The Positive set, *P*_*pos*_, is created from a copy of *T* in two stages: First, we draw a random integer, *N*_*shift*_, between 0 and the size of *P*_*pos*_, and select *N*_*shift*_ random rows from *P*_*pos*_. Next, the GO term, *t*_*corr*_, of every selected row is replaced with a semantically similar term. This term is selected from the *k* nearest parents of *t*_*corr*_. We refer to this procedure as *shifting to semantic neighbours* and it forms the basis of our signal model. Figure 1 in supplementary text S2 Text demonstrates this step. We have tested *k* = (2, 3, 4) and found results to be similar.

Second, we introduce a percentage of errors, *ϵ*, by switching GO terms between genes for a controlled fraction of annotation lines (see Fig 2 for a toy example). We refer to this as the *permutation of the positive set*. Here a random pair of annotations, *a* = (*gene*_*a*_, *t*_*a*_) and *b* = (*gene*_*b*_, *t*_*b*_), where *gene*_*a*_ ≠ *gene*_*b*_ and *t*_*a*_ ≠ *t*_*b*_, is selected from *P*_*pos*_. If *t*_*a*_ is sufficiently far away in the GO structure from every GO term annotated to *gene*_*b*_ and vice versa for *t*_*b*_ and the annotations of *gene*_*a*_, then the GO labels are exchanged. The distance between *t*_*x*_ and *t*_*y*_ is defined as the Jaccard correlation between the two sets created from the ancestral classes of *t*_*x*_ and *t*_*y*_ (see S1 Text). In our experiments we used a threshold, *th*_*noise*_, of 0.2 and required that the correlation was lower than *th*_*noise*_. This step represents our noise model.

**Fig 2.**
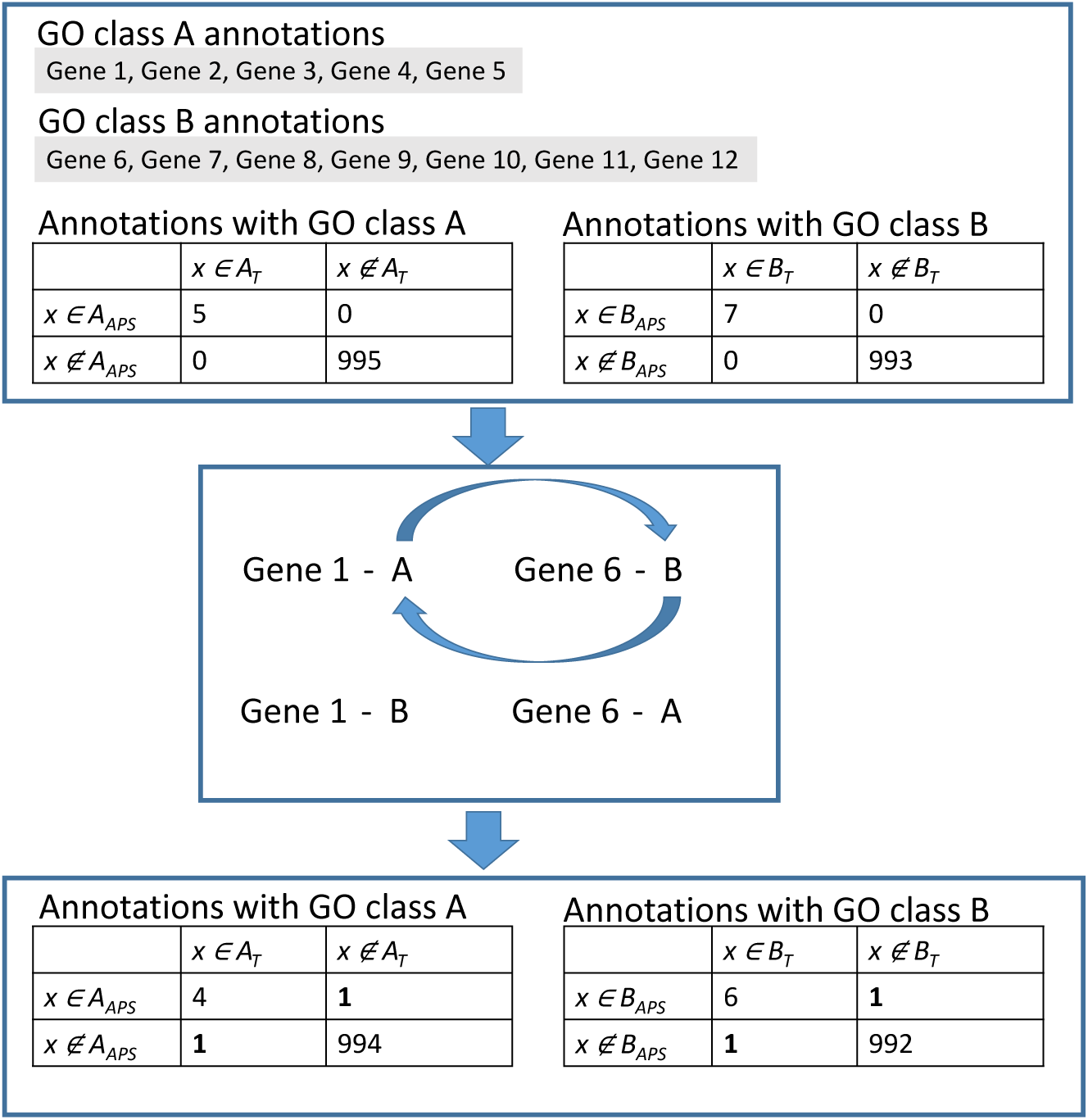
ADS permutation example. In this toy example we start with a set of correct annotations using only two GO terms A and B, taken from truth set *T*. Initially, in the upper part the Artificial Prediction Set (*APS*) matches perfectly to *T*. This would give us 100% correct predictions. Next, the permutation step, shown in the middle, switches GO terms for a pair of genes. This increases false positives and false negatives by 1 and decreases true positives and true negatives by 1. We obtain as a result an *APS* with a known level of errors (0.002 in this example). The permutation is repeated with other gene pairs and other GO terms until the required noise level is obtained.

The permutation process is repeated until the required percentage of errors, *E*, have been introduced. This adds a controlled fraction of false positives and false negatives among the predictions. We refer to this fraction as the *ADS noise level* and the remaining fraction of correct positive annotations as the *ADS signal level* (1 – ADS noise level).

#### Step 2: Create the Negative Set

The negative set, *P*_*neg*_, is created by assigning genes in *T* with random GO annotations. Here, for every unique gene name, *gene*_*x*_, that occurs in *T*, we sample a random GO term, *t*_*r*_, from the GO structure. As in the construction of the positive set, (*gene*_*x*_, *t*_*r*_) is included in *P*_*neg*_ only if it is located far away in the GO structure compared to all GO terms linked to *gene*_*x*_. This process is repeated until four erroneous GO annotations have been added to each gene.

#### Step 3: Add Prediction Scores and Merge Sets

All entries in *P*_*pos*_ and *P*_*neg*_ are assigned prediction scores, *sc*. Predictions in *P*_*pos*_ are assigned scores by sampling *sc* from a normal distribution with *mean* = 1 and *SD* = 0.5. This creates a triple (*gene*_*x*_, *t*_*x*_, *sc*) for each annotation in *P*_*pos*_. For predictions in *P*_*neg*_, we sample *sc* from a normal distribution with a lower mean (*mean* = *-*1 and *SD* = 0.5) and add it similarly to *P*_*neg*_.

If scores need to be within a bounded range, such as [0, 1], a sigmoid function could be used to scale the score values. Next, *P*_*pos*_ and *P*_*neg*_ are merged to form an AP set, *APS* = *P*_*pos*_ *∪ P*_*neg*_. What we now have is a set of GO predictions with A) a set of errors, represented by *P*_*neg*_, that can be separated by setting a threshold on *sc* values and B) a set of errors embedded into *P*_*pos*_, that cannot be separated with a threshold (ADS noise level). Furthermore, A stays constant for all AP sets, whereas B is adjusted with *ϵ*. Finally, the predictions are extended to include ancestor nodes in the GO hierarchy for a given metric. In some cases, such as semantic similarity-based metrics, the metric will already account for dependencies between GO terms [19] and this step is omitted.

#### Step 4: Creating AP Sets with All ADS Signal Levels

Full ADS analysis repeats the AP set creation, while incrementally changing the ADS signal level over the selected signal range, *SR* = [*x*_1_, *x*_2_…*x*_*k*_] (illustrated by the outer loop in Fig 1). We used an *SR* that spans from 100% to 0% signal in steps of ten percentage points. However, our implementation allows for any range with any number of steps. At each signal level the AP set generation is repeated *N* times, each time applying the shifting to semantic neighbourhood and permutation procedures as described above (the inner loop in Fig 1). The final ADS series will contain:

1. Evenly sampled ADS signal levels between the selected minimum and maximum signal levels.
2. *N* separate AP sets representing each signal level.
3. The proportion of annotations shifted to semantic neighbours varying randomly across AP sets at the same signal level.
4. Annotations selected for permutation and shifted to semantic neighbours varying across AP sets at the same signal level.
5. The proportion of permuted annotations changing as a function of the ADS noise level.

These artificial data sets have variations within the ADS signal level that evaluation metrics should be relatively insensitive to (points 3 and 4) and variation across ADS signal levels that evaluation metric should be able to distinguish (point 5).

### Motivation for Signal and Noise Models

We have two reasons for allowing *t*_*signal*_ in the signal model to move to nearby terms in the ontology. First, real-life GO prediction tools often predict nearby terms as these tend to be correlated with *t*_*corr*_. Second, omitting these changes would favour metrics that focus on *t*_*corr*_ in the evaluation and ignore the ontology structure during evaluation, which would bias our analysis results.

The noise model, embedded in the Positive Set, aims to represent the worst type of error a classifier could make as a) *t*_*noise*_ is clearly separated from *t*_*corr*_ in the ontology and b) prediction scores for erroneous predictions, *sc*_*noise*_, come from the same distribution as correct predictions. This means that a threshold on the *sc* scores is unable to separate signal and noise predictions, ensuring that noise cannot be excluded from the evaluation. An EvM should consider our noise predictions as negative cases and score the data set accordingly. Furthermore, generating noise by permuting the GO classes from *T* keeps many other variables constant, such as average class size and average distance to the root node of predicted terms. The comparison of evaluation metrics is therefore not biased towards metrics that consider unrelated variables.

The noise that is added with lower prediction scores is motivated by trends in classifier results, where lower *sc* corresponds to lower prediction quality. Note that most metrics take this into consideration by using varying thresholds on the *sc* values. If this noise was excluded, there would be no benefit in using a threshold on *AP* set results as an EvM could simply average over all prediction results.

### Testing Evaluation Metrics with ADS

EvMs are tested by scoring the AP sets and comparing the results to the truth set, *T*. This gives us an output matrix:

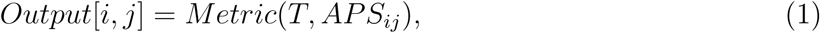

where *Metric* is a given evaluation metric, columns, *j* = [1, 2, ..*k*], correspond to different ADS signal levels, rows *i* = [1, 2..*N*] represents the different repetitions with the same ADS signal level and *APS*_*ij*_ is the *i*^*th*^ AP set obtained at signal level *j*. A good metric should have low variance within one ADS signal level (within *Output* columns) and a clear separation between the different ADS signal levels (between *Output* columns). We test this in two ways: A) visually by plotting *Output* columns as separate boxplots at different ADS noise levels (see Fig. 4) and B) numerically by calculating the rank correlation, *RC*, between *Output* matrix and ADS signal levels. Boxplots give an overview of the performance of the tested metric. *RC* monitors how closely EvM is able to predict the correct ranking of the ADS signal levels. One could also consider the linear correlation, but that would favour linear correlation over the correct rank.

**Fig 3.**
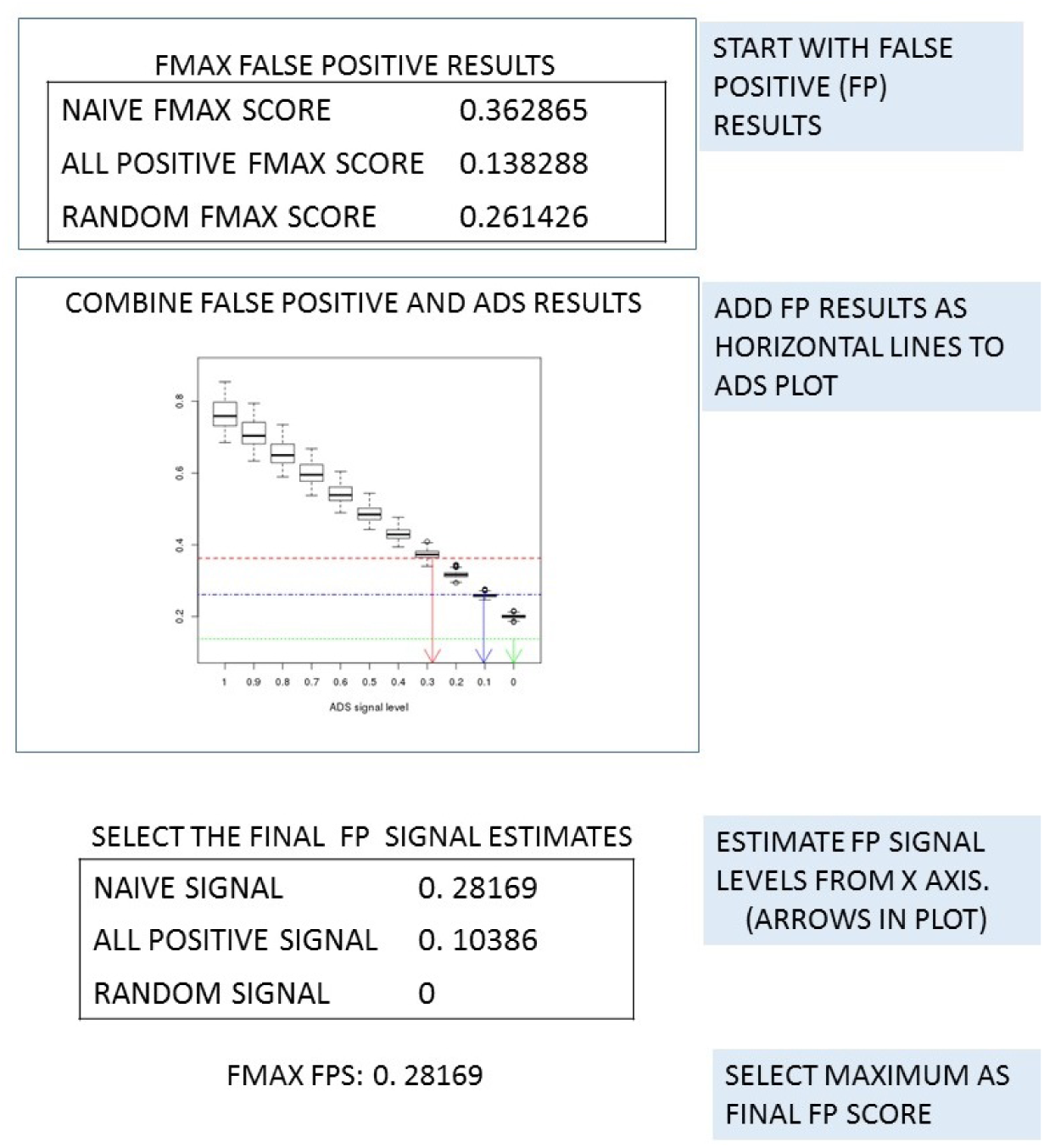
Combining FP sets with ADS results. Here *F*_*max*_ results from False Positive (FP) sets are combined with *F*_*max*_ boxplots from ADS. FP results are added as horizontal lines to the visualisation. A robust metric would score FP sets near the lowest boxplot. An FP signal estimate is obtained by comparing each FP result with medians of each signal level. This is shown in figure with vertical arrows. Resulting signals are here limited to between 0 and 1.

**Fig 4.**
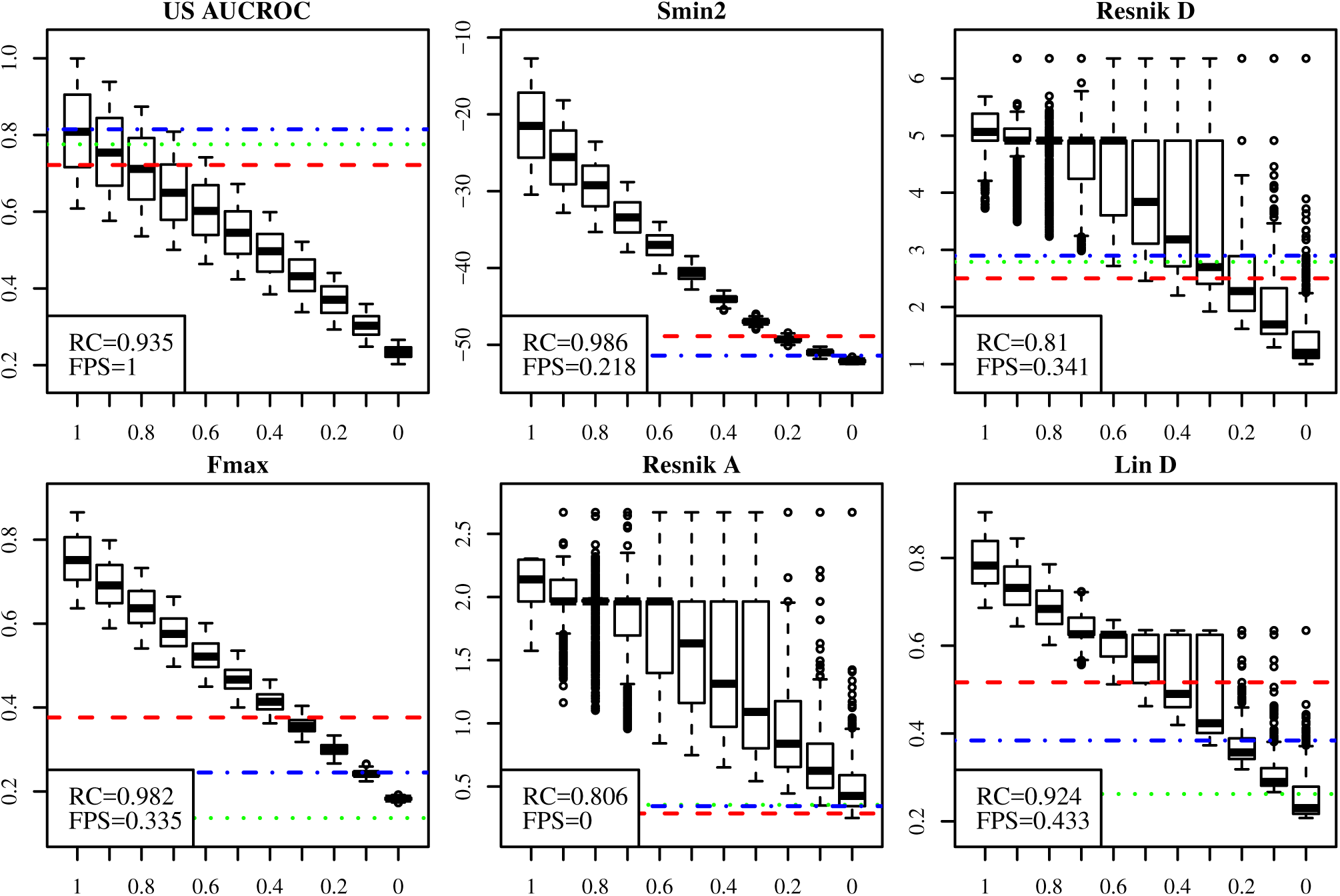
Six popular evaluation metrics compared with ADS. Visual analysis of ADS results for six evaluation metrics: US AUC-ROC, *F*_*max*_, *S*_*min*2_, Resnik score A and D, and Lin score D, obtained with CAFA data. Scores for AP sets at each ADS signal level are shown as boxplots and scores for FP sets as horizontal lines. *RC* value shows rank correlation of the AP sets with ADS signals and *FPS* shows the highest signal from FP sets. *RC* should be high and *FPS* should be low. Note the drastic differences between methods. We discuss these in the main text. We flipped the sign of *S*_*min*_ results for consistency. ADS signal is plotted on the *x*-axis and the evaluation metric, shown in headings, is plotted on the *y*-axis. Horizontal lines for FP sets are: Naive = red line, All Positive = blue line and Random = Green line

### False Positive Data Sets

In addition to AP sets, we also generated several prediction sets where all genes were annotated with the same set of non-informative GO terms or with randomly chosen GO terms. We wanted to be able to identify evaluation metrics that give high scores to prediction sets containing no signal. We refer to these sets as *False Positive Sets* (FP sets). We included the following FP sets:

1. Each gene has the same large set of the smallest GO terms. We call this the *all positive set*.
2. Each gene has the same small set of the largest GO terms, where *sc* is taken from the term frequency in the database. We call this the *naïve set*.
3. Each gene has a set of randomly selected GO terms. We call this the *random set*.

The motivation for the all positive set is to test the sensitivity of the evaluation metric to false positive predictions (low precision). Note that a set where each gene has all GO terms would be impractical to analyse. Therefore, we decided to select a large set of the smallest GO terms, each having many ancestral GO terms. The random set shows the performance when both precision and recall are very low and should therefore always have a low score. The naïve set tests if the evaluation metric can be misled by bias in the base rates to consider the results meaningful. The naïve set uses the same list of GO classes, ranked in decreasing order of size, for every analysed gene. The naïve set was originally proposed by Clark and Radivojak [21] and it has been used extensively in CAFA competitions [2]. They refer to the naïve set as naïve method or naïve predictor. Note that all these FP sets are decoupled from the annotated sequences and contain no real information about the corresponding genes.

There is a parameter that needs to be considered for FP set generation: the number of GO terms reported per gene. We selected this to be quite high (800 GO terms). This emphasises the potential inability of the metric to separate the noise signal in the FP sets from the true signal in predictions output by classifiers. We used three false positive sets:

1. Naïve-800: all genes assigned to the 800 most frequent GO terms. Here larger GO classes always have a better score.
2. Small-800: all genes assigned to the 800 least frequent GO terms. Here smaller GO classes always have a better score.
3. Random-800: all genes assigned by a random sample of 800 GO terms (sampling without replacement).

GO frequencies for FP set sampling was estimated using the UniProt GO annotation^3^.

In the visual analysis, we plot the score for each FP set as a horizontal line over the boxplot of AP set scores. A well-performing evaluation metric is expected to score all FP sets close to AP sets with minimal or no signal (*signal* ∼ 0), i.e. as a horizontal line at the lowest region of the plot (see Fig. 3).

In our numerical analysis, we summarise FP sets by **1**) Finding the approximate point where the horizontal FP set line crosses the curve created by the medians of each signal level (see Fig. 3), **2**) choosing the corresponding signal level from the *x*-axis as the FP score for the FP set in question and **3**) selecting the maximum over all FP sets as the final FP Score (referred to later as *FPS*). Note that in step **2** we limit the signal between zero and one. In step **3** we focus on the worst result over all tested FP sets. A good EvM should not be sensitive towards any of the FP sets, making the worst signal (=highest FP score) the best measure for the insensitivity towards these biases.

### Evaluation Metrics

We selected representatives from various types of EvMs commonly used in AFP evaluation (see supplementary text S2 Text). We also included modifications of some EvMs. Note that there is a large variety of EvMs available, especially in the machine learning literature [8, 10], and we cannot cover them all. The tested EvMs can be grouped into three families: **rank-based metrics, GO semantic similarity-based metrics** and **group-based metrics**.

We evaluated two rank-based metrics: area under ROC curve (AUC-ROC) and area under precision-recall curve (AUC-PR). Both strategies capture the amount of false positive and false negative errors at different threshold levels. We evaluated three semantic similarity-based metrics: Lin [22], Resnik [23] and Ancestor Jaccard (AJacc). AJacc is a novel method taking the Jaccard correlation between two sets of GO terms (i.e. the ancestor terms for both GO terms). Group-based methods compare a set of predicted GO terms to the ground truth. From this group, we evaluated SimGIC [20], Jaccard correlation (Jacc) and *S*_*min*_ [24]. *We also evaluated F*_*max*_, one of the most popular metrics in machine learning.

Most of the metrics compared here, have been used actively in AFP evaluation (see Table 1 in supplementary text S2 Text). Although AUC-PR has not been used directly, it is related to the popular precision-recall plots. All metrics except rank-based metrics can be calculated using different thresholds over prediction scores. We selected the maximum score as the final score for each metric over all possible threshold values.

The semantic similarity metrics we tested return a matrix, *M*_*g*_, of similarity values for each analysed gene, *g*. Here each *m*_*i,j*_ in *M* is a similarity score between the predicted GO term, *i*, and the correct GO term, *j*, and is calculated using the selected semantic similarity metric. Altogether *M*_*g*_ compares the set of correct GO terms for *g, S*_*corr*_, and the set of predicted GO terms for *g, S*_*pred*_, with each other. Here *S*_*corr*_ is a fixed set, whereas the *S*_*pred*_ is altered, including GO predictions for *g* that have stronger prediction scores than the acceptance threshold, *th*. We iterate through thresholds from the largest to the smallest values of the prediction scores resulting in a series of *S*_*pred*_ sets for each gene with the smallest set corresponding to the largest *th* and the largest to the smallest *th*. Next, we generate *M*_*g*_ for every *S*_*pred*_ and summarise *M*_*g*_ into a single score with a procedure described below. From threshold-wise scores we then select the best (for most metrics the maximum) score.

When semantic similarity is used as an evaluation metric, *M*_*g*_ needs to be summarised into a single score for each gene at each tested threshold. As there is no clear recommendations for this, we tested six alternatives:

**A** Mean of matrix. This is the overall similarity between all classes in *S*_*corr*_ and *S*_*pred*_

**B** Mean of column maxima. This is the average of best hits in *M*_*g*_ for classes in *S*_*corr*_.

**C** Mean of row maxima. This is the average of best hits in *M*_*g*_ for classes in *S*_*pred*_.

**D** Mean of B and C.

**E** Minimum of B and C.

**F** Mean of concatenated row and column maxima.

Methods A and D have been used previously [19]. A is considered to be weaker than D because A is affected by the heterogeneity of GO classes in *S*_*corr*_ (see Supplementary Text S2 Text and [19]). Methods B and C represent intentionally flawed methods, used here as negative controls. B is weak at monitoring false positive predictions and C is weak at monitoring false negatives. D aims to correct B and C by averaging them, but is still sensitive to outliers.

Therefore, we propose methods E and F as improvements to method D. E monitors the weaker of B and C, and is therefore more stringent at reporting good similarity. F combines two vectors used for B and C. This emphasises the longer of two vectors as the size of *S*_*pred*_ is altered. Section 5.7 in our supplementary text S2 Text motivates these summation methods in more detail. Finally, the obtained scores for each gene at each *th* threshold are averaged and the maximum over all *th* values is reported.

Most EvMs can be further modified by using the same core function with different *Data Structuring*. By data structuring we refer to the gene centric, term centric or unstructured evaluations. In **Gene Centric evaluation (GC)**, we evaluate the predicted set of GO terms against the true set separately for each gene and then summarise these values with the mean over all genes. In **Term Centric evaluation (TC)** we compare the set of genes assigned to a given GO term in predictions against the true set separately for each GO term and take the mean over all GO terms. In **UnStructured evaluation (US)** we compare predictions as a single set of gene–GO pairs, disregarding any grouping by shared genes or GO terms. In total we tested 37 metrics (summarised in S1 Table). Further discussion and metric definitions are given in S2 Text.

### Data Sets and Parameters

All metrics were tested on three annotation data sets: the evaluation set used in CAFA 1 [2], the evaluation set from the MouseFunc competition [4] and a random sample of 1000 well-annotated genes from the UniProt database ^4^. These data sets represent three alternatives for evaluating an AFP tool (see Introduction). We used annotations from the beginning of 2019 with all data sets. It should be noted that the annotation densities of the tested data sets vary over time, potentially affecting the ranking of metrics.

These data sets have different features, shown in S2 Table. MouseFunc is the largest with 45,035 direct gene–GO annotations over 1634 genes. After propagation to ancestors, this expands to 291,627 GO annotations, composed of 13,691 unique GO terms. This gives a matrix with1.42% of cells containing annotations. The CAFA data set had the smallest number of genes, with 7,092 direct gene–GO annotations over only 698 genes. After propagation, there are 61,470 GO annotations, mapping to 6,414 unique GO terms. This gives a matrix with 1.1% of cells containing annotations. The Uniprot data had the smallest number annotations, with 5,138 GO annotations over 1000 genes. These gave 46,884 annotations to 5,138 classes after propagation with 0.9% of cells in the resulting matrix containing annotations. All three data sets are intentionally small, as this is a frequent challenge in AFP evaluation. Furthermore, the number of annotations is skewed across both genes and GO terms in all data sets (see S2 Table).

We tested all metrics with three values of *k* (semantic neighbourhood size): *k* = 2, 3, 4. *RC* and *FPS* scores for these runs are listed in S3 Table, S4 Table and S5 Table. We also tested *th*_*noise*_ = 0.8 with *k* = 3. Metric scores for all AP and FP sets are available in S1 File, S2 File and S3 File.

## Results

First, we investigate the stability of results across parameter settings. Second, we show differences within groups of EvMs with the CAFA data set. We then highlight differences between data sets and identify the best performing methods overall. All results were generated using scripts available from our web page (http://ekhidna2.biocenter.helsinki.fi/ADS/).

### Stability of ADS results

ADS data generation includes two main parameters: the size of the accepted neighbourhood, *k*, while shifting to semantic neighbours and a threshold for accepting noise classes, *th*_*jacc*_. Here we show that results are stable as these values vary. We first compared the results with *k* = 2, 3, 4. We set *th*_*jacc*_ = 0.2. We created ADS and FP results with each value of *k* and calculated the Pearson and rank correlations of the results for k=2 vs. 3 and 2 vs. 4. As these were already highly correlated, it was unnecessary to proceed any further. Analysis was done separately on RC and FP results and repeated on each tested data set.

The results, shown with sub-table A in S3 Table, show that variations appear insignificant. The Pearson correlation ranges between 0.9944 and 0.9999. Similarly, rank correlation ranges between 0.9935 and 1. This suggests that results are stable against small variations in *k*. We selected results with *k* = 3 for all further analyses.

We also tested the effect of Jaccard threshold, *th*_*jacc*_, in our noise model. We compared our standard value *th*_*jacc*_ = 0.2 to *th*_*jacc*_ = 0.8. We performed a similar evaluation as with values of *k* (see sub-table B in S3 Table). The minimum and maximum Pearson correlation was 0.9991 and 0.9999, respectively, and the minimum and maximum rank correlation was 0.9945 and 0.9997, suggesting that our results are stable against *th*_*jacc*_

### ADS Identifies Differences Between Evaluation Metrics

We compared the following evaluation metrics: US AUC-ROC, *F*_*max*_, *S*_*min*_ (referred as *S*_*min*2_ in figure), Resnik with summary method D and A, and Lin with summary method D to see if ADS can find performance differences. Visualisations for these metrics are shown in Fig 4 (for exact values see S6 Table).

Results with CAFA data show drastic differences between evaluation metrics. We see a clear separation across ADS signal levels in the boxplots for *F*_*max*_ (*RC* = 0.982) and *S*_*min*_ (*RC* = 0.986), and next best separation is for US AUC-ROC. Three of the semantic similarity measures performed poorly: they have unstable distributions in the obtained RC scores. Resnik in particular, is shown to have the worst separation (*RC* = 0.806 and 0.81 for summary A and D, respectively). Our results show that it is impossible to infer the original signal levels using these particular semantic similarity measures.

It could be argued that comparing random data and data with a strong signal should be sufficient for metric evaluation. Our results, however, show that in most plots the first (*signal* = 1) and last (*signal* = 0) boxplots do not fully reveal how good the evaluation metric is. For Lin score and Resnik score, for example, two extreme boxplots might be acceptable, assuming clear separation. However, the intermediate levels reveal considerable variability.

With FP sets visualised with horizontal lines, we see different methods failing. Now AUC-ROC has the worst performance as it ranks the FP sets as equally good as *signal* = 1. Note, that FP sets do not convey any real annotation information, but represent biases related to the GO structure. Altogether, only Resnik with method A has excellent performance with the FP signal and only *S*_*min*_ has good performance in both tests. These results demonstrate that neither ADS nor FP sets are sufficient to expose all weaknesses in isolation. They clearly show orthogonal views on the evaluated methods, both of which are important.

### ADS Confirms Known Flaws in AUC-ROC

Here we focus on the performance of AUC methods, using area under receiver operating characteristic curve (AUC-ROC) and area under precision-recall curve (AUC-PR) with the CAFA data set. AUC-ROC has been used extensively in AFP evaluation, whereas AUC-PR has not. We look at the unstructured (US), gene centric (GC) and term centric (TC) versions of both AUC methods.

AUC-ROC has been criticised for being a noisy evaluation metric with sensitivity to sample size and class imbalance [8, 25]. ADS demonstrates that AUC-ROC metrics correlate reasonably well with signal level, but fail with FP sets (Fig 5 and S6 Table). These results are expected, as FP sets monitor sensitivity to class imbalance. Surprisingly, two of our of three AUC-ROC methods rank the FP sets as good as *signal* = 1 (Fig 5, top row). This means that near-perfect prediction results could not be distinguished from false positives using this metric.

**Fig 5.**
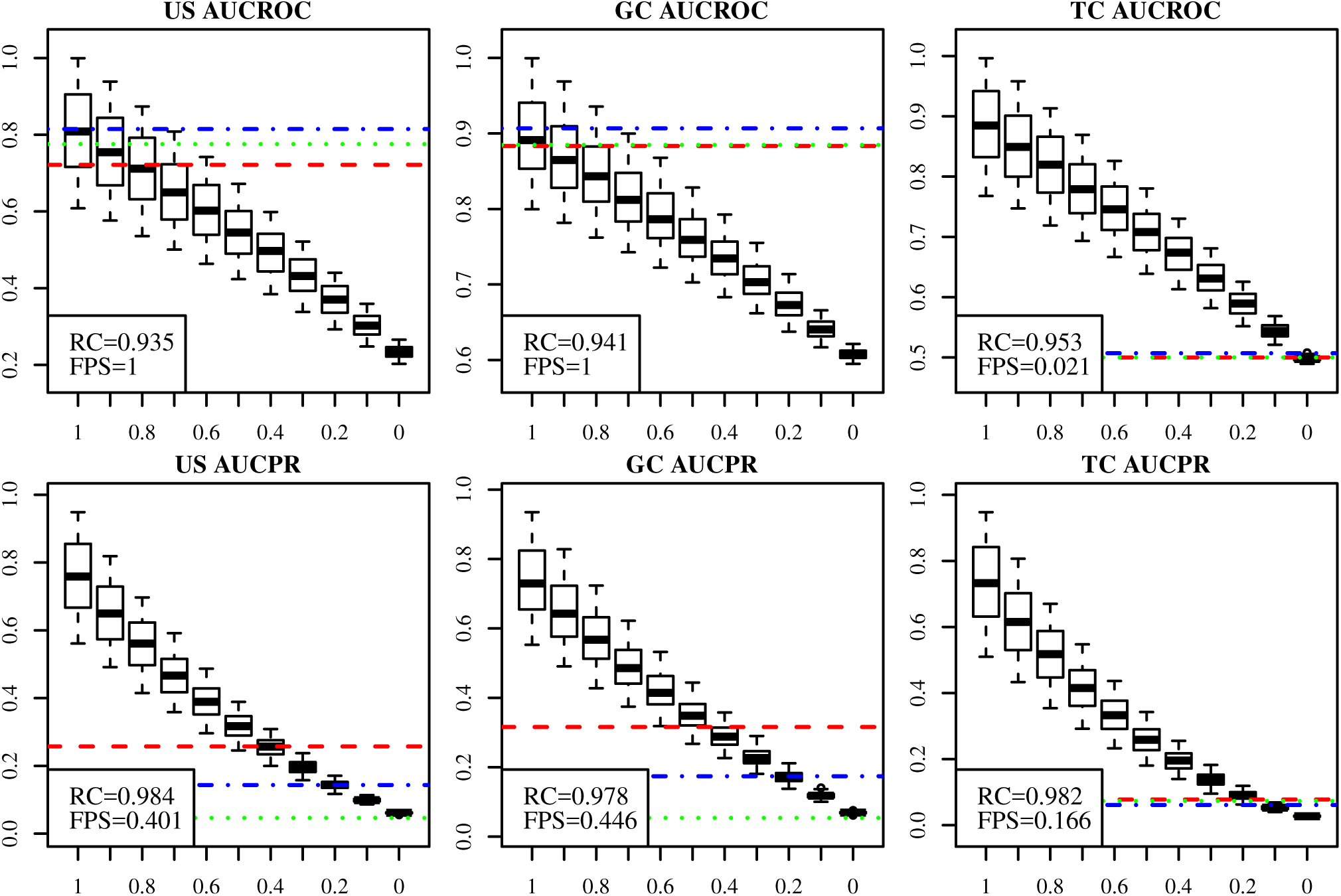
AUC metrics compared using ADS. We compared standard AUC (AUC-ROC) and AUC for precision-recall curve (AUC-PR) with CAFA dataset. We test unstructured (US), gene centric (GC) and term centric (TC) versions. US AUC-ROC and GC AUC-ROC exhibited exceptionally poor performance with the FP sets. TC AUC-PR showed good performance with both ADS and FP sets. *RC* and *FPS* are explained in previous figure text. ADS signal is plotted on the *x*-axis and the evaluation metric, shown in headings, is plotted on the *y*-axis.

The poor FP set performance of some AUC metrics is caused by the identical treatment of small and large GO classes in US and GC analysis. However, a positive result for a large class is more probable from a random data set. This is worsened by the extreme class size differences in GO. This problem was corrected in the MouseFunc competition [4] by dividing GO classes to subsets based on class sizes and in CAFA 2 [3] by analysing each GO term separately, as in TC AUC-ROC. Indeed, the TC analysis displayed in the last column performs well with the FP sets. This is replicated across all data sets.

When AUC-ROC and AUC-PR are compared, while we see that FP sets have a weaker signal with US AUC-PR and GC AUC-PR, the problem persists. In the correlation scores, however, AUC-PR metrics consistently outperform AUC-ROC metrics. The difference between AUC-ROC and AUC-PR might be explained by the definition of AUC-ROC (see supplementary S2 Text). Since TC AUC-PR assigns low scores to FP sets and achieves high correlation, it is our recommended method with TC AUC-ROC in second place. Furthermore, these results argue strongly against the usage of US and GC versions of AUC metrics.

### ADS Reveals Drastic Differences Between Semantic Summation Methods

We compared the following GO semantic similarities: Resnik, Lin and Ancestor Jaccard (AJacc), each coupled with six different semantic summation methods, A-F (see Methods). The results are shown in Fig. 6 and S6 Table. Fig. 6 is organised so that the columns contain the compared semantic similarities and rows contain the summation methods. Ideally, we want a combination that delivers good separation in boxplots and ranks FP sets next to AP sets with *signal* = 0.

**Fig 6.**
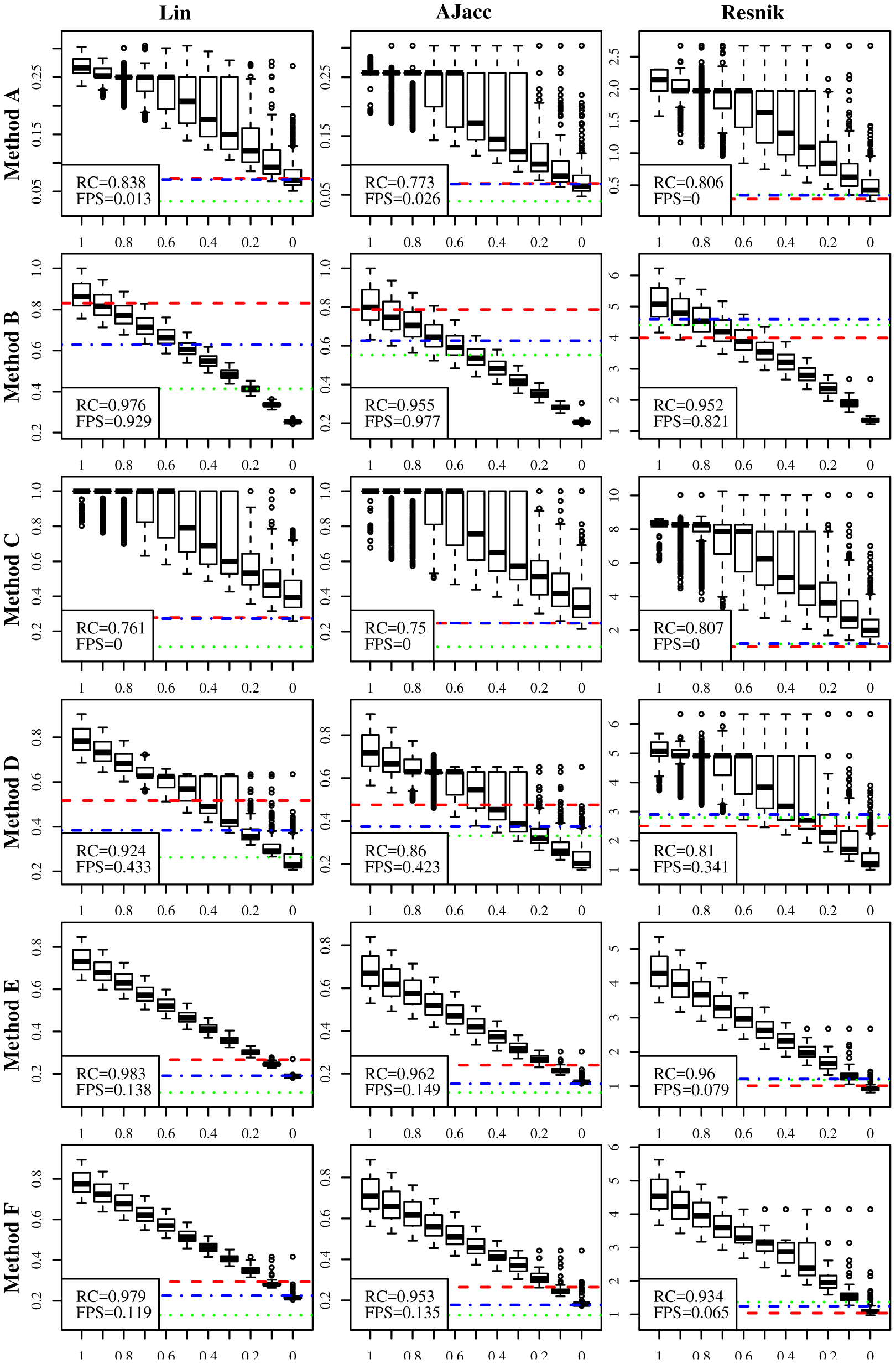
Semantic similarities with different summation methods. We present every combination of three semantic similarity-based methods (Resnik, Lin, AJacc) in columns and six semantic summation methods (A-F, see Methods) in rows. We show again the results for CAFA dataset. The summation methods have a bigger impact on performance than the actual metric. The novel summation methods, E and F, outperform the previous standards, A and D. ADS signal is plotted on the *x*-axis and the evaluation metric combined with a given summation method is plotted on the *y*-axis.

This comparison shows the power of ADS. Our first observation is that the choice of summation method has a bigger impact than which semantic measure is used. Methods A and C fail at separating signal levels in ADS and methods B and D fail at ranking FP sets next to the *signal* = 0. The only methods that perform well in both ADS and FP tests are E and F. Furthermore, the intentionally flawed methods, B and C, were found to be weak by either ADS or FP tests.

Our second observation is that Lin similarities exhibit marginally better performance than the other metrics tested, as shown in RC values. While these results might suggest that Lin similarity summarised with methods E or F performs the best, this is dependent on data set (see below).

### S_min_ and SimGIC have Similar Good Performance

The third group of metrics we tested were of methods that score sets of predicted GO terms. Some methods, like SimGIC and SimUI [19], calculate the Jaccard correlation between predictions and the ground truth, whereas *S*_*min*_ calculates a distance between them [26]. Most of these metrics include Information Content (IC) weights [19] in their function. We have two alternatives for IC, but limit the results here to IC weights proposed in the *S*_*min*_ article [26] (details on two versions of information content, *ic* and *ic*2, are discussed in the S2 Text). We still show results for both *ic* and *ic*2 in later analyses. We further include gene centric and unstructured versions of the following functions:

**SimGIC** Gene centric original version (weighted Jaccard)

**SimGIC2** Unstructured modified version of SimGIC

**S**_**min1**_ *S*_*min*_ original version

**S**_**min2**_ Gene centric version of *S*_*min*_

**GC Jacc** Gene centric unweighted Jaccard (original SimUI)

**US Jacc** Unstructured version of SimUI (unstructured Jaccard)

The results are shown in Fig. 7 and S6 Table. Both *S*_*min*_ and SimGIC show very high correlation with the ADS signal level. However, in FP tests, SimGIC has better performance. Also, US Jacc shows quite good performance, but GC Jacc is weaker. Introducing gene centric terms in *S*_*min*2_ resulted in a notable increase in FP scores in all three data sets. SimGIC2, our unstructured version of SimGIC, showed a minor improvement in correlation with all three data sets. Overall, the unstructured versions (*S*_*min*1_, SimGIC2 and US Jacc) outperformed the gene centric versions (*S*_*min*2_, SimGIC, GC Jacc) in correlation values, FP tests or both. Performance of *S*_*min*_ and SimGIC metrics across different data sets was surprisingly similar. Our recommendation is to use *S*_*min*1_ or SimGIC2. Alternatively, one can use US Jacc, despite weaker performance than *S*_*min*_ and SimGIC2, when a simpler function is required.

**Fig 7.**
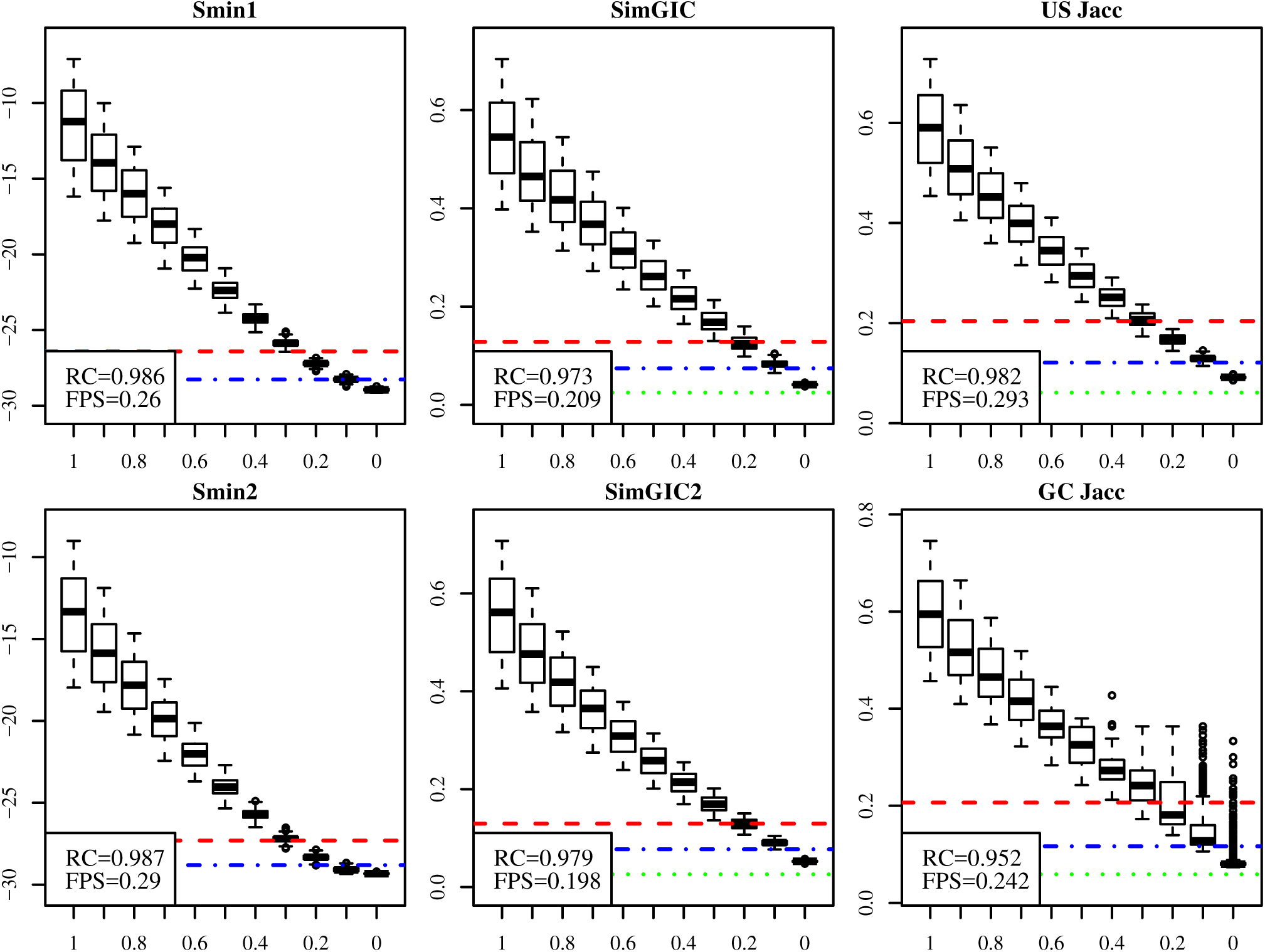
S_min_ and SimGIC metrics. We compared two versions of *S*_*min*_, SimGIC and Jaccard each with Uniprot data. Performance was similar with all metrics, with the exception of GC Jacc which had a weaker correlation. *S*_*min*1_ and SimGIC2 are slightly better methods. ADS signal is plotted on the *x*-axis and the evaluation metric, shown in headings, is plotted on the *y*-axis. We flipped the sign of the *S*_*min*_ metrics for consistency.

### Metric Performance Varies Between Data Sets

We wanted to compare all the discussed evaluation metrics to see which of them performed best overall. We plotted *RC* against *FPS* for all metrics tested on the CAFA data set (see Fig. 8). High quality metrics appear in the upper left of the plot. Metrics that fail with FP tests will be on the right of the plot. Metrics failing in ADS correlation will be lower in the plot.

**Fig 8.**
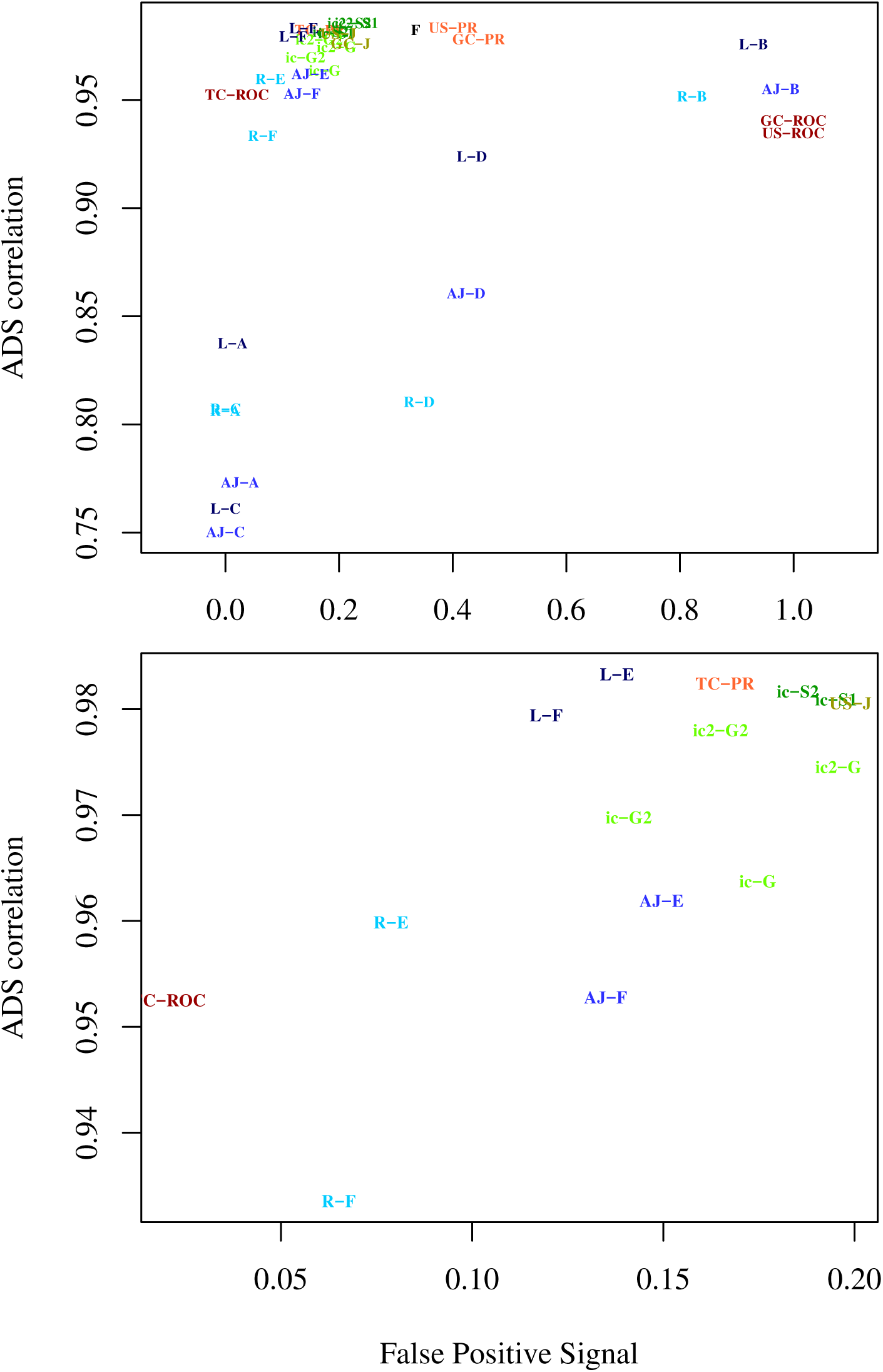
Top metrics for CAFA data set. Metrics are shown with different colours. *F*_*max*_ is black, AUC-ROC and AUC-PR are in red, Jacc, *S*_*min*_ and SimGIC are in green and semantic similarity metrics are in blue. Colours are varied between different metrics in the same group. The figure uses the following abbreviations: AJ = AJacc, F = *F*_*max*_, G = SimGIC, J = Jacc, L = Lin, PR = AUC-PR, R = Resnik, ROC = AUC-ROC and Sm = *S*_*min*_. Abbreviations are combined with summation or data structuring method. For example, Resnik F is R-F, Lin A is L-A, TC AUCROC is TC-ROC, ic2 SimGIC is ic2-G and ic *S*_*min*_ is ic-S. High quality metrics appear in the upper left corner (high correlation and low False Positive Signal). This clustered part is zoomed in the lower panel for clarity.

The upper panel in Fig. 8 shows big differences in performance. The semantic similarity metrics (shown in blue) show drastic scatter on the *y*-axis and AUC metrics (red) show drastic scatter on the *x*-axis, in agreement with results from earlier sections. Clearly, usage of unstructured and gene centric Area Under Curve methods (GC AUC-ROC, US AUC-ROC, US AUC-PR, GC AUC-PR) and semantic similarities based on signal summation methods A, B, C and D cannot be recommended. The metrics with the best performance (lower panel in Fig. 8) are Lin E, Lin F, TC AUCPR, two versions of SimGIC2 and *S*_*min*2_.

When we repeated the same analysis with the other two data sets, the rankings are different. We show only the best-scoring metrics in Fig. 9 for MouseFunc and Uniprot. The performance of all metrics is shown in S1 Figure and S2 Figure. In MouseFunc, the best performing group is the SimGIC - *S*_*min*_ methods with *S*_*min*2_ as the best method. Lin E and Lin F are also among the top performers. TC AUC-PR, one of previous best performers, has weaker performance. *S*_*min*_ improved its performance, while TC AUC-PR got weaker. The Uniprot data set had differing results as well, with TC AUC-PR, Resnik E, Resnik F and TC AUC-ROC showing the best performance. The biggest drop among the top performing metrics is with Lin semantics (Lin E and Lin F).

**Fig 9.**
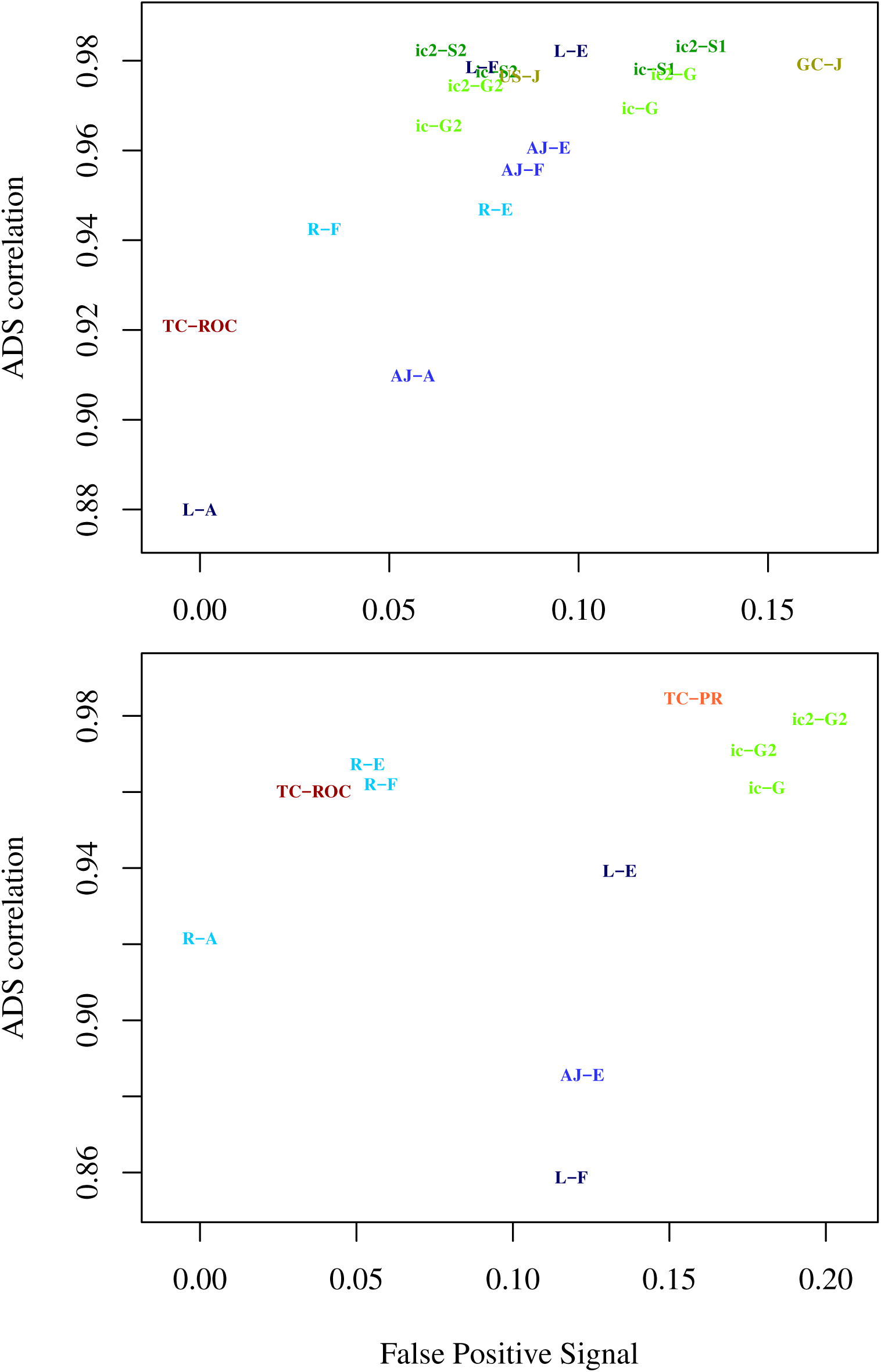
Top metrics for MouseFunc (top panel) and Uniprot (lower panel) data sets. Here we repeat the lower plot from Fig. 8 on the other two data sets. The performance of individual EvMs varies between data sets. Colouring and labels are explained in Fig. 7.

Semantic similarities had surprising behaviour in these experiments. Lin similarity (with methods E and F) was clearly the best performing semantic similarity-based metric in CAFA and MouseFunc, with Resnik as the weakest. However, on Uniprot data, Resnik improves its performance, making it the best semantic similarity method, while Lin’s performance drops significantly. We are unable to explain these differences.

Another surprising finding is that we see higher FP signals for TC AUC-PR than expected, particularly in the MouseFunc data. As term centric analysis evaluates each GO class separately, it should be insensitive to biases created by favouring larger or smaller GO classes. We will explain this in the next section.

Note that these data sets differed from each other in size and density, with Uniprot being the smallest and sparsest data set, CAFA being intermediate and MouseFunc data being the largest and densest data set. We see the following tendencies in the results: a) Lin semantic performs well until the size and sparsity of the Uniprot data is reached, b) term centric AUC-ROC and AUC-PR perform well except with denser and larger mouse data and c) simGIC methods appear to perform consistently. Elucidating the exact reasons for these performance differences would require further studies. Altogether, this suggests that selecting the best evaluation metric is data-dependent, showing the necessity of ADS-style tests.

### Selecting Overall Best Evaluation Metrics

Results above showed variation in the ordering of metrics across the datasets. Still, it is important to see if we can select for AFP method comparisons metrics with best overall performance. Therefore, we collected the results of top performing metrics for each data set into Table 1, showing RC (Rank Correlation) and FPS (False Positive Set) values. In addition, we show the same results for popular methods with weaker performance. Again, a good metric should have high RC values and low FP values. Therefore, we show in RC tests the five highest scores in each column with bold and five lowest scores in underlined italics. In FP tests we similarly show five lowest scores in bold and five highest scores in underlined italics. Excellent metrics should have bold numbers and no italics.

Furthermore we wanted to highlight the consistent high performance of metrics by setting a threshold on *RC, t*_*RC*_, and threshold on *FPS, t*_*F P S*_, and requiring that *RC > t*_*RC*_ and *FPS < t*_*F P S*_. We highlight the failures of these tests in Table 1 in red. For RC values, we observed that the top performing metrics would be selected with threshold *t*_*RC*_ = [0.94, 0.98] and therefore set *t*_*RC*_ = 0.95. For FP values, we found that the top performing metrics would be selected with *t*_*F P S*_ = [0.10, 0.25] and set *t*_*F P S*_ = 0.16. There is a trade off when setting these thresholds as different values can emphasise either RC or FPS. However, both tests should be monitored for metric selection. In conclusion, a good metric should have as little red shading as possible and high performance in both RC and FP tests. Our supplementary table S6 Table shows these results for all the tested metrics. We selected the best performing metrics from supplementary table to the upper portion of Table 1.

#### Top Performing Metrics

The upper portion of Table 1 shows metrics with encouraging performance. We first identify metrics that show good performance in a majority of tests. TC AUC-PR shows good performance with the highest ranking RC values. Its FP values are around 1.5, except for MouseFunc data, where its FP result is ∼0.3. Therefore, we mark it only as a potential recommended method. TC AUC-ROC shows very good performance in FP results. However, its RC results are weaker and its RC result from mouse data is far lower. Still, it is a potential recommendation. Lin score with method E shows the most consistent high performance among the Lin and AJacc semantics in the upper block. Differences are seen in RC values. However, Lin E fails in RC test on the Uniprot data set. Therefore, we mark it also as a potential recommendation. Resnik semantic shows different behaviour than Lin and AJacc in these results. Still, it also fails the RC test with MouseFunc data. All these recommended metrics showed good performance in all, but one test.

Finally, ic SimGIC and ic SimGIC2 shows quite good performance across the tests, with ic SimGIC2 often outperforming ic SimGIC. We mark also ic2 SimGIC2 as a potential recommended method, although it fails in FP test with UniProt data. Note that ic2 SimGIC2 goes only slightly over the selected threshold (0.18) and in other data sets it stays below *t*_*F P S*_. However, ic SimGIC2 mostly outperforms other tested SimGIC functions. Overall, based on these results, ic SimGIC2 is our strongest overall recommendation.

We are surprised to see that the ic2 *S*_*min*1_ [24] has an elevated FP signal, despite using *ic* weights to emphasise small GO classes. However, we still point out that FP results for *S*_*min*_ are not bad, except for Uniprot data and has the strongest RC results in the table. Indeed, *S*_*min*_ was one of the best performing metrics for MouseFunc data. Therefore, we included it in our potential recommendations.

We were also surprised by FP signals with TC AUC-PR, as term centric analysis should be insensitive to FP signal. This is explained by our calculation method for AUC-PR. Even when there is no variance among prediction scores across all the genes, we allow precision-recall curve to return a single point when calculating the area. While this is in agreement with how to calculate AUC-PR [27], it can generate a non-zero result that can bias our analysis in some data sets. This can be corrected by setting AUC-PR = 0 when the range of prediction scores across all the analysed data points is zero. Note that predictions in this case do not contain any information, making the AUC-PR = 0 reasonable. So here our analysis proposes simple improvement to TC AUC-PR that would potentially make it a potential top performing metric.

In conclusion, our results suggest that ic SimGIC2 as the most stable, well-performing metric, despite being beaten by other potential recommendations (TC AUC-PR, TC AUC-ROC, Lin E, Resnik E and *S*_*min*_) with some data sets.

#### Issues with Some Widely-used Metrics

Metrics in the lower part of Table 1 show poor performance. All these metrics have been used in AFP research, and some of these metrics have been widely-used. The popular *F*_*max*_ and US AUC-PR show high FP values. Although *F*_*max*_ is preferred for its simplicity, our results show that even the simple US Jacc performs better, as it shows similar RC values but has roughly 30% lower FP scores (values: 0.273, 0.184, 0.073 shown in S6 Table). Furthermore, the weighted Jaccard metrics (SimGIC and SimGIC2) correct FP error even further. US AUC-PR shows even worse FP results. Still, TC AUC-PR would correct this problem while maintaining almost the same level of RC values. Similar results are seen with AUC-ROC methods. GC and US versions of AUC-ROC should be avoided, as they are consistently the worst performing methods in our FP tests. Again, the TC AUC-ROC corrects this bias. Finally, the results for semantic similarities in the lower block are very weak, often with the weakest RC scores in the table.

## Discussion

GO classifiers and other methods for automated annotation of novel sequences play an important role in the biosciences. It is therefore important to assess the quality of the predictions that they make. Evaluation depends heavily on the metrics used as they define how methods will be ultimately ranked. However, little research has been done on the evaluation metrics themselves, despite their impact in shaping the field.

Artificial Dilution Series (ADS) is a novel method to systematically assess how reliably different evaluation metrics cope with assessing GO classifiers. ADS is a valuable new tool for developing and complementing the existing evaluation methodology. Using ADS we have revealed drastic differences between popular evaluation metrics and metrics that give inflated scores to false positive annotations.

We demonstrate ADS by testing a large number of evaluation metrics including many variations. We were able to show that:

- One should only use term centric AUC-ROC and AUC-PR methods and avoid their gene centric and unstructured versions.
- Our novel summary methods, E and F, both outperform existing semantic similarity summary methods.
- The simGIC metric shows consistently high performance across all data sets.

Other well performing metrics, besides simGIC2, were TC AUC-PR, TC AUC-ROC, Lin score with method E and Resnik score with method E. These even outperformed ic simGIC2 on some data sets, which suggests that the selection of the best evaluation metric is data-dependent. Note that our analysis scripts and ADS are freely available, allowing testing with any GO data set.

One motivation for the ADS project is the development of better evaluation metrics for GO classifier comparisons. We tested, for example, the following variations on existing metrics:

1. We modified the summary methods for semantic similarities.
2. We tested Area Under Precision Recall curve (AUC-PR).
3. We tested the unstructured variant of the SimGIC function.

Here 2, 3 and some of the methods tested in 1 improved performance. All of these modifications were simple to implement using our existing evaluation code. Furthermore, we have distributed a standalone C++ program that calculates all of the evaluation metrics tested here, allowing them to be used in practice.

Our results included metrics that were expected to exhibit poor performance. These were included as negative controls in our analysis to prove that ADS works as expected. Indeed, we recognised that US AUC-ROC and GC AUC-ROC show very bad performance with FP tests. This is in agreement with the results by Ferri et al. [8] with AUC-ROC and its weakness with uneven class sizes. As expected, semantic summation methods B and C had poor performance with FP tests and ADS RC values, respectively (see supplementary S2 Text).

The importance of a good evaluation metric is best demonstrated with flawed metrics. A metric with sensitivity to our naïve FP set, *EvM*_*naive*_, would allow “cheating” in method comparison. For example, when the classifier lacks predictions for a gene, *G*, it could improve its results simply by returning a set of naïve predictions for *G* (i.e. the base rate for each GO class). A better alternative for the end user would be to not make any predictions. Also, a bad *EvM*_*naive*_ would evaluate the correct GO term as an equally good prediction as any of its parents. This would promote predicting large unspecific GO terms for all genes, although the small terms contain more biological information. Likewise, a metric penalising false positives weakly would promote methods that predict as many terms as possible and a metric penalising false negatives weakly would promote predicting as few GO terms as possible. Note that these artefacts are often hard to observe without the kinds of comparisons we have presented.

Our results are dependent on the used signal and noise model, as these are used to create the correct and wrong predictions. Although these models were motivated in the Methods section, we emphasise that this is only the first version of ADS. Our plan is to extend these models and use them in addition to our current models. ADS analysis can be easily extended in this manner. We propose the ADS approach in general, without limiting it to specific signal and noise models.

Furthermore, we use a real dataset as a starting point, thus maintaining its challenging features. Also, there are no such real classifier result sets available, to our knowledge, where the correct ordering of the classifiers is known. So one is bound to use some kind of simulation in this type of analysis. Here the controlled percentage of noise makes the correct ordering of the results known. We point out that the only similar evaluation, extensive work by Ferri et al. [8], uses fully artificial datasets to compare evaluation metrics. Supplementary text 2 S2 Text compares our work in more detail to this and other previous research.

We envision several application scenarios for the ADS. The first, is a general user, who would like an overall answer to the question: which metrics are optimal? For this case a set of metrics that have stably high performance on very different datasets is needed. In this work we give our recommendations with tests on three data sets. Further testing on a larger spectrum of datasets and with altered noise models is planned for the follow up articles. The second scenario is a specialised user who wants to optimise metrics for the type of data he or she is testing. Here the researcher would perform the ADS analysis using the selected dataset(s). This ensures the applicability of the selected group of evaluation metrics on the used dataset(s). Finally, we expect that ADS would be useful for the developers of evaluation metrics, as it allows thorough testing of novel metrics.

ADS is designed to compare evaluation metrics that are used in GO term prediction for proteins, however, similar problems with evaluation metrics occur in other areas of the biosciences. Complex hierarchical classifications are used, for example, for classifying disease genes. Also the use of ontologies in the biosciences is growing [28], creating similar classification challenges. Furthermore, similar hierarchical structures are seen in other fields (e.g. in WordNet [29]) and evaluation metrics are shown to fail even without hierarchical structures, when there is a class imbalance problem [17].

The motivation for the ADS project came from our own GO classifier work [6, 7], where we noted that standard classifier metrics gave misleading results. Indeed, we expect that the method development community and researchers doing comparisons between GO prediction methods will benefit most from this work. However, the benefits do not end there. Currently it can be difficult for readers and reviewers to estimate the importance of new predictive bioinformatics methods as different articles use different metrics and test data sets. Indeed, the reliability of scientific findings [30, 31] and over-optimistic results [18, 32] have generated much discussion recently. We hope that ADS will improve this situation by developing robust and transparent standards to select evaluation metrics in future bioinformatics publications.

## Conclusion

Science needs numerous predictive methods, making the comparison and evaluation of these methods an important challenge. The central component of these comparisons is the Evaluation Metric. In the case of Automated Function Predictors the selection of the evaluation metric is a challenging task, potentially favouring poorly performing methods. In this article we have described Artificial Dilution Series (ADS) which is, to our knowledge, the first method for testing classifier evaluation metrics using real data sets and controlled levels of embedded signal. ADS allows for simple testing of evaluation metrics to see how easily they can separate different signal levels. Furthermore, we test with different False Positive Sets representing data sets that the evaluation metric might mistakenly consider meaningful. Our results show clear performance differences between evaluation metrics. We also tested several modifications to existing evaluation metrics, some of which improved their performance. This work provides a platform for selecting evaluation metrics for specific Gene Ontology data sets and will simplify the development of novel evaluation metrics. The same principles could be applied to other problem domains, inside and outside of the biosciences.

## Acknowledgements

IP was supported by Emil Aaltonen Foundation and University of Helsinki. PT was supported by Helsinki Institute of Life Science (HiLife). We would like to thank Rishi Das Roy and Alan Medlar for valuable comments and fruitful discussions. We also acknowledge Alan Medlar for reviewing our text.

## Supporting Information

**S1 Figure Scatter plot visualization of EvM performance** This scatter plot represents EvM performances on MouseFunc data set. Labels and abbreviations, used here, are explained in fig 8.

**S2 Figure Scatter plot visualization of EvM performance** This scatter plot represents EvM performances on Uniprot data set. Labels and abbreviations, used here, are explained in fig 8.

**S1 Text ADS pseudocode** Pseudocode for generation of ADS.

**S2 Text Supplementary text** Text shows an example on how the signal and noise is defined in ADS. Text also shows labels, definitions and discussion of tested metrics. Finally, we discuss the summary methods for semantic similarities and motivate our novel summary methods.

**S1 File Metric scores** Metric scores for all generated AP and FP sets (*k* = 2).

**S2 File Metric scores** Metric scores for all generated AP and FP sets (*k* = 3).

**S3 File Metric scores** Metric scores for all generated AP and FP sets (*k* = 4).

**S1 Table Abbreviations and features for all compared EvMs** We represent a summary table of all the compared EvMs. Table shows the used abbreviation, core function, used data summary method, used threshold function over the classifier prediction score, used IC weighting and summary method for semantic similarities. In addition we mark the EvMs that have been popular in AFP evaluation, ones that have some novelty and ones that we consider to be simple. We also mark EvMs that we expect to perform badly as negative controls. More detailed description of these EvMs is in suppl. text S2 Text.

**S2 Table Stability of the results when parameters are varied** Here we evaluate the stability of the results when we vary parameter *k* and *th*_*jacc*_. Table A shows results for different *k* values and table B shows results for two values of *th*_*jacc*_. We selected in each comparison one data set, created results for all EvMs using two parameter values and compared the obtained results using Pearson and Spearmann rank correlation. FPS and RC results are evaluated here separately. Tested parameters show very little effect on the results.

**S3 Table Features of the three used data sets** Table explains the size and density of the three used data sets (CAFA1, MouseFunc, UniProt). Data sets are explained in main text. Table shows skewed distribution of GO terms per gene by representing selected quantiles from this distribution. This is shown separately for each data set with and without the propagation to parent terms. We also show the same size distribution for the GO terms, again showing the skewness of the size distribution.

**S4 Table ADS results** *RC* and *FPS* scores for tested metrics using *k* = 2. Table is tab-delimited text file.

**S5 Table ADS results** *RC* and *FPS* scores for tested metrics using *k* = 4. Table is tab-delimited text file.

**S6 Table ADS results for** *k* = 3 **with highlighting** *RC* and *FPS* scores for tested metrics using *k* = 3. Table is an excel file that shows similar results as table 1 but for all the metrics. Colouring highlights cases where metric fails a test. Thresholds can be adjusted in this table to change the highlighting.

## Footnotes

1 http://ekhidna2.biocenter.helsinki.fi/ADS/

2 https://bitbucket.org/plyusnin/ads/

3 (ftp://ftp.ebi.ac.uk/pub/databases/GO/goa/UniProt/, 03.2016)

4 ftp://ftp.ebi.ac.uk/pub/databases/GO/goa/UniProt/, 01.2019

## Notes

#### Summary of Updates

Datasets used for evaluation updated to 01.2019.

http://ekhidna2.biocenter.helsinki.fi/ADS/

https://bitbucket.org/plyusnin/ads/

