## Supplementary figures and images for "Novel Comparison of Evaluation Metrics for Gene Ontology Classifiers Reveals Drastic Performance Differences"

### Supplementary Figure 1

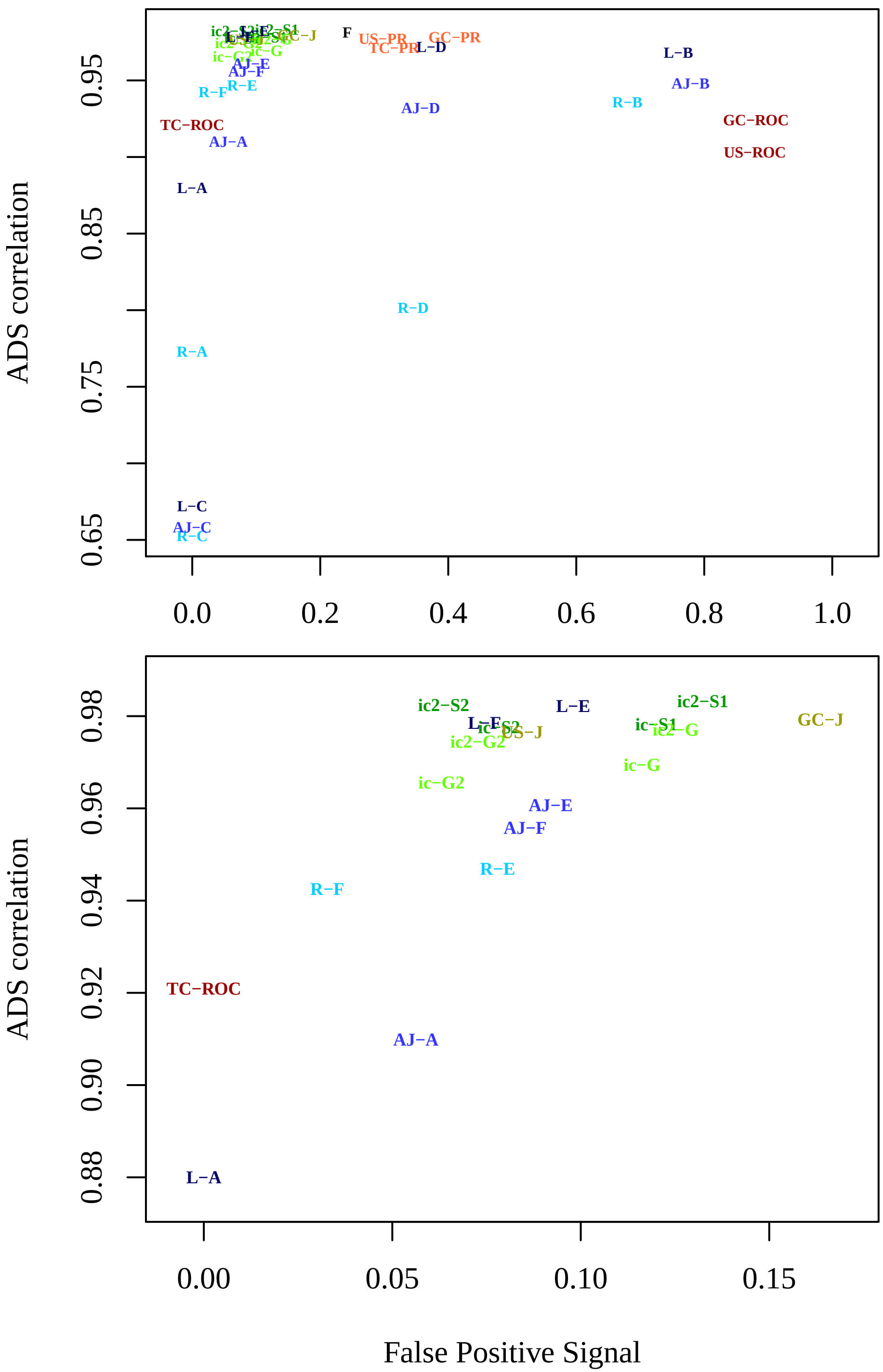

### Supplementary Figure 2

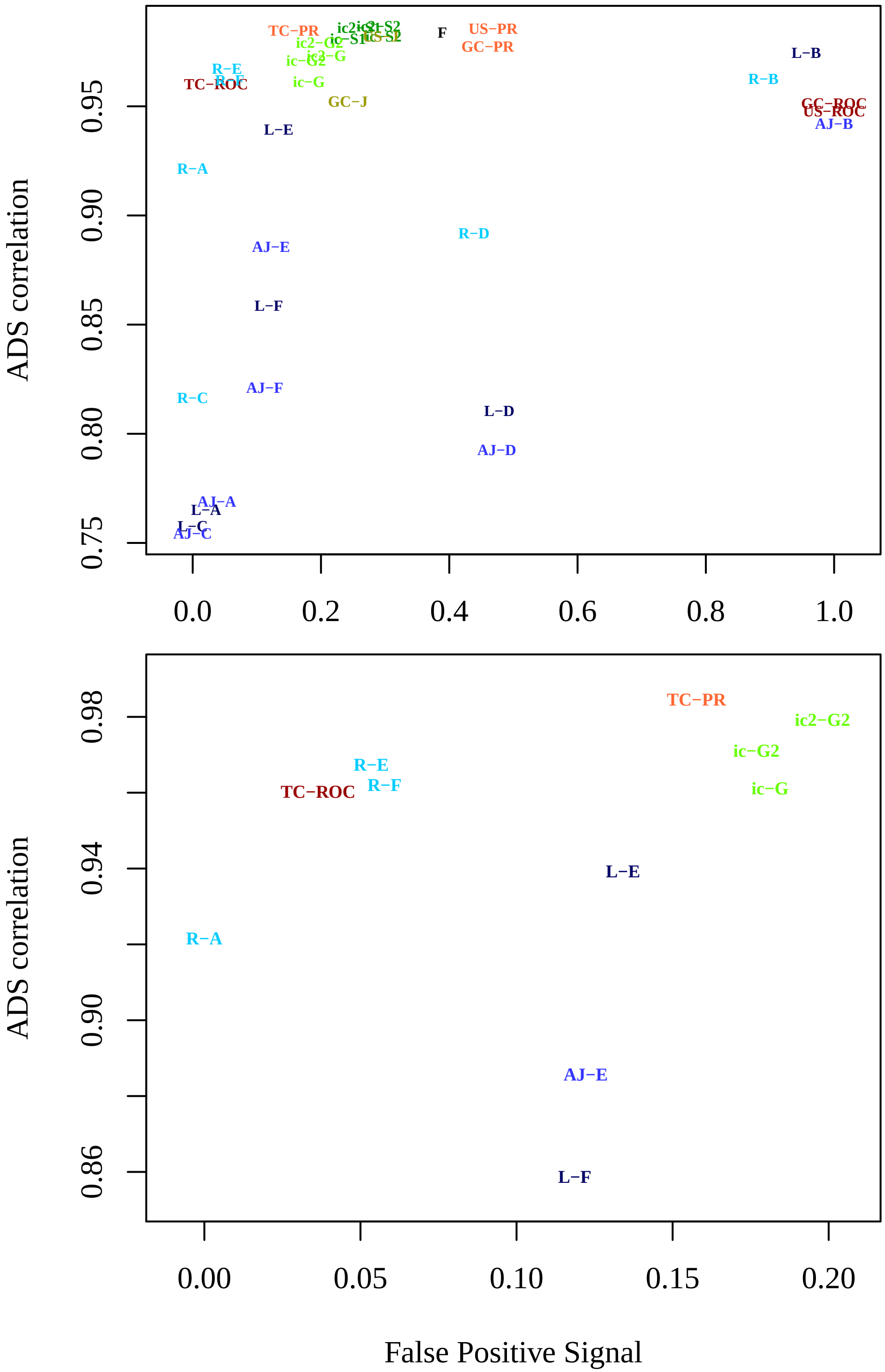

### uniprot.1000_boxplot1.jpeg

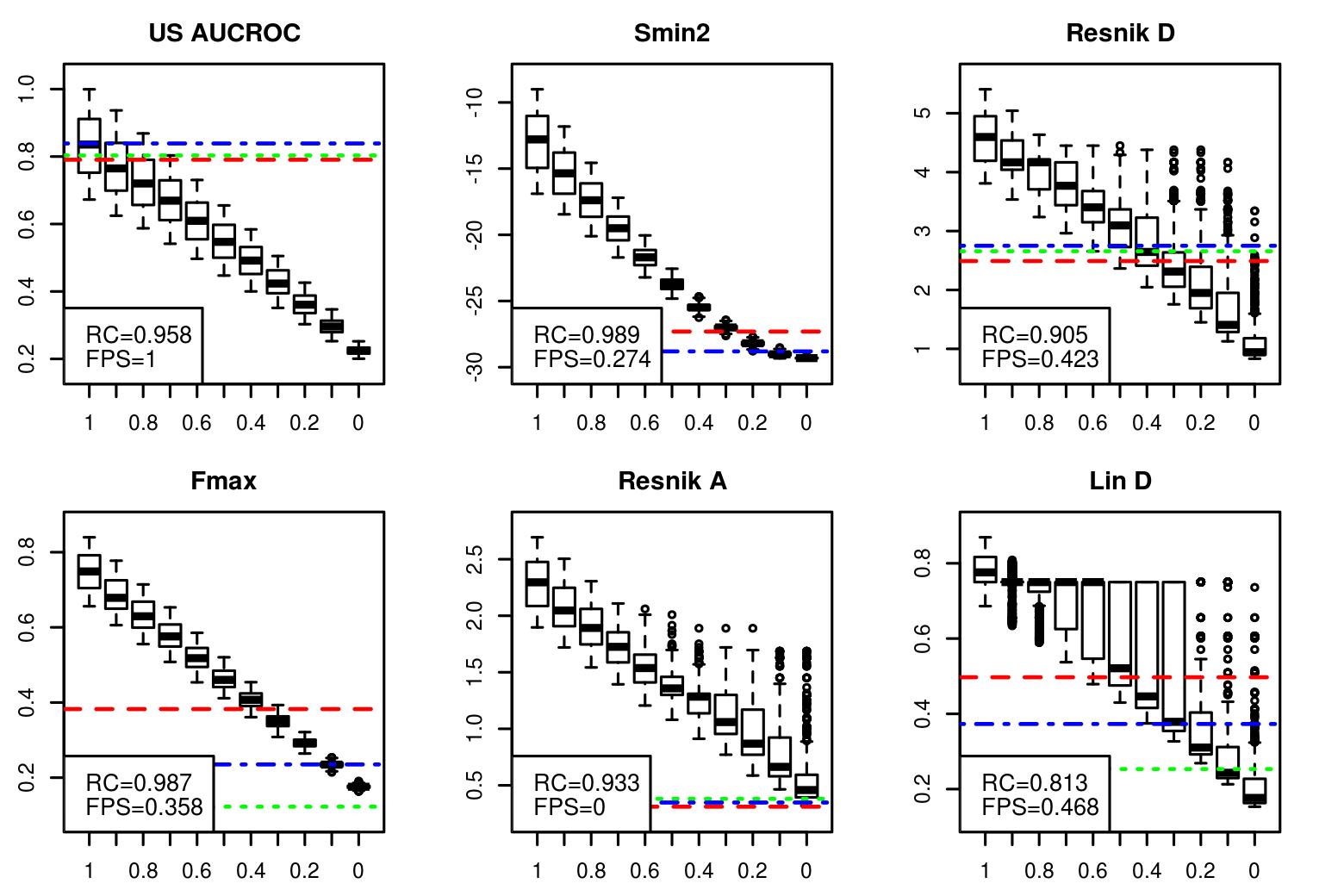

### uniprot.1000_boxplot1.jpeg

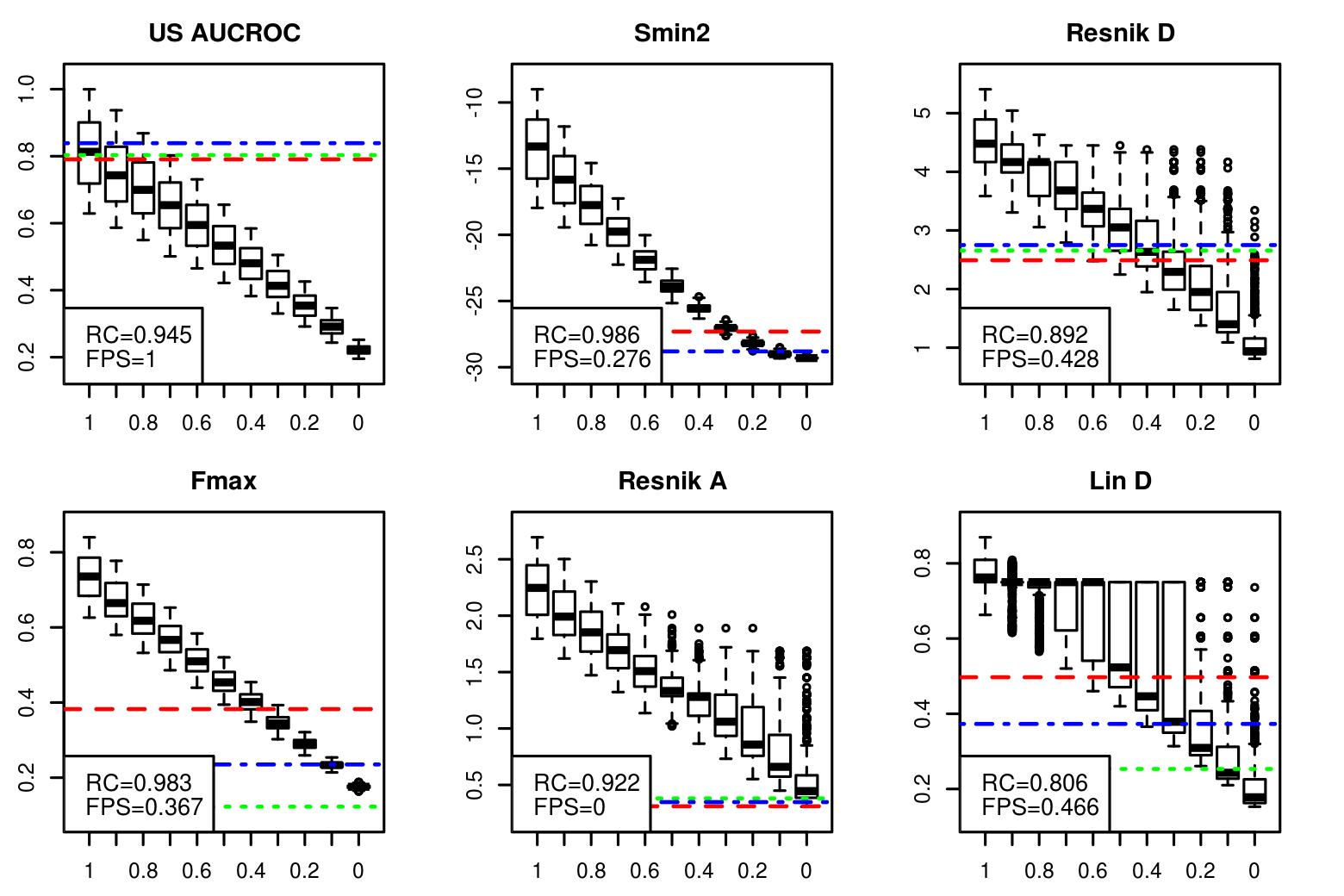

### uniprot.1000_boxplot1.jpeg

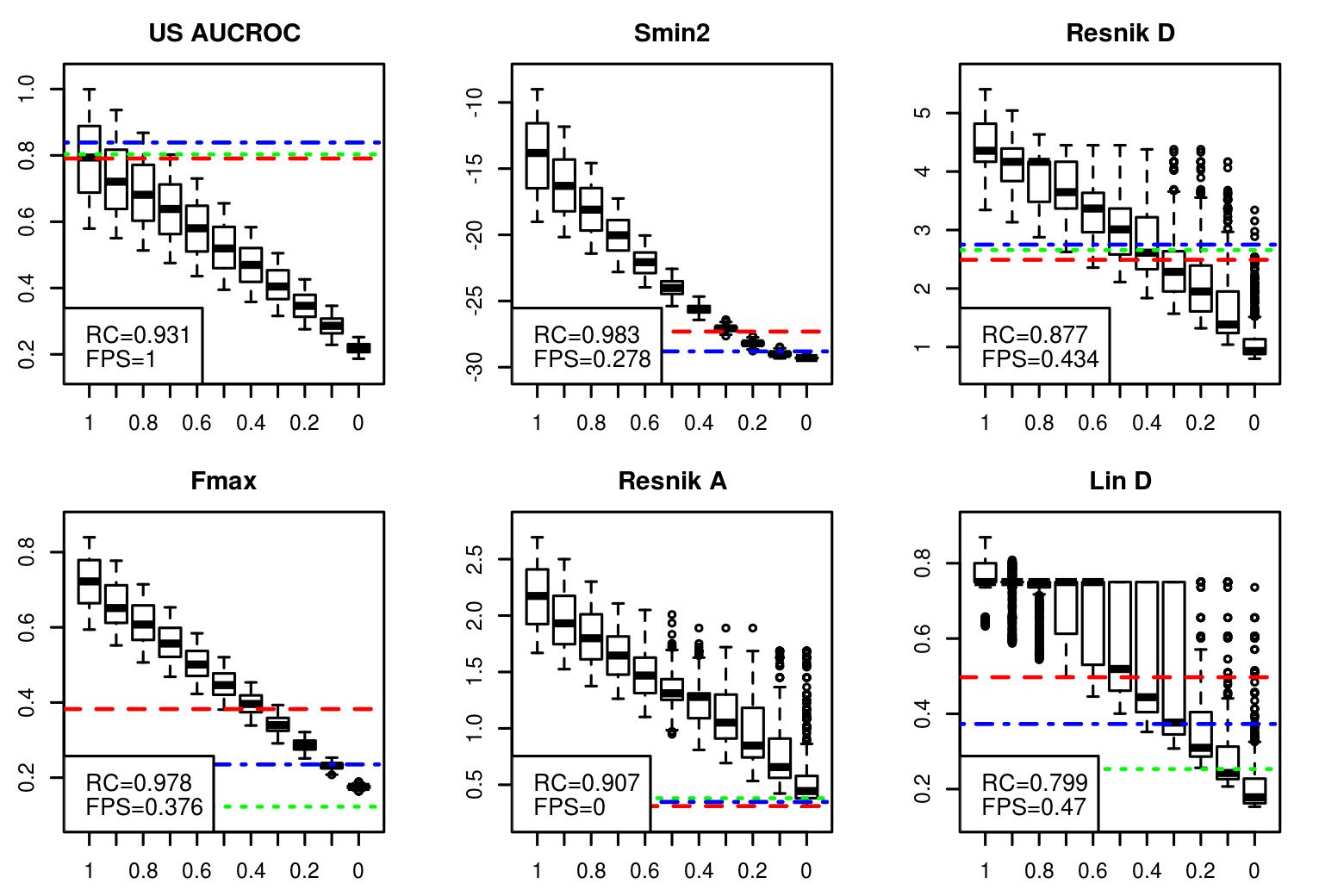

### uniprot.1000_boxplot2.jpeg

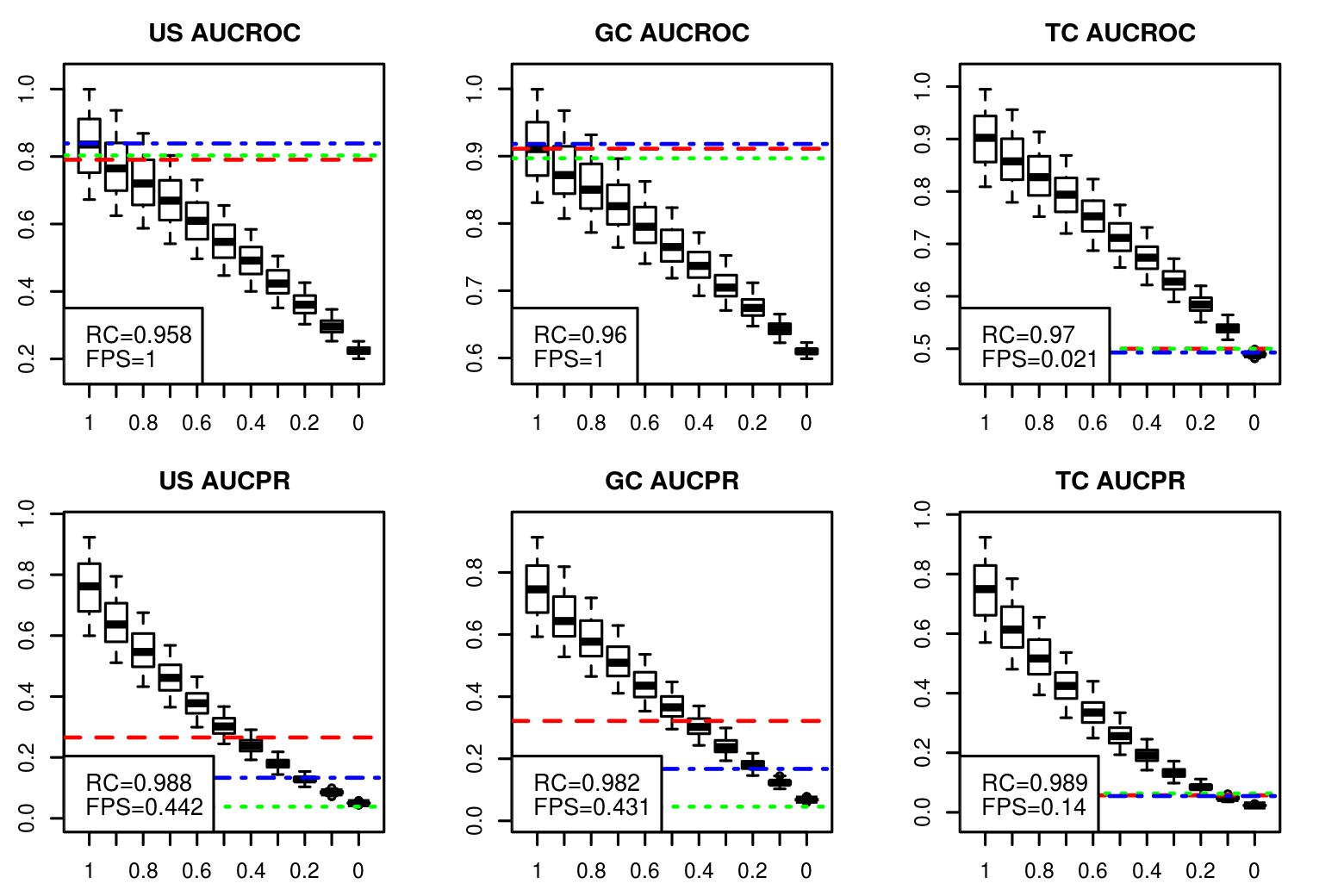

### uniprot.1000_boxplot2.jpeg

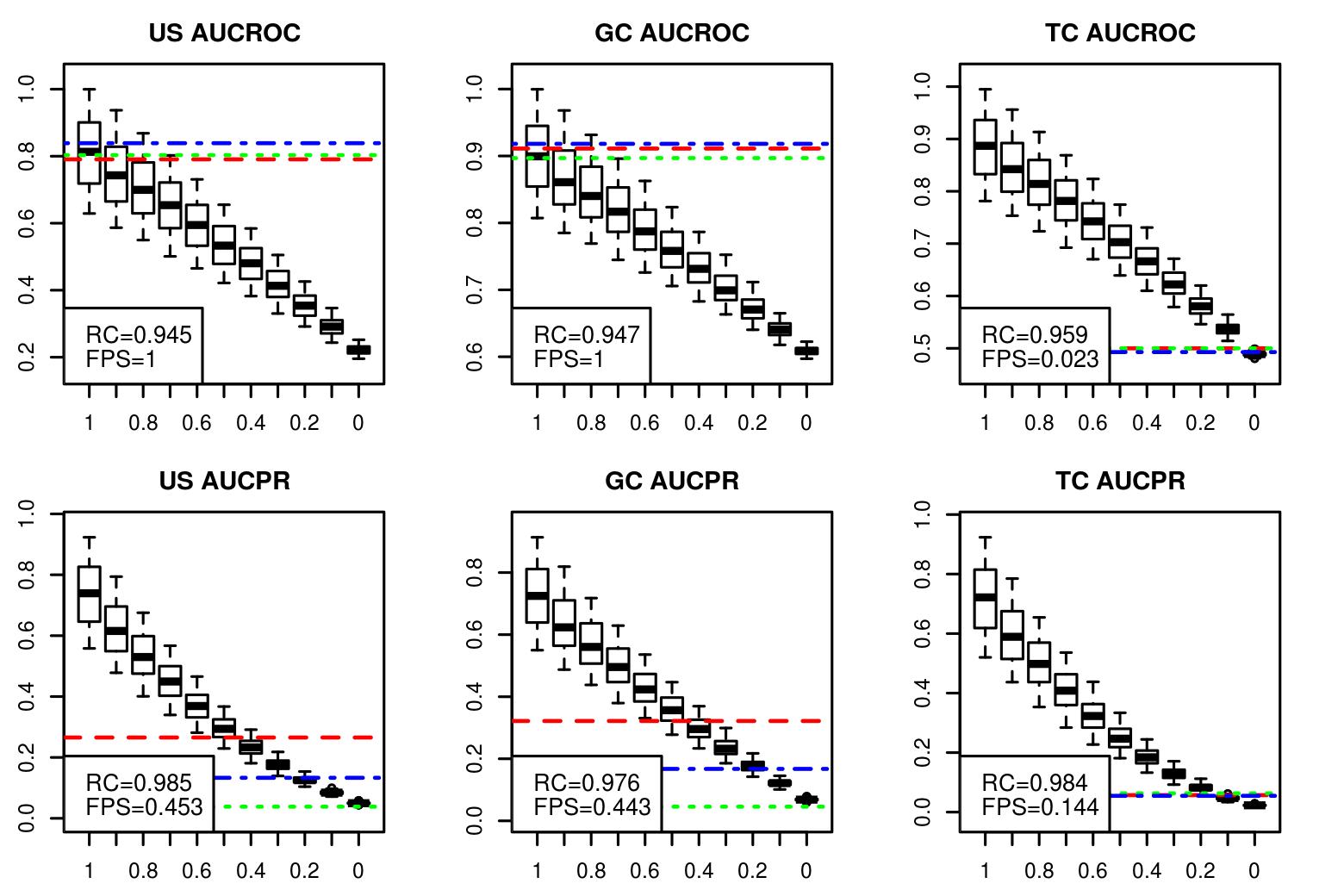

### uniprot.1000_boxplot2.jpeg

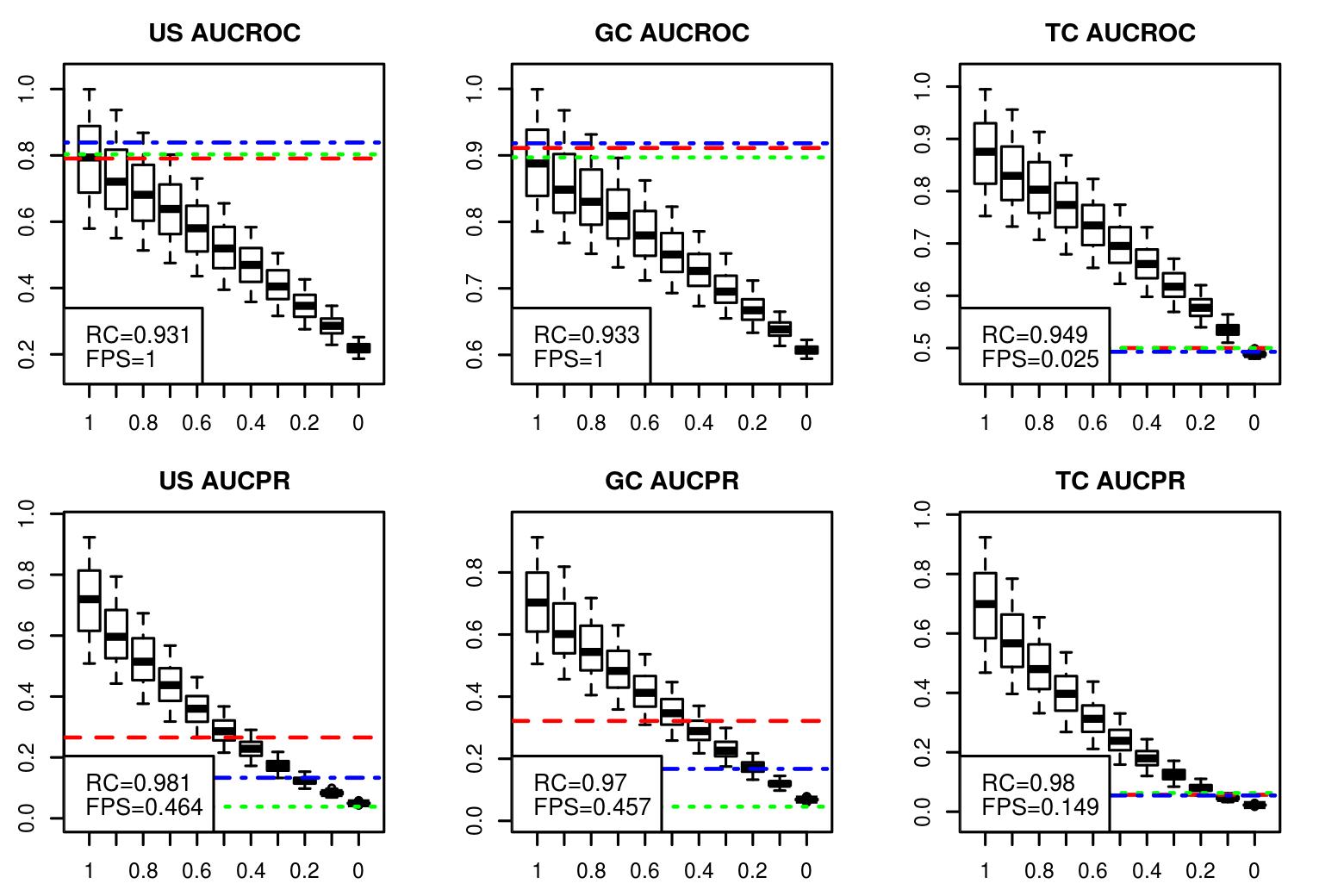

### uniprot.1000_boxplot3.jpeg

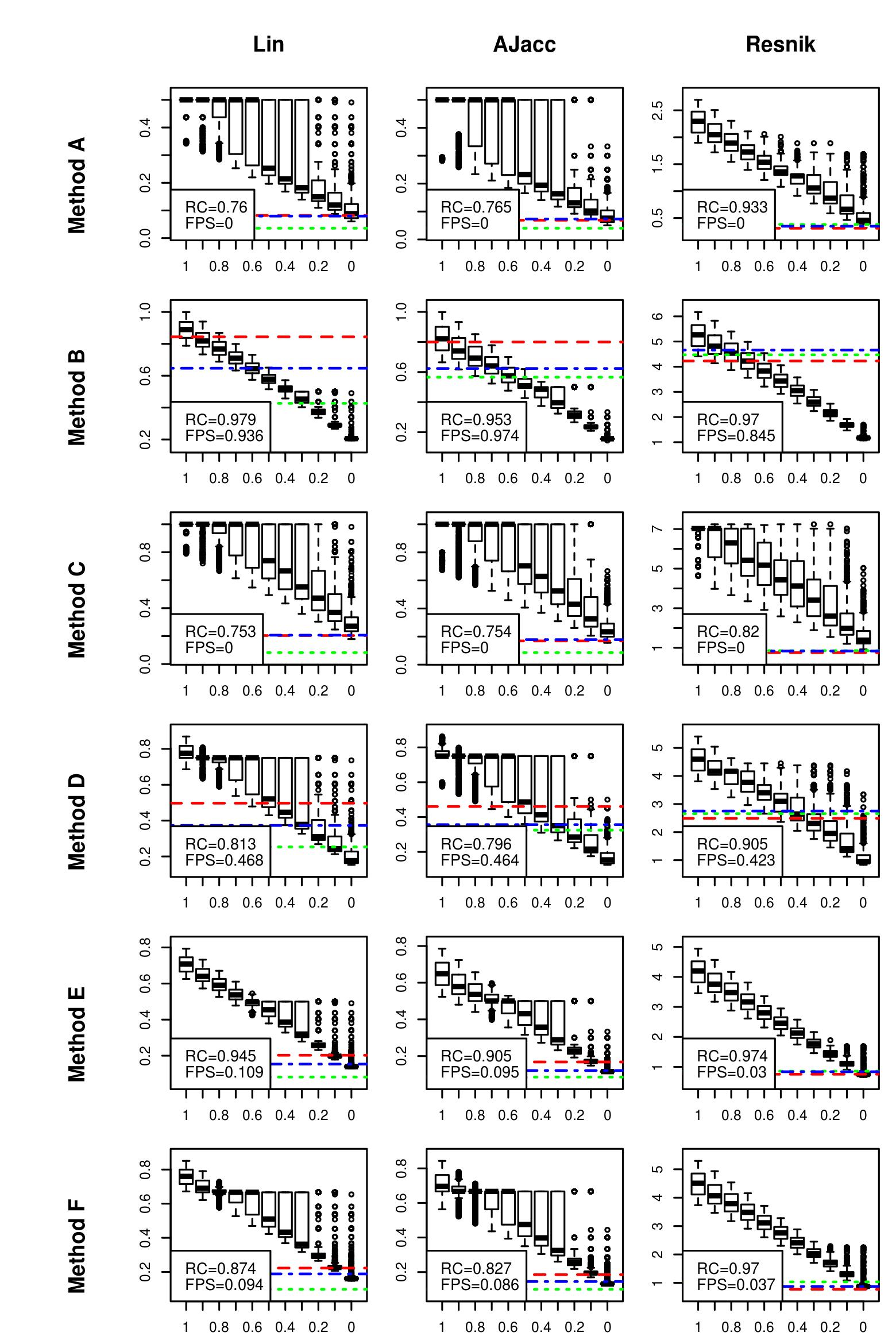

### uniprot.1000_boxplot3.jpeg

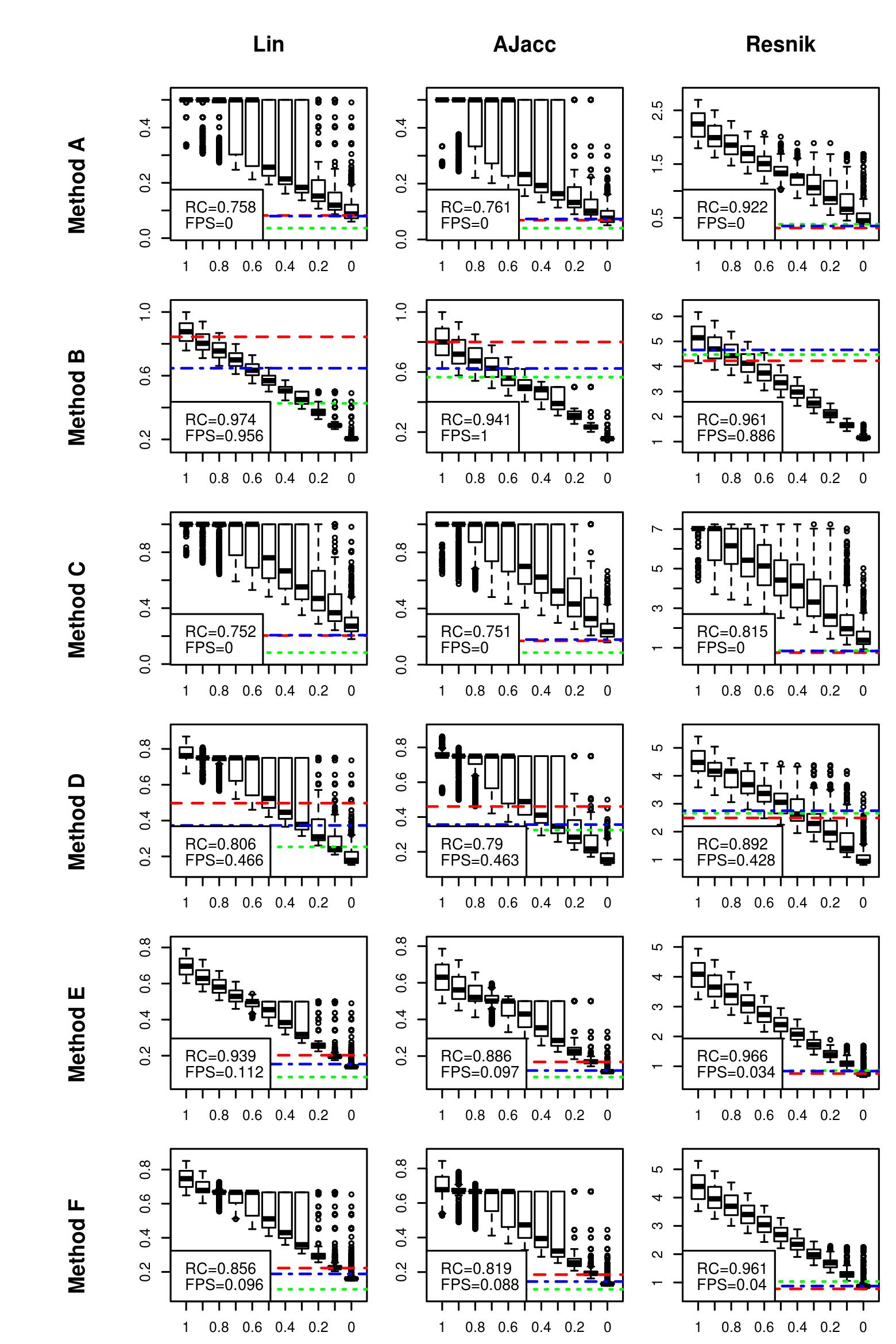

### uniprot.1000_boxplot3.jpeg

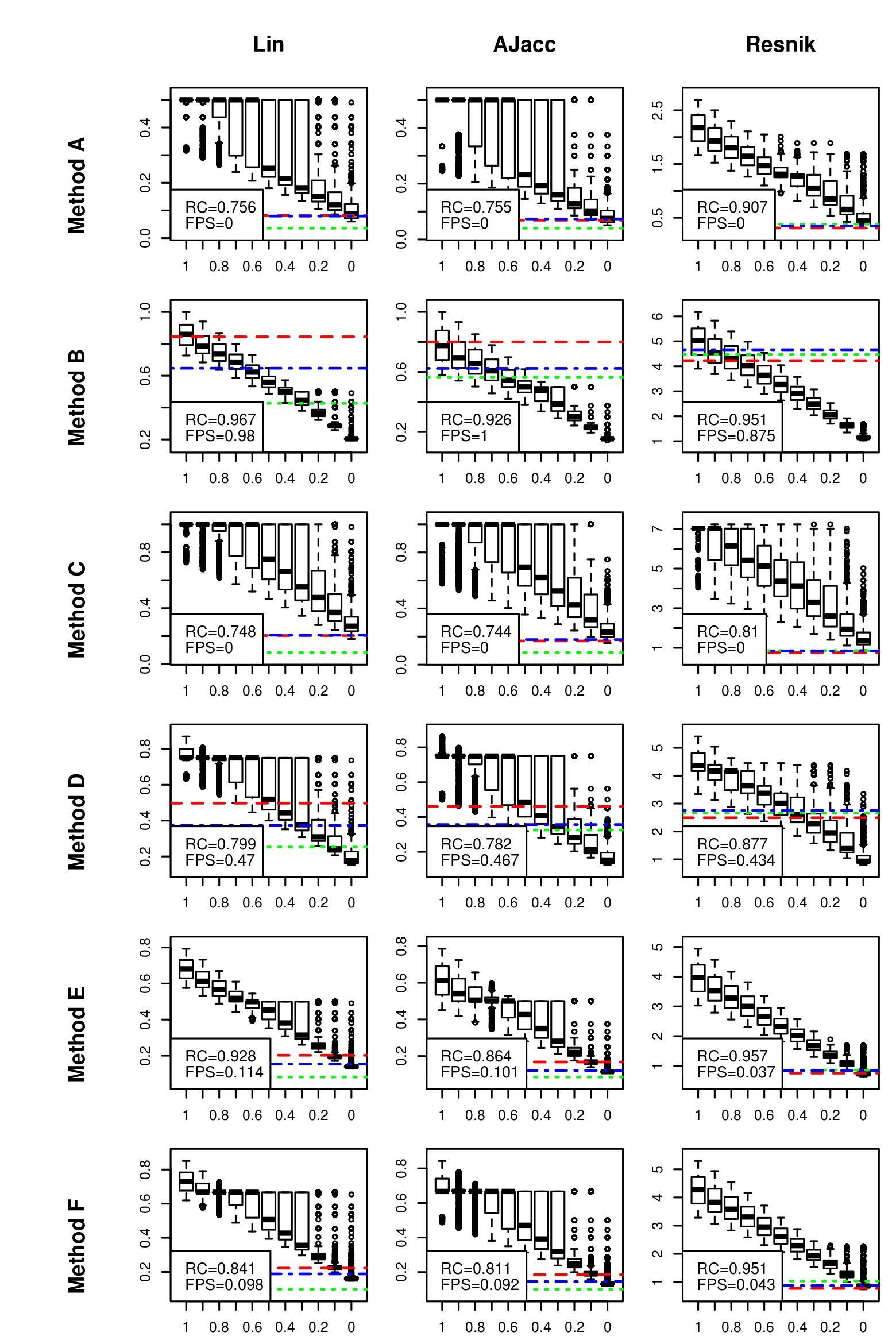

### uniprot.1000_boxplot3b.jpeg

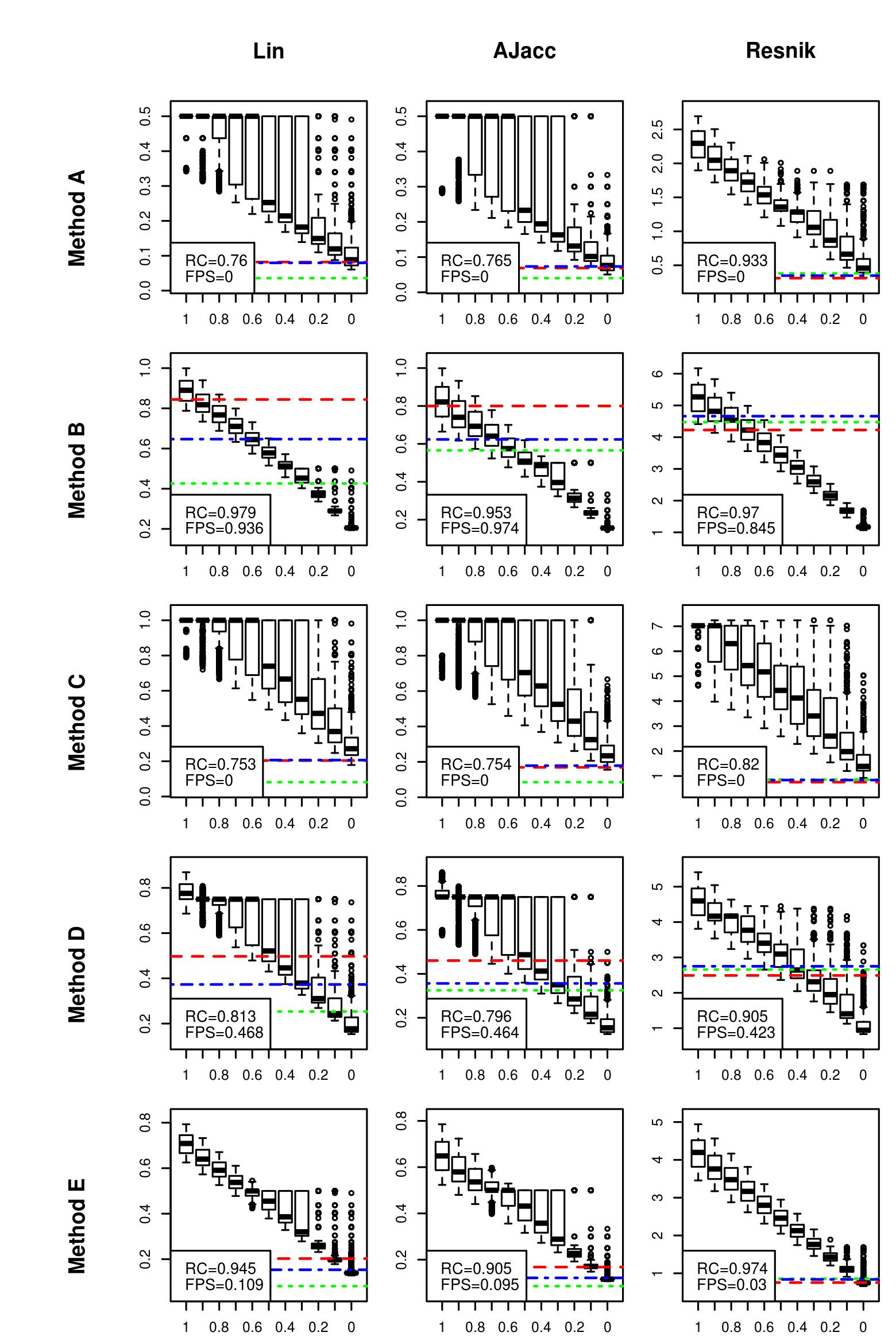

### uniprot.1000_boxplot3b.jpeg

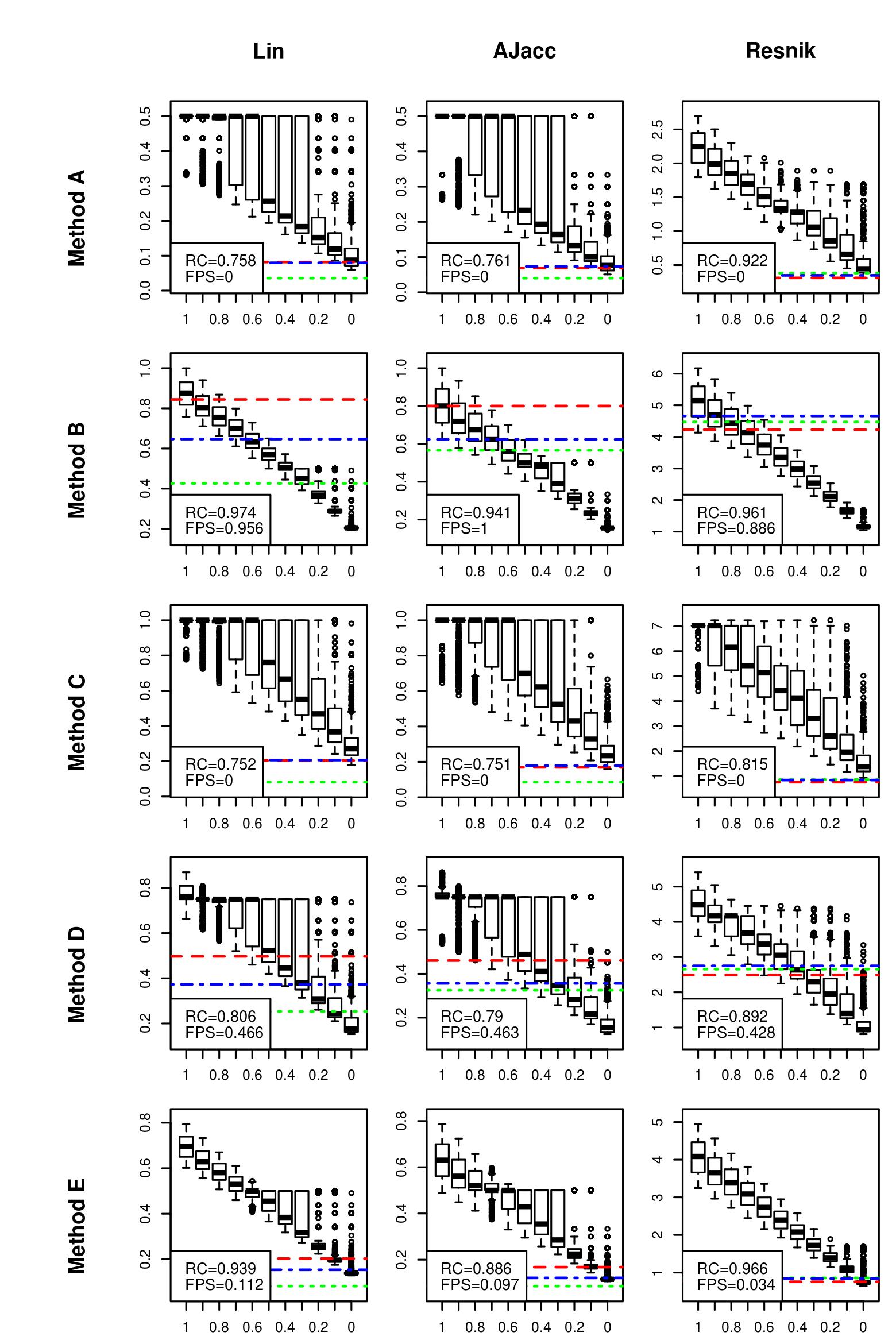

### uniprot.1000_boxplot3b.jpeg

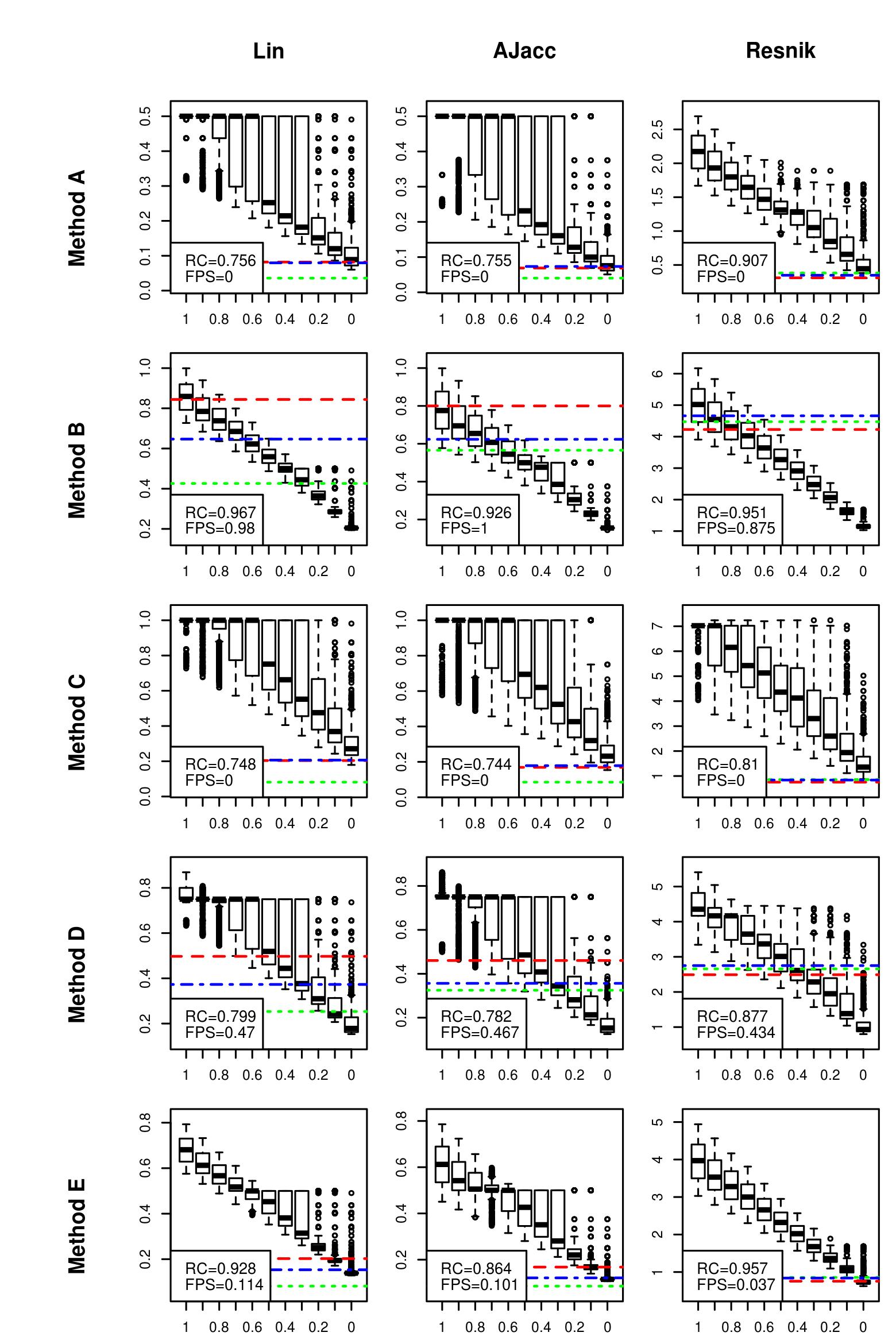

### uniprot.1000_boxplot3c.jpeg

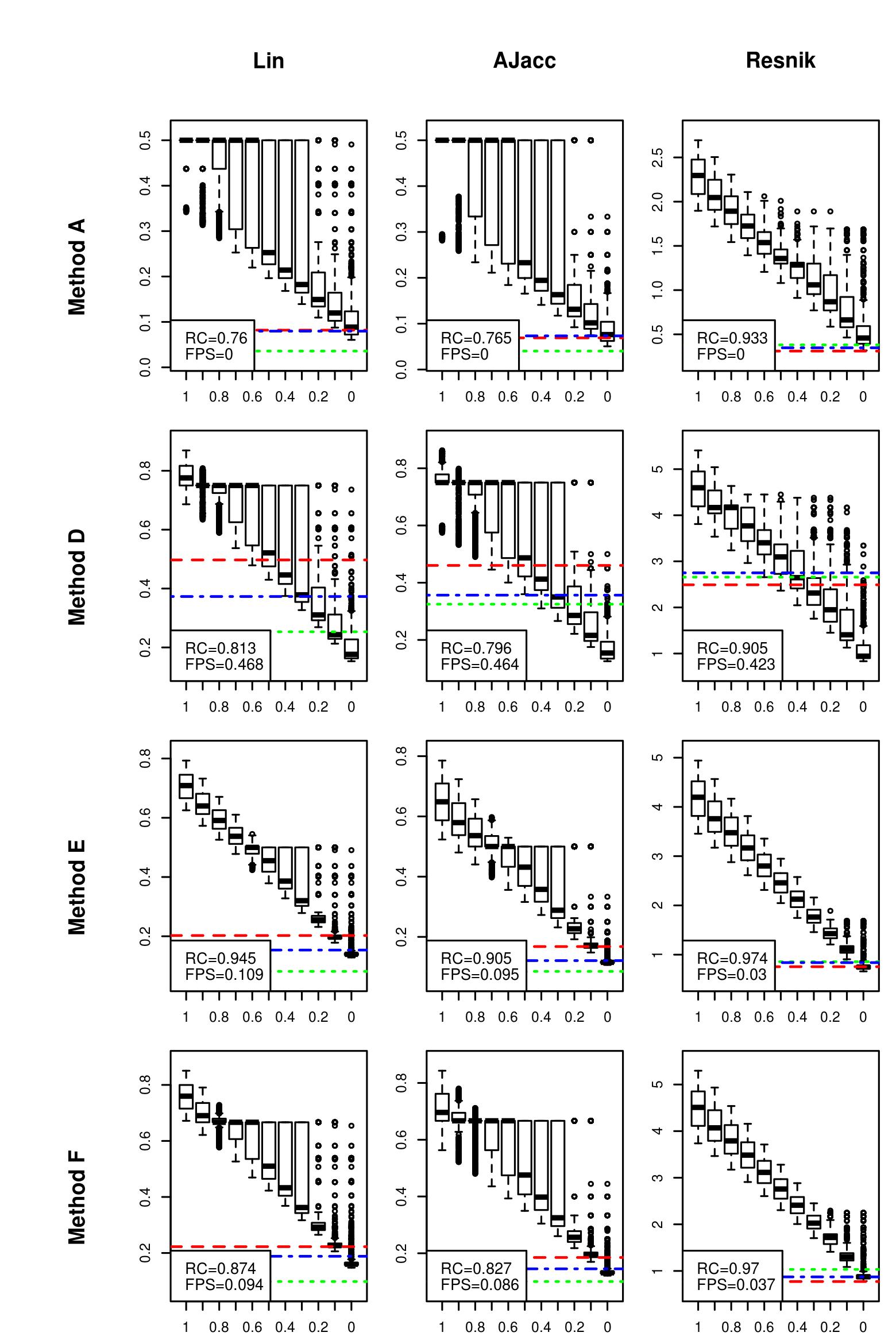

### uniprot.1000_boxplot3c.jpeg

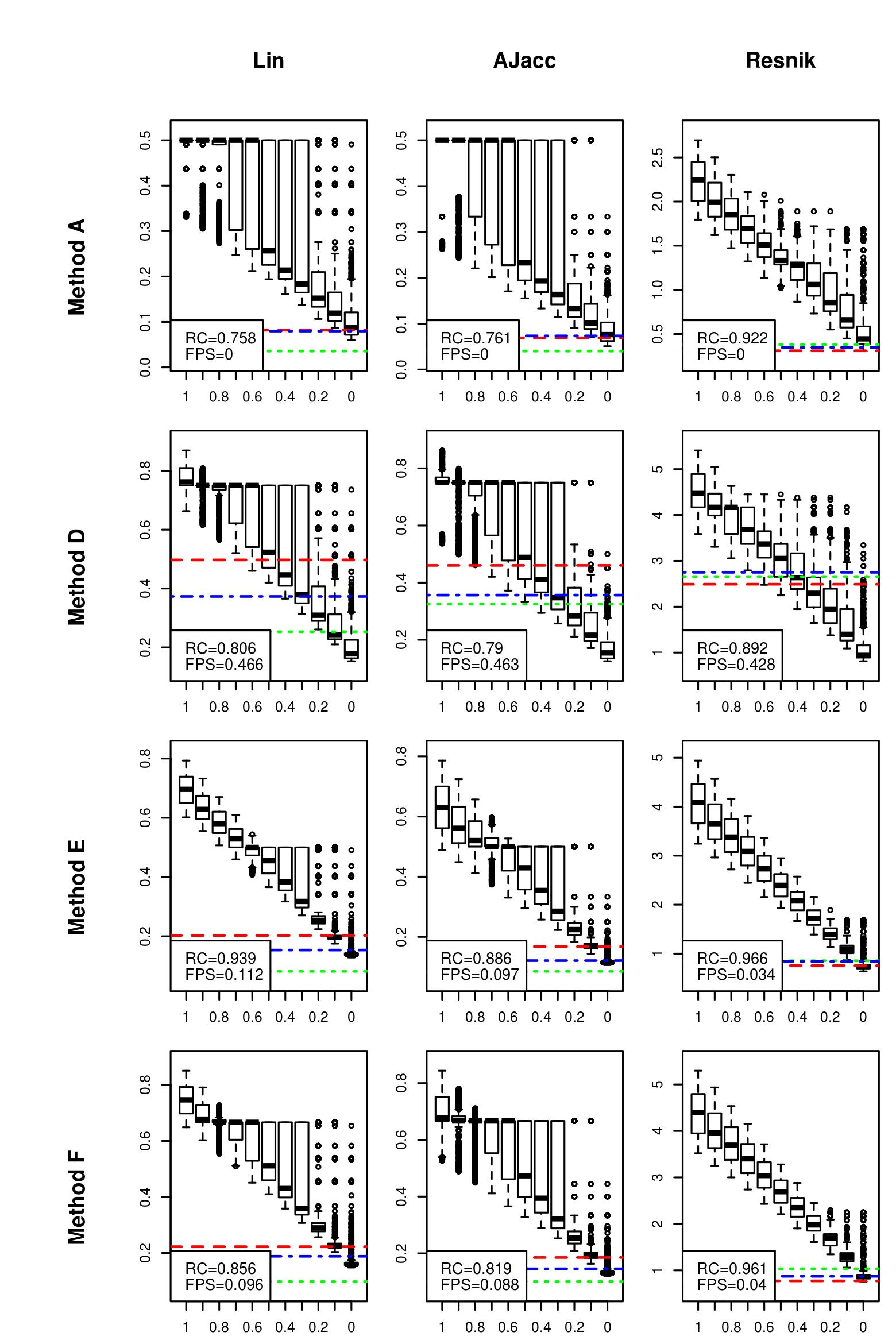

### uniprot.1000_boxplot3c.jpeg

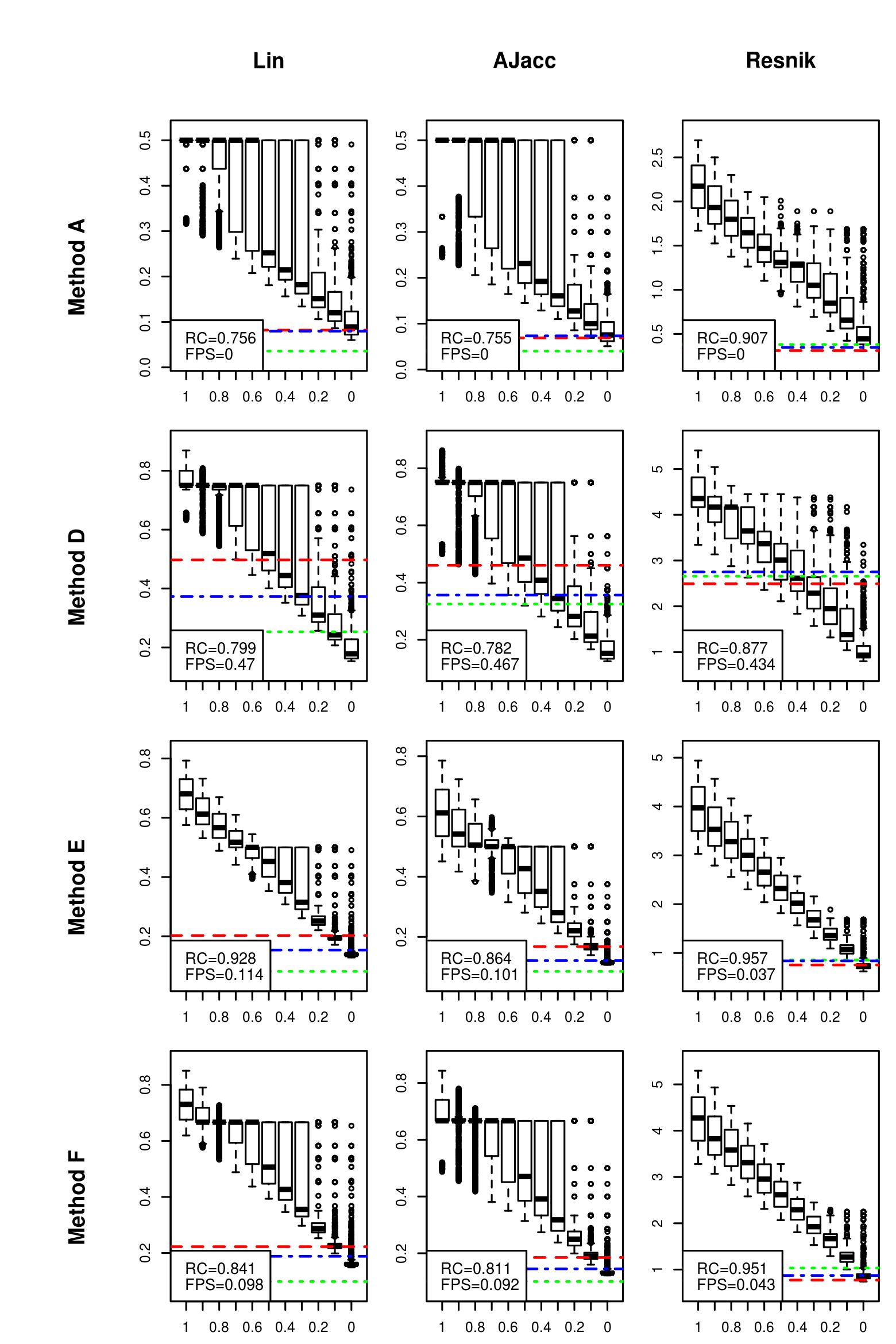

### uniprot.1000_boxplot4.jpeg

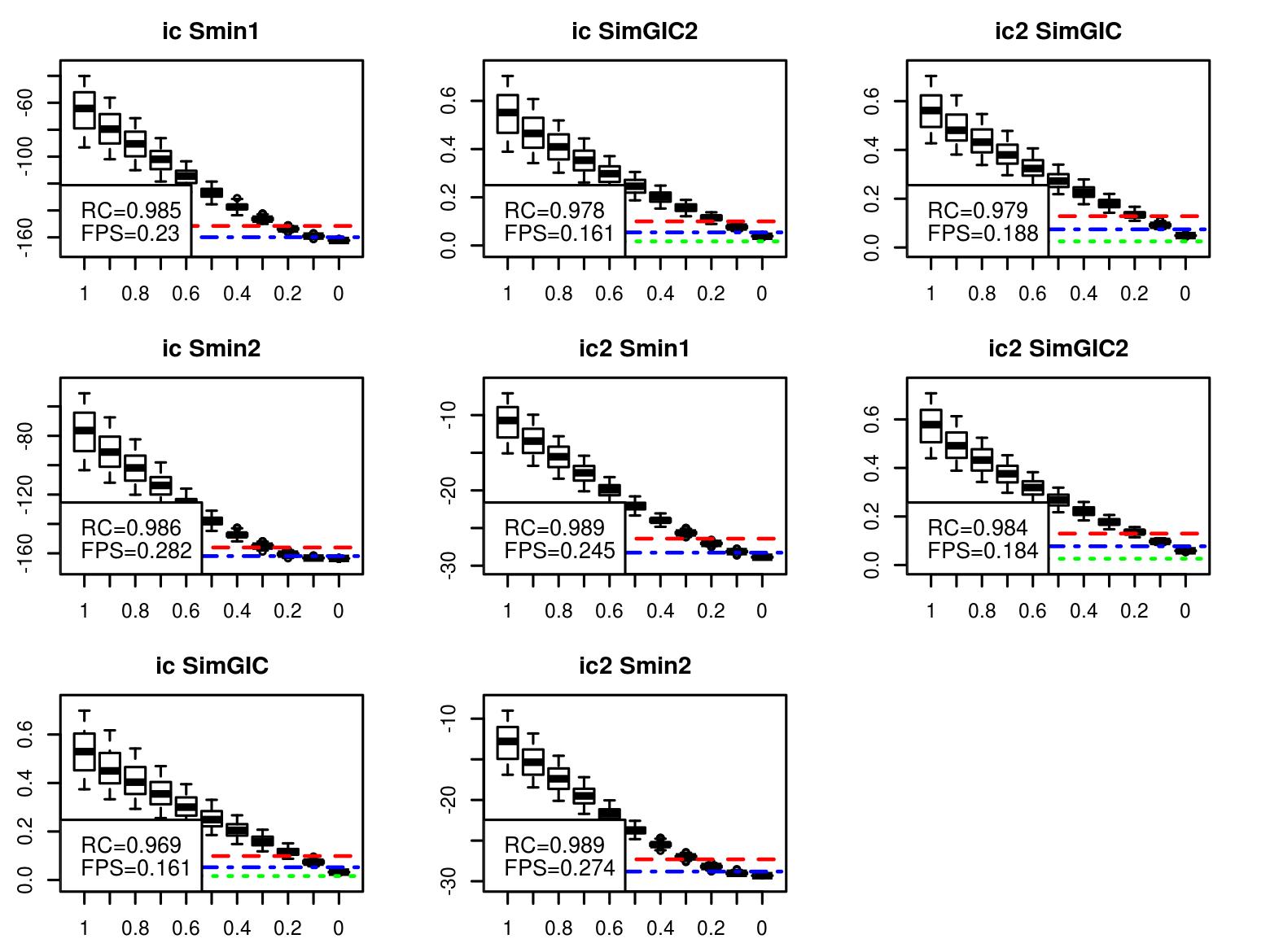

### uniprot.1000_boxplot4.jpeg

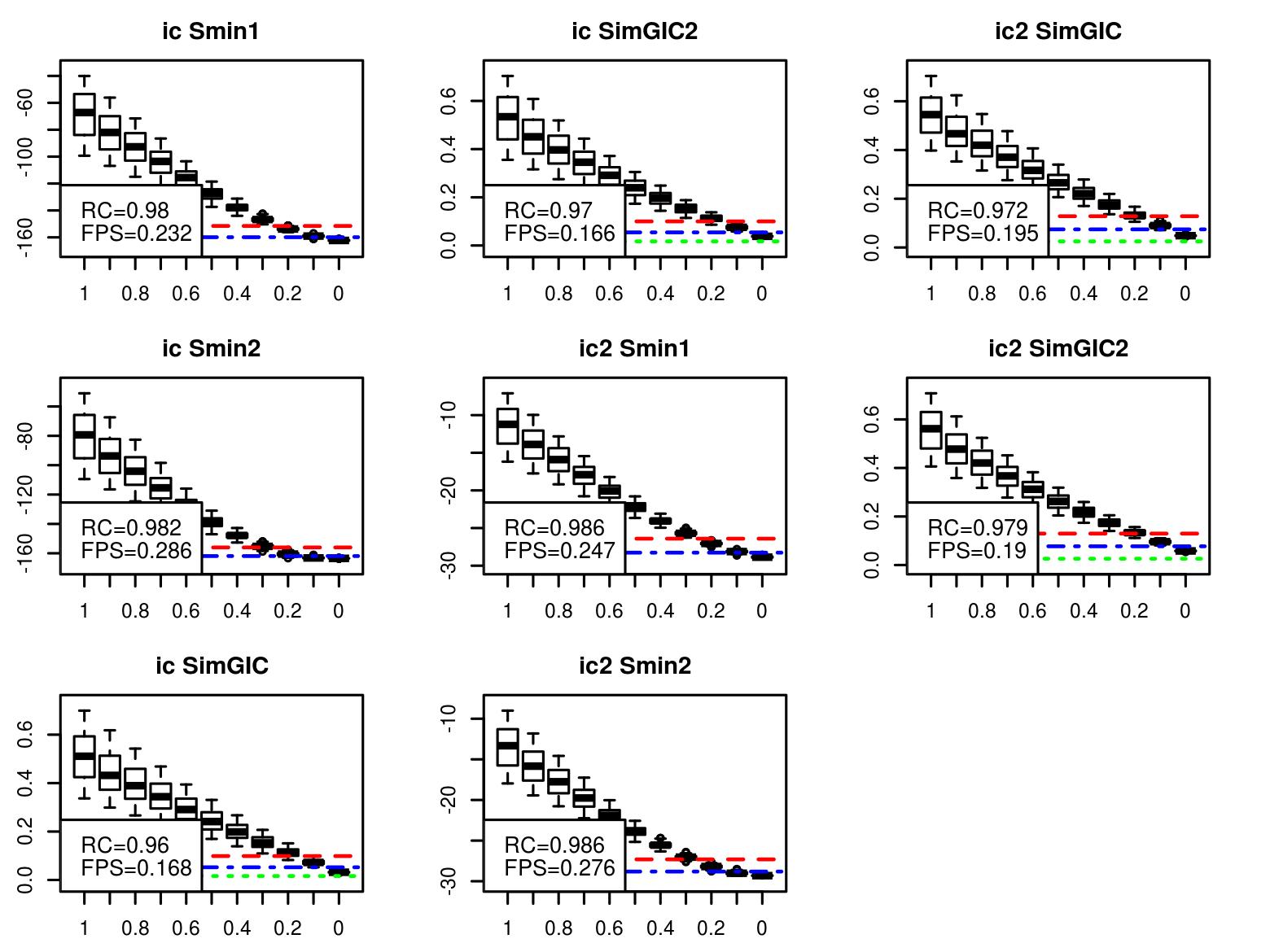

### uniprot.1000_boxplot4.jpeg

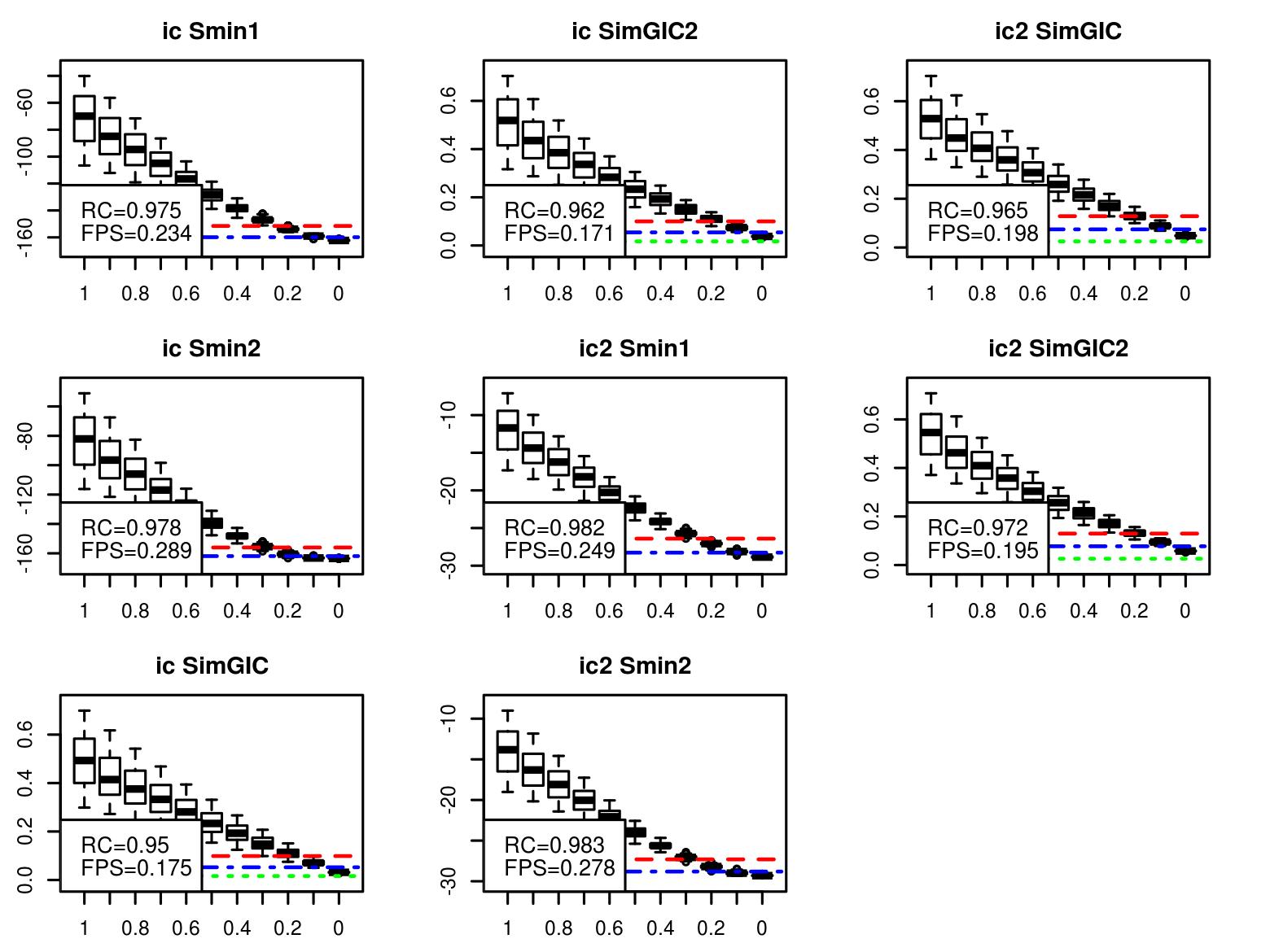

### uniprot.1000_boxplot4b.jpeg

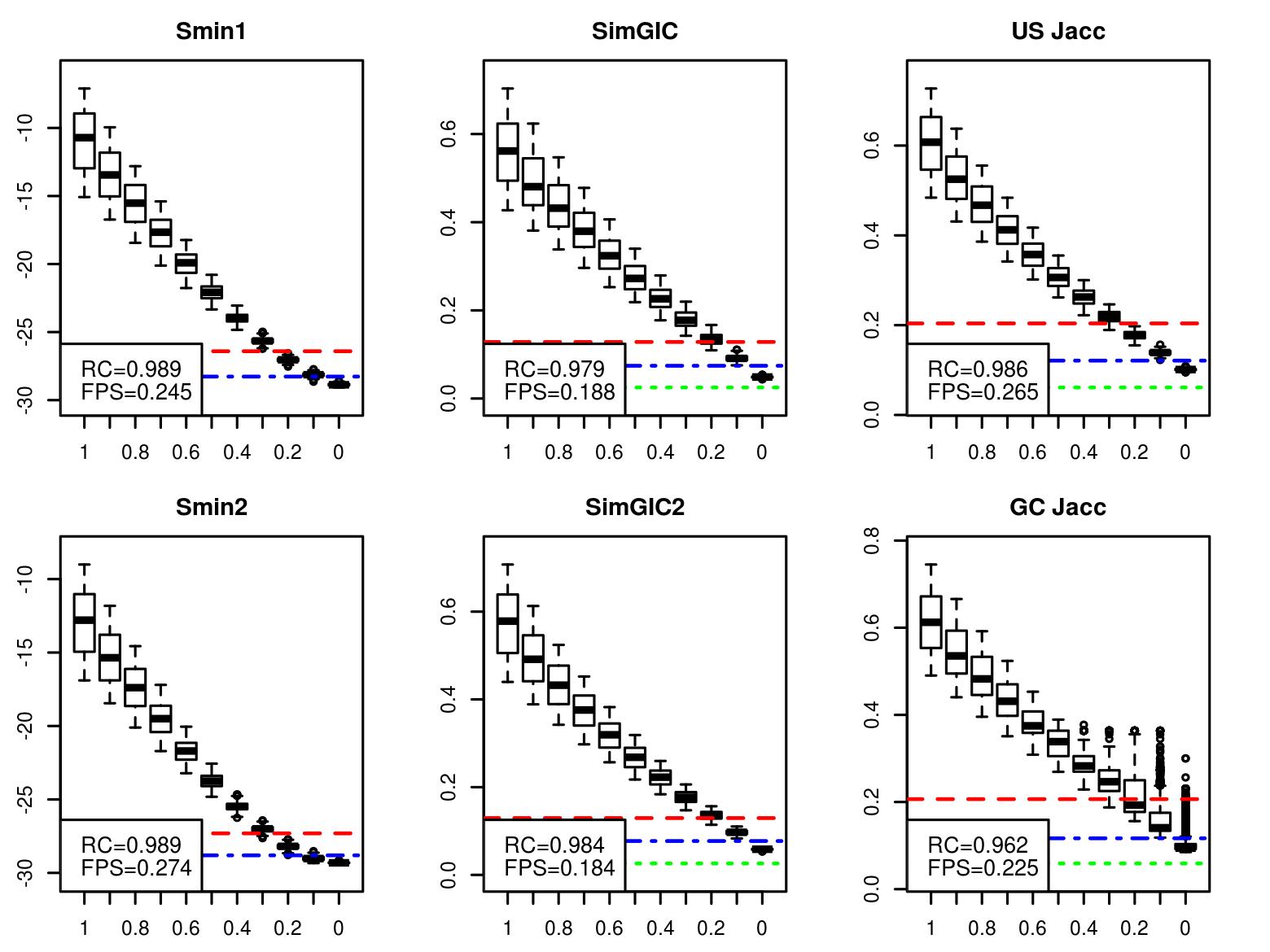

### uniprot.1000_boxplot4b.jpeg

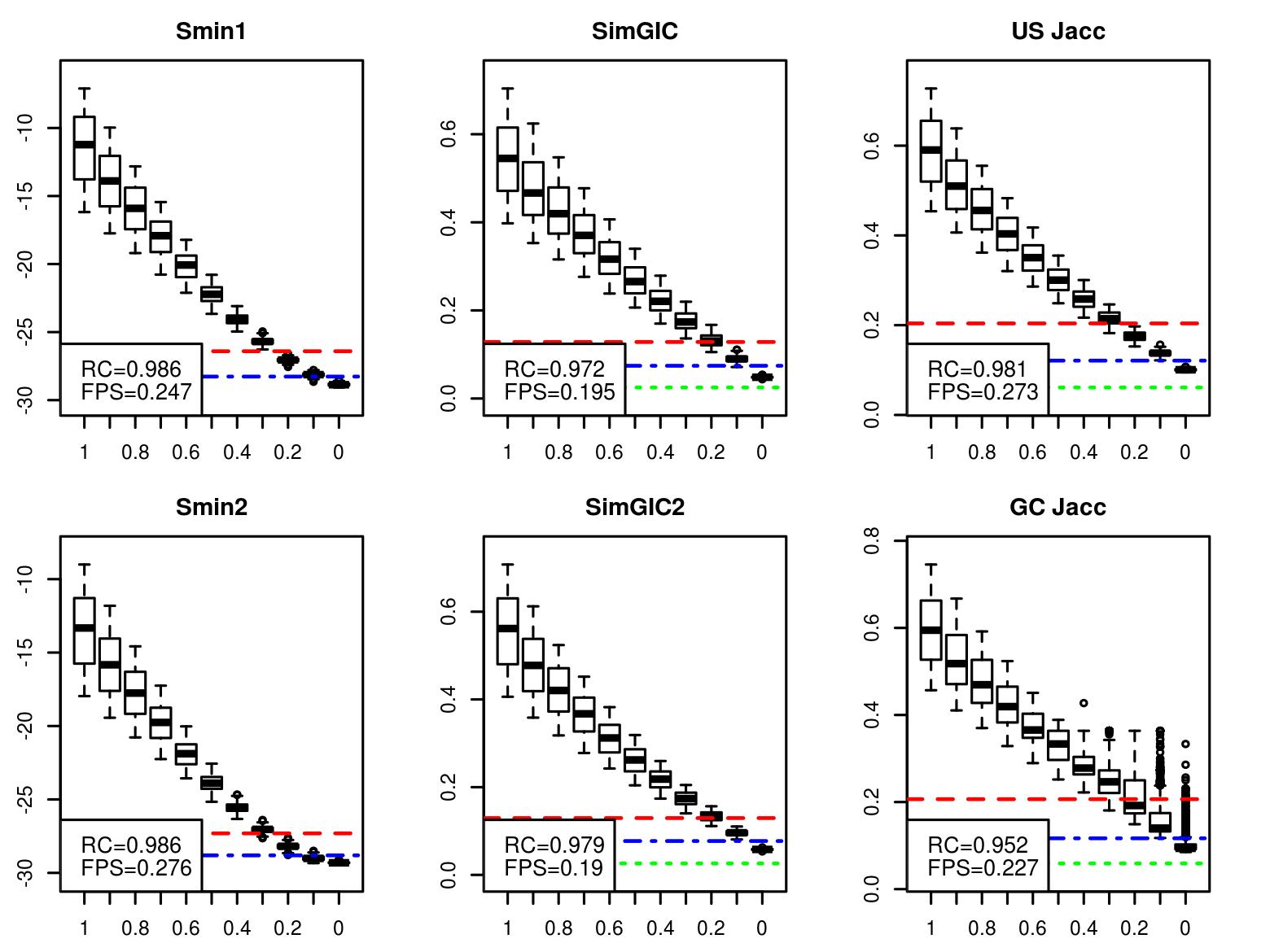

### uniprot.1000_boxplot4b.jpeg

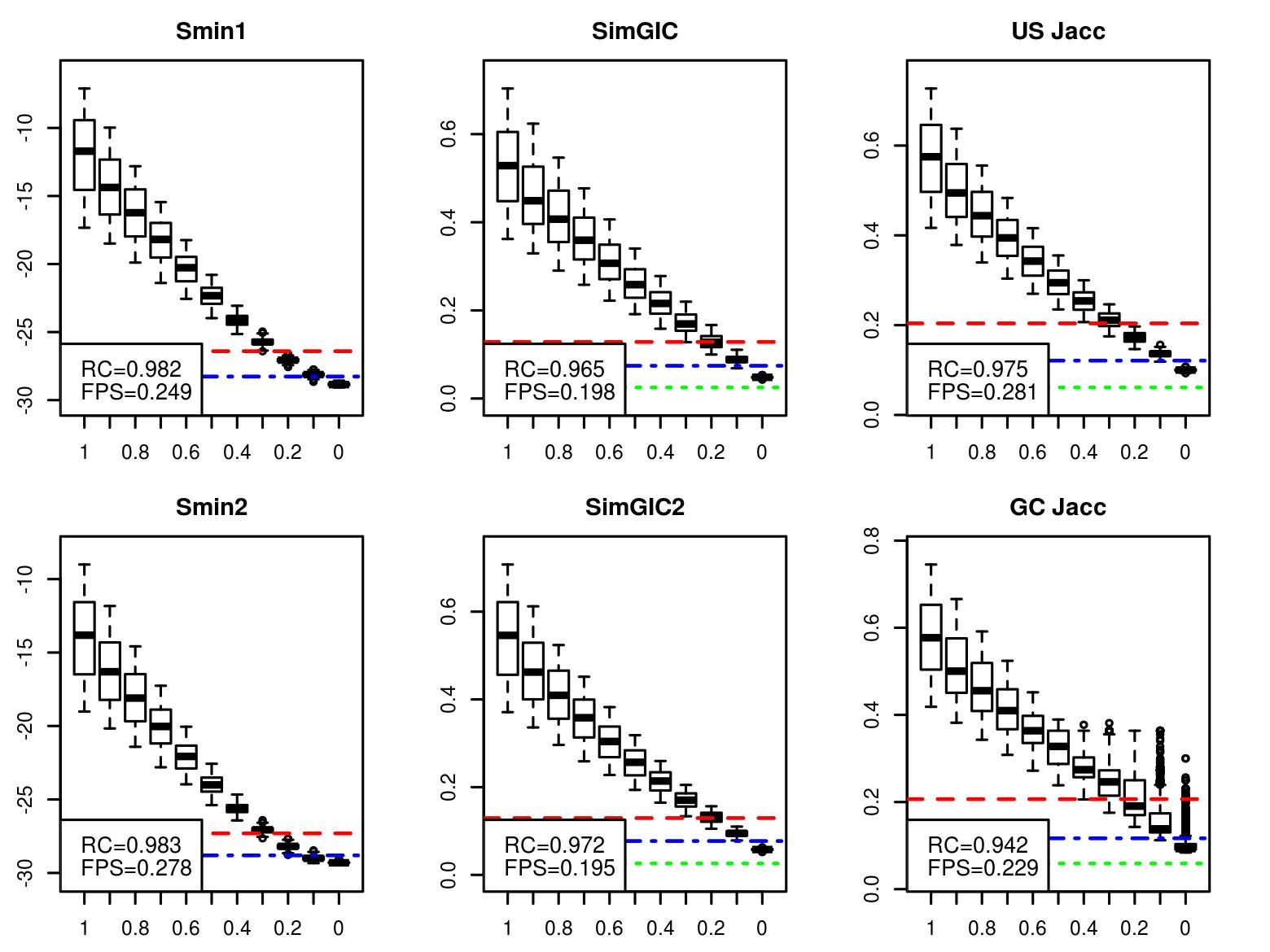

### uniprot.1000_scattered_labels.jpeg

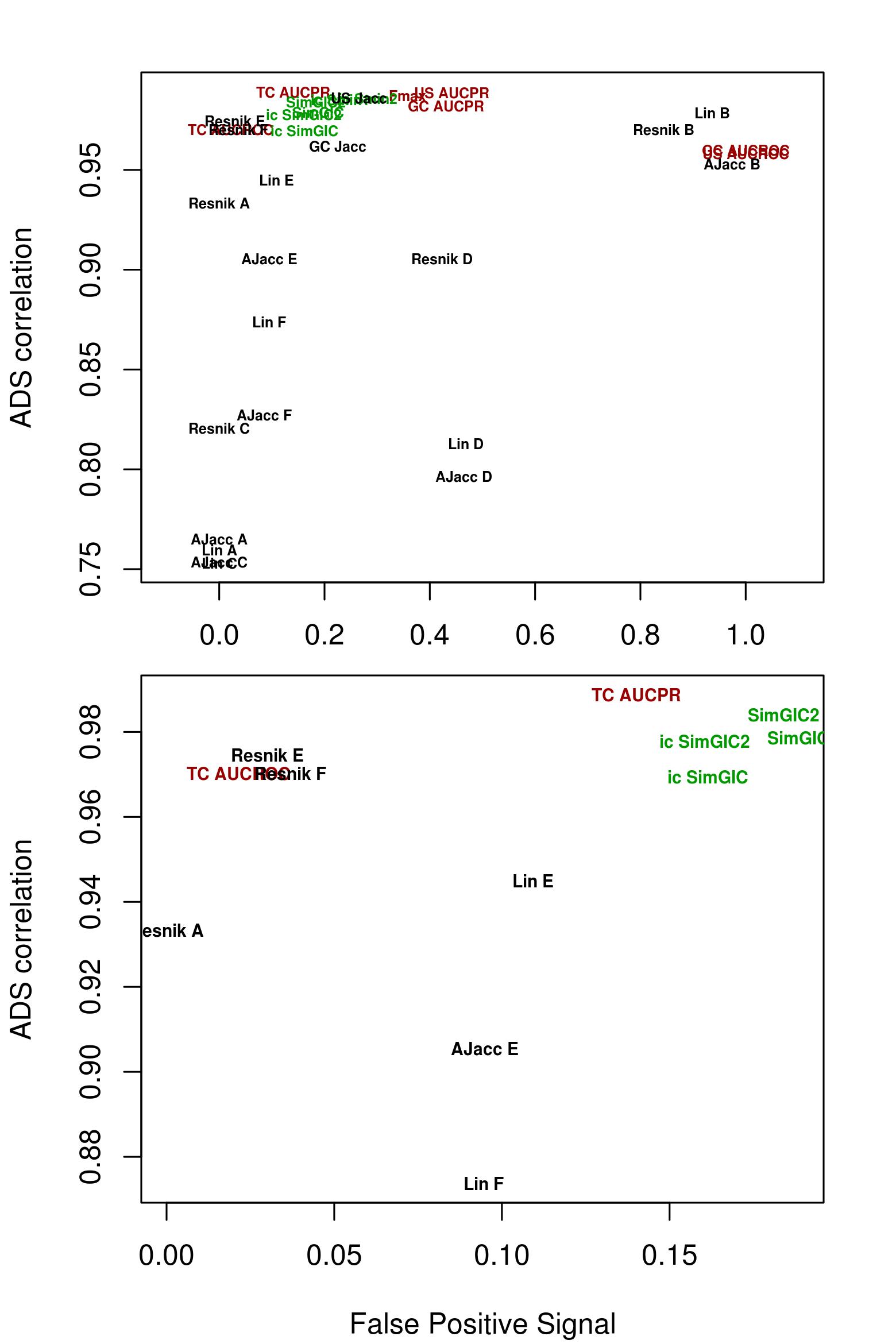

### uniprot.1000_scattered_labels.jpeg

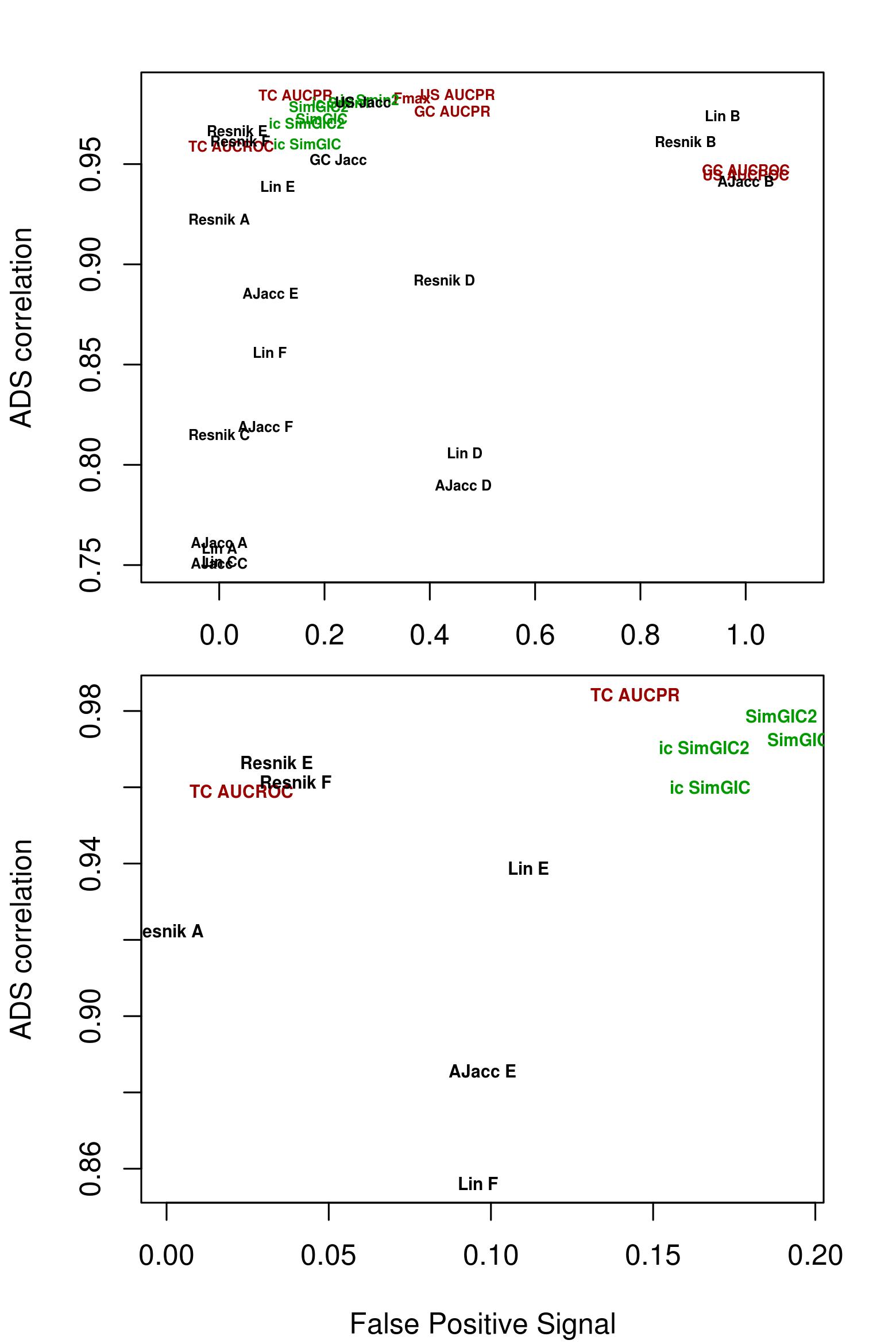

### uniprot.1000_scattered_labels.jpeg

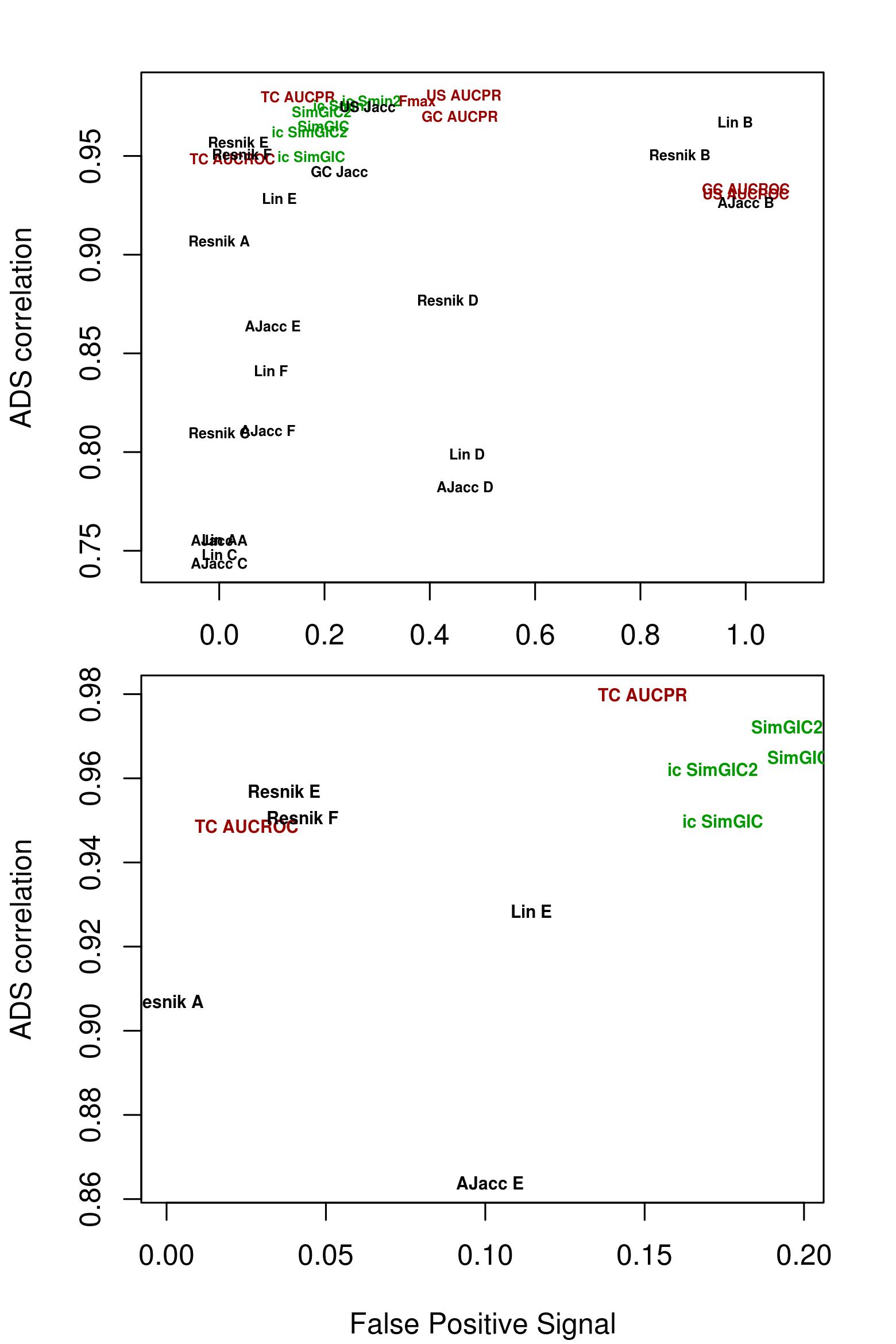
