## Supplementary Text 1 for "Novel Comparison of Evaluation Metrics for Gene Ontology Classifiers Reveals Drastic Performance Differences"

```
Data: Correct GO predictions, T, for the selected set of genes
Data: Key list representing parent classes for every GO class
parameter: Vector of L Noise Levels, NL
           default: NL = [0, 0.1, 0.2...1]
parameter: Number of repetitions, K, within each NL
           default: K = 1000
Result: K by L Matrix, scores with selected Evaluation Metric
begin
    Define L as the length of NL ;
    Define output as K by L matrix ;
    Set N_neg = 4 (GO classes per gene in neg. set) ;
    foreach l in [1, 2, .., L] do
        p = NL[l], current noise level ;

        #1. Create positive and negative data
        P_pos = GeneratePosData(T, p) ;
        P_neg = GenerateNegData(T, N_neg) ;

        #2. Create artificial classifier scores to sets
        foreach row in P_pos do
            Select predictor score, s, from Normal(μ = 1, σ = 0.5);
            Add s to the current row;
        end
        foreach row in P_neg do
            Select predictor score, s, from Normal(μ = −1, σ = 0.5);
            Add s to the current row;
        end

        # 3. Combine positive and negative datasets
        P = P_pos ∪ P_neg ;

        # 4. Run Evaluation Metric, EvM
        Output[k, l] = EvM(P, T);
    end
end
```

**Algorithm 1:** Artificial Dilution Series (ADS) pipeline. Code uses subfunctions *GeneratePosData* and *GenerateNegData*, explained later.

```
Data: Correct GO predictions, T, for the selected set of genes
Data: Key list representing parent classes for every GO class
parameter: Noise proportion, p
output    : Modified GO predictions, P
begin
    Define N_T as the size of T ;
    P = T;
    th_noise = round(N_T * p), the size threshold for the Noise Set ;

    # Shifting step
    Define N_shift, a random integer between 0 and N_T;
    Select P_shift, a random subset of P of size N_shift;
    foreach row in P_shift do
        Replace GO class with one of its nearest parents;
    end

    # Permutation step
    Define NoiseSet = [] ;
    Define N_err = 0 ;
    while th_noise > N_err do
        Select two random rows A and B from P \ NoiseSet ;
        # Genes of these rows will be gene_A and gene_B ;
        # GO classes of these rows will be GO_A and GO_B ;
        if IsNoiseClass(gene_A,GO_B) and IsNoiseClass(gene_B,GO_A) then
            Swap GO classes between rows A and B in P ;
            Add A and B to NoiseSet ;
            N_err = N_err + 2 ;
        end
    end
    return P ;
end
```

**Algorithm 2:** GeneratePosData: Generation of positive data with noise proportion $p$ in ADS pipeline. Code uses IsNoiseClass function, explained later.

```
Data: Correct GO predictions, T
parameter: N_neg, number of reported negative GO classes per gene
output    : Negative GO predictions, P_neg
begin
    Define genes(T), the set of unique gene names in T ;
    Define P_neg = [] ;
    foreach gene in genes(T) do
        GO_count = 0 ;
        while GO_count < N_neg do
            pick random GO class, GO_rand, from GO structure ;
            if IsNoiseClass(GO_rand, gene) then
                Add (gene, GO_rand) as item to P_neg ;
                GO_count = GO_count + 1 ;
            end
        end
    end
    Return P_neg ;
end
```

**Algorithm 3:** GenerateNegData: Generation of negative data in ADS pipeline. Code uses IsNoiseClass function, explained later.

**Data:** Correct GO predictions, $T$
**Data:** Key list representing all ancestor classes for every GO class
**Data:** Evaluated GO class, $GO_{eval}$
**Data:** Evaluated gene, $gene_{eval}$
**parameter:** Noise threshold, $th$
default: $th = 0.1$
**output** : TRUE or FALSE

```
begin
    Get the set of ancestor classes, A_eval, for GO_eval ;
    Define J_max = 0 ;
    foreach row in T do
        Let gene_test be the gene of the current row ;
        if gene_test is gene_eval then
            Let GO_test be the GO class of the current row;
            Get the set of ancestor classes, A_test, for GO_test ;
            Calculate J_test, Jaccard correlation between A_test and A_eval;
            if J_test > J_max then
                J_max = J_test
            end
        end
    end
    if J_max < th then
        Return TRUE ;
    else
        Return FALSE ;
    end
end
```

**Algorithm 4:** IsNoiseClass. This algorithm tests if GO class qualifies as noise. This is tested against the annotations of the selected gene
