## Supplementary Text 2 for "Novel Comparison of Evaluation Metrics for Gene Ontology Classifiers Reveals Drastic Performance Differences"

### Artificial Dilution Series
Supplementary Information

Ilya Plyusnin, Liisa Holm, Petri Törönen

#### 1 Introduction

This is the supplementary text for the main article, "Novel Comparison of Evaluation Metrics for Gene Ontology Classifiers Reveals Drastic Performance Differences". Here we compare this work to related previous research. We give an in-detail descriptions of the compared evaluation metrics. We also give definitions of signal and noise in the Artificial Dilution Series (ADS) and describe the tested methods for combining semantic similarities.

We also discuss some alternative evaluation metrics from the literature. We want to underline that we had to limit the number of compared evaluation metrics and we included only the most relevant ones and their variations to comparison.

#### 2 Comparing ADS with related research

To our knowledge, there has been little research on evaluation metrics with Gene Ontology (GO). Clark and Radivojac compare seven methods by looking at the threshold position that optimizes each evaluation metric, and explore if the selected position is rational for classification purposes [1]. GO Semantic similarities [2] have been evaluated using correlation against other datasets, like sequence similarity [3], but again their performance as classifier evaluation metric has not been thoroughly tested. These tests lack the clear reference point, as the ADS noise level in our work, for comparing EvMs.

Machine learning field has a large and thorough comparison by Ferri et al. [4]. They test large number of evaluation metrics under different challenging situations with artificial datasets. Sokolova and Lapalme discuss on how measures are invariant towards the changes in the classifier results [5]. Seliya et al. and Ferri et al. compare similarities between evaluation measures [4, 6]. These comparisons, however, use mainly artificial datasets and focus on few challenging features at the time. Real data, on the other hand, consists a combination of the challenging features.

Furthermore, the classification structures, used in previous machine learning articles, differ significantly from GO. These articles discuss almost solely either binary or multinomial classification tasks, where each item cannot be correctly classified to many classes simultaneously. With GO prediction we have a task that reminds more multiple binary classification, where item can correctly belong to many classes or it might not belong to any of the available classes. In addition, GO prediction is further complicated by the correlations, created by the hierarchical structure of GO. All this creates more challenges to used EvMs.

Artificial Dilution Series, in comparison, has the following benefits:

1. We start with real-life data (with correct classifications), and propagate predictions to parental nodes with real Gene Ontology structure. This way we keep the real correlation structures and class size differences.
2. We know exactly how the different datasets should be ranked by evaluation metric. This simplifies analysis of the results.
3. Permuted datasets allow repetitive testing of evaluation metrics with large number of datasets having the same amount of error.
4. Dilution series lets us see the separation between intermediate amounts of noise. Not only difference between 0 vs 100% noise, as has been standard in the statistics and data mining research.

Points 2. and 3. have not been available in previous evaluations with real datasets [1, 3]. There analysis has usually been based on correlation between different data types and lacked repetitive tests that we present. Point

1. is difficult to replicate with fully artificial datasets, as all the features of real datasets are not always known. Furthermore, point 4 has not been used in this kind of testing before. Our results support point 4 and show that intermediate signal levels reveal more weaknesses in compared metrics than just the first and last signal level.

### 3 Signal and Noise in the ADS

Figure 1 represents here a toy example on the definition of signal and noise in ADS system. Figure shows a case where gene has one correct GO class. A signal class would be selected from the immediate neighborhood of the GO class. This is shown by a circle in our figure 1. Size of the circle is parameter in our rotation step. Noise class, on the other hand, is required to have very little overlapping tree path with the correct GO class. Therefore only the GO classes inside the box on the right would qualify as noise classes. This is the requirement that the noise classes must fulfill, when classes are swapped.

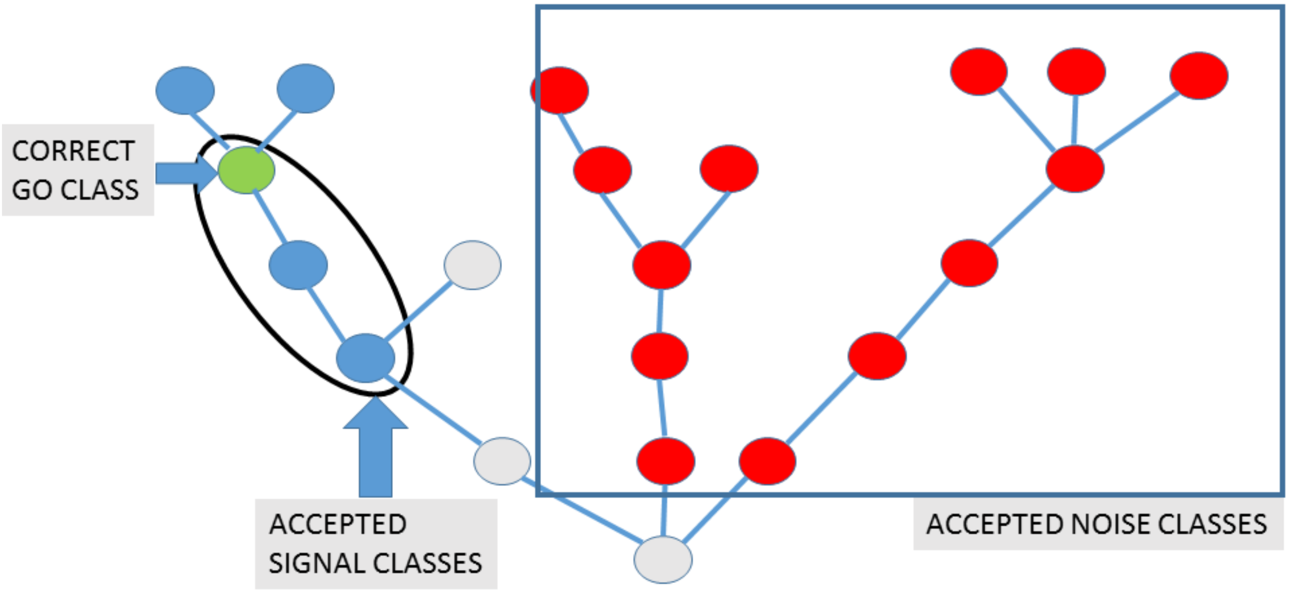


**Fig 1. Signal and Noise definition** Figure shows an example how signal and noise are defined in ADS. We have correct GO class, shown in green, in left branch. It's neighborhood in GO tree is shown in blue and we select either class itself or one of nearest parents as signal class. Noise class would be selected from distant branches, demonstrated with a red nodes.

### 4 Heterogeneity of use Evaluation Metrics

Literature has used quite different evaluation metrics for AFP methods. Table 1 demonstrates this variety. Metrics vary across the listed publications.

### 5 Evaluation metrics tested with ADS

This chapter describes evaluation metrics for Gene Ontology (GO) classifiers (hereinafter referred as "metrics") that we tested. These included metrics previously described in the literature as well as novel modifications of these metrics. Table 1 shows a collection of currently used metrics. In total we describe here 37 metrics including variations of Jaccard correlation, Receiver Operating Characteristics (ROC) Area Under Curve (AUC), precision-recall (PR)

| Article | Method evaluated | Publ.year | Evaluation Metric |
|---|---|---|---|
| Martin et al. [7] | Gotcha | 2004 | Selectivity vs. p-value<br>coverage vs. p-value |
| Engelhardt et al. [8] | Sifter | 2005 | ROC curve<br>error percentages |
| Götz et al. [9] | BLAST2GO | 2005 | Accuracy with threshold |
| Friedberg [10] | AFP 2005 | 2006 | Resnik semantics |
| Hawkins et al. [11] | PFP | 2008 | Schlicker semantics |
| Wass et al. [12] | ConFunc | 2008 | Prec-Recall curve |
| Chitale et al. [13] | ESG | 2009 | Schlicker semantics<br>Precision and Recall values |
| Engelhardt et al. [14] | Sifter v.2 | 2011 | ROC curve<br>Prec-Recall curve |
| Fontana et al. [15] | ARGOT | 2012 | Prec-Recall curve |
| Minneci et al. [16] | FFPred 2.0 | 2013 | $F_{max}$<br>Prec-Recall curve<br>SimGIC |
| Radivojac et al. [17] | CAFA1 | 2013 | $F_{max}$<br>Prec-Recall curve<br>weighted Prec–Recall curves<br>TC AUC |
| Gillis, Pavlidis [18] | CAFA1, re-eval. | 2013 | Prec-Recall curve,<br>TC AUC<br>Resnik semantics<br>Lin semantics |
| Koskinen et al. [19] | PANNZER | 2015 | weighted Prec-Recall curve<br>Lin semantics |
| Kahanda et al. [20] | CAFA2, re-eval. | 2015 | $F_{max}$,<br>TC $F_{max}$ |
| Jiang et al. [21] | CAFA2 | 2016 | $F_{max}$,<br>Prec-Recall curve,<br>$S_{min}$,<br>TC AUC |

**Table 1. Overview of some Evaluation Metrics used in AFP literature**. Table represents selected AFP method papers, AFP competition papers and re-evaluations of competitions (re-eval.) Prec-Recall refers to Precision Recall. Semantics refers to semantic similarity measure. Other abbreviations are explained under the Evaluation Metrics section in Materials and Methods. Notice the variety of metrics used.

AUC, $F_{max}$, $S_{min}$ [22], SimUI [23], SimGIC [24], Resnik [25] and Lin [26]. We also test six different methods for summarizing pairwise GO-term similarities. Supplementary table 1 presents all these metrics and their abbreviations.

#### 5.1 General notations for metric definitions

We assume that a GO classifier outputs a set of gene annotations with scores. More technically GO classifier outputs a set of triples each consisting of three values: gene label $g$, GO class $go$ and a classifier prediction score $sc$. We refer to this entire set of predicted gene annotations as $P$. A subset of $P$ containing all annotations for gene $g = i$ is denoted $P_i$ and a subset of $P$ containing all annotations with GO class $go = j$ is denoted $P_j$. Subset of $P$ with annotations scored greater or equal to threshold $sc \geq th$ is denoted $P^{th}$. Subscripts $i$ and $j$ and superscript $th$ can be combined to denote subsets of $P$ or other annotation sets with desired conditions (e.g. $P_i^{th}$ is a subset of all annotations for gene $i$ with scores equal or greater then $th$). Finally, let $n$ denote the number of unique genes in $P$, $n(th)$ the number of unique genes in $P^{th}$, $m$ the number of unique GO classes in $P$ and $m(th)$ the number of unique GO classes in $P^{th}$.

Let's further assume that we also have a set of correct or true gene annotations $T$. Given $T$ and $P$ we can define true positive annotations, $TP = P \cap T$, false positive annotations, $FP = P \setminus T$, and false negative annotations, $FN = T \setminus P$.

For ROC AUC metrics we will also need the set of true negative annotations $N$. We define true negatives as the Cartesian product of the set of genes in $T$ and the set of GO classes in GO subtracted by the correct set $T$:
$N = genes(T) \times classes(GO) \setminus T$

#### 5.2 Simple metrics based on Jaccard index

Jaccard index or Jaccard similarity coefficient between predicted $P$ and true $T$ annotation sets at threshold $th$ is defined as

$$Jacc(P,T,th) = \frac{|P^{th} \cap T|}{|P^{th} \cup T|} = \frac{|TP^{th}|}{|TP^{th}| + |FP^{th}| + |FN^{th}|}$$

Jaccard index is a function of threshold $th$. In order to convert this into a scalar metric we can take maximum value over all thresholds. Basic Jaccard formula can be applied to evaluate a set of predicted annotations $P$ in three ways. First, we can treat all annotations $x \in P$ equally disregarding any grouping of individual annotations by shared genes or GO classes. This is our definition of unstructured Jaccard, $USJacc$

$$US\ Jacc = \max_{th} \frac{|TP^{th}|}{|TP^{th}| + |FP^{th}| + |FN^{th}|} \tag{1}$$

Second, we can calculate Jaccard index separately for each gene and take the average of these values. This metric has been proposed earlier as $SimUI$ by Gentleman [23]. To remain consistent in our terminology we refer to this metric as Gene-Centric Jaccard, $GCJacc$:

$$GC\ Jacc = \max_{th} \frac{1}{n(th)} \sum_{i=1}^{n} \frac{|TP_i^{th}|}{|TP_i^{th}| + |FP_i^{th}| + |FN_i^{th}|} \tag{2}$$

Third, we can interpret predictions $P$ as assignment of genes to GO classes. In this case we calculate Jaccard index separately for each GO class in $P$. This is our definition of Term-Centric Jaccard (omitted from analysis):

$$TC\ Jacc = \max_{th} \frac{1}{m(th)} \sum_{j=1}^{m} \frac{|TP_j^{th}|}{|TP_j^{th}| + |FP_j^{th}| + |FN_j^{th}|}$$

Note that we propagate the annotations to ancestor nodes with these Jaccard methods.

#### 5.3 Metrics based on ROC and PR curves

When evaluating a binary classifier that produces a list of scored predictions we are generally interested in estimating the level of false positive and false negative errors. Generally the number of false positives and false negatives depend on the score threshold and are in inverse relation to each other: by increasing threshold the number of false negatives decreases but the number of false positives increases. In classical ROC analysis this balance is quantified by plotting True Positive Rate (TPR) against False Positive Rate (FPR):

$$TPR(P,T,th) = \frac{|P^{th} \cap T|}{|T|} = \frac{|TP^{th}|}{|TP^{th}| + |FN^{th}|}$$

$$FPR(P,T,th) = \frac{|P^{th} \setminus T|}{|N|} = \frac{|FP^{th}|}{|FP^{th}| + |TN^{th}|}$$

Performance is quantified as area under ROC curve (ROC AUC). AUC is closely related to Mann-Whitney U-statistic and is an estimate of the probability that a binary classifier will rank an instance of the positive class higher than an instance of a negative class [27]. Let $S$ denote a set of correct annotations, $j$ the positive subset and $k$ the negative subset. Let $rank(x, S)$ denote ranks assigned by the classifier to $x \in S$.

Then ROC AUC for positive class $j$ and negative class $k$ is:

$$AUC(rank, j, k) = \frac{1}{|j||k|}\Big(\sum_{x \in j} rank(x, S) - \frac{|j|(|j|+1)}{2}\Big)$$

AUC is a metric for binary classification. Here we consider a number of ways to extend this to a multi-class problem. Hand and Till [27] define the M metric, which is an arithmetic average of pairwise class comparisons. When applied to GO this means an arithmetic average of all pairs of GO classes in $T$. Ferri et all [4] define the AUNU metric, which is an arithmetic mean of one-vs-all class comparisons. This definition is similar to that used in CAFA2 competition [28] where $j$ was set to one of GO-classes in $T$ and $k$ to a union class covering all genes not in $j$. Further, we suggest that $j$ can be set cover all annotations in $T$ and $k$ to all other possible annotations i.e. the $N$ set. These approaches lead to different AUC variants.

First, let us define positive class as the entire set of correct annotations $T$ and negative class as all other possible annotations $N$. We refer to this metric as the unstructured AUC:

$$US\ AUC(rank, T, N) = \frac{1}{|T||N|}\Big(\sum_{x \in T} rank(x, T \cap N) - \frac{|T|(|T|+1)}{2}\Big) \quad (3)$$

Second, we calculate similar metric gene-wise. We refer to this metrics as the gene-centric AUC:

$$GC\ AUC(rank, T, N) = \frac{1}{n}\sum_{i=1}^{n}\frac{1}{|T_i||N_i|}\Big(\sum_{x \in T_i} rank(x, T_i \cap N_i) - \frac{|T_i|(|T_i|+1)}{2}\Big) \quad (4)$$

Third variant is the arithmetic mean of one-vs-all class comparisons given in general terms by Ferri et all [4] (i.e. the AUNU metric) and applied to GO-classifiers in CAFA competitions. We refer to this as the term-centric AUC. Let $genes_j$ denote the subset of genes in $T$ annotated with GO-class $j$ and let $n_j$ denote the size of that subset. Then term-centric AUC is:

$$TC\ AUC(P, T) = \frac{1}{m}\sum_{j=1}^{m}\frac{1}{n_j * (n - n_j)}\Big(\sum_{x \in genes_j} rank(x, genes) - \frac{n_j * (n_j+1)}{2}\Big) \quad (5)$$

Another method to quantify the balance between false positives and false negatives is to plot precision against recall. Precision, $pr(th)$, is defined as the fraction of true positives from predicted and recall (syn. sensitivity), $rc(th)$, as the fraction of true positives from correct annotations:

$$pr(P,T,th) = \frac{|TP^{th}|}{|TP^{th}| + |FP^{th}|}$$

$$rc(P,T,th) = \frac{|TP^{th}|}{|TP^{th}| + |FN^{th}|}$$

Area under precision-recall curve (AUCPR) is a scalar metric that reflects classifiers overall accuracy. Several authors have argued that AUCPR is better suited for classification involving class imbalance problem (for a discussion see [29]). We note that both $US\ AUC$ and $GC\ AUC$ have negative class manyfold larger then the positive class and are thus susceptible to class imbalance.

In our implementation we calculated AUCPR from $pr(th)$ and $rc(th)$ curves using trapezoidal method [29]. As with ROC AUC, basic AUCPR formula can be applied to evaluated a set of predicted GO annotations in three ways. For unstructured AUCPR we treat all $x \in P$ equally disregarding any grouping of individual annotations by shared genes or GO classes. For gene-centric AUCPR we calculate $pr$ and $rc$ separately for each gene and calculate the arithmetic mean of these values prior to AUC calculus. For term centric AUCPR we calculate $pr$ and $rc$ for each GO class separately and calculate the arithmetic mean of these values prior to AUC calculus. Definitions for variations of precision, recall and AUCPR are then

$$pr_{us}(th) = \frac{|TP^{th}|}{|TP^{th}| + |FP^{th}|}$$

$$rc_{us}(th) = \frac{|TP^{th}|}{|TP^{th}| + |FN^{th}|}$$

$$US\ AUCPR(P,T) = AUC(pr_{us}(th), rc_{us}(th)) \tag{6}$$

$$pr_{gc}(th) = \frac{1}{n(th)} \sum_{i=1}^{n} \frac{|TP_i^{th}|}{|TP_i^{th}| + |FP_i^{th}|}$$

$$rc_{gc}(th) = \frac{1}{n(th)} \sum_{i=1}^{n} \frac{|TP_i^{th}|}{|TP_i^{th}| + |FN_i^{th}|}$$

$$GC\ AUCPR(P,T) = AUC(pr_{gc}(th), rc_{gc}(th)) \tag{7}$$

$$pr_{tc}(th) = \frac{1}{m(th)} \sum_{j=1}^{m} \frac{|TP_j^{th}|}{|TP_j^{th}| + |FP_j^{th}|}$$

$$rc_{tc}(th) = \frac{1}{m(th)} \sum_{j=1}^{m} \frac{|TP_j^{th}|}{|TP_j^{th}| + |FN_j^{th}|}$$

$$TC\ AUCPR(P,T) = AUC(pr_{tc}(th), rc_{tc}(th)) \tag{8}$$

F-metric is a simple metric based on precision and recall, or the harmonic mean of precision and recall at fixed threshold. It's maximum over all the thresholds, $F_{max}$ metric, is a popular implementation of it, and has been used extensively in CAFA competitions [17, 28] Note that the precision and recall values are calculated gene-wise at each threshold $th$ and average values $pr_{gc}(th)$ and $rc_{gc}(th)$ are used:

$$GC\ F_{max} = \max_{th} \frac{2 \times pr_{gc}(th) \times rc_{gc}(th)}{pr_{gc}(th) + rc_{gc}(th)} \tag{9}$$

Ferri et al. [4] considered an alternative usage for F-measure, where an average is calculated across all predicted classes. This would correspond to our Term Centric analysis. In their analysis the threshold was predefined for the analysis.

#### 5.4 Information content

One major challenge with GO structure is the variation in the class sizes we see in the data. This causes the naive prediction of largest GO classes perform well on comparisons (see main article). This problem can be corrected by adding weights, called *Information Contents* to the class predictions that emphasize the smaller more meaningful classes. Resnik defined the information content of an individual GO class $x$ as the negative log likelihood of $x$ [25]. Likelihoods $p(x)$ for GO classes can be estimated from a corpus of GO annotations such as UniProt Gene Ontology Annotation database (UniProt-GOA). Thus information content of a GO class $x$ can be defined in terms of it's frequency $f(x)$ in a given database:

$$ic(x) = log\frac{1}{p(x)} = log\frac{1}{f(x)}$$

Clark and Radivojac defined information content of GO subgraph $G$ as a negative log likelihood of $G$ [22]. Let $\mathcal{P}(x)$ denote the set of immediate parents of $x$ in the GO graph. Then the likelihood of subgraph $G$ can be factorized as a product of conditional probabilities, $p(G) = \prod_{x \in G} p(x|\mathcal{P}(x))$ [22]. Conditional probabilities $p(x|\mathcal{P}(x))$ and a quantity corresponding to $ic(x)$ can be estimated from a given database (e.g. from UniProt-GOA):

$$ic2(x) = log\frac{1}{p(x|\mathcal{P}(x))}$$

Information content of a subgraph $G$ is then defined as a sum of $ic2(x)$ values for all $x \in G$:

$$ic(G) = S_{ic2}(G) = \sum_{x \in G} ic2(x)$$

For the sake of comparison we also used a simpler definition of subgraph information content that is based on $ic(x)$ values:

$$ic'(G) = S_{ic}(G) = \sum_{x \in G} ic(x)$$

The motivation for using both $ic(G)$ and $ic'(G)$ is to evaluate how different definitions will effect metric performance. When both definitions are used we label metrics based on $ic(G)$ with "ic2." prefix and metrics based on $ic'(G)$ with "ic." prefix according to the $ic(x)$ or $ic2(x)$ values that are used in the calculus.

#### 5.5 Metrics incorporating information content

Pesquita et al. introduced an $ic$ weighted Jaccard index [24]:

$$SimGIC(P,T) = \max_{th} \frac{1}{n} \sum_{i=1}^{n} \frac{S_{ic}(TP_i^{th})}{S_{ic}(TP_i^{th}) + S_{ic}(FP_i^{th}) + S_{ic}(FN_i^{th})} \quad (10)$$

Note that gene-wise quotients in this equation are ratios that can get similar values for genes with very different number of annotations. This can overweight genes that have few annotations and underweight genes with many annotations. To overcome this limitation we introduce a modified version that treats all annotations equally as in unstructured AUC and Jaccard metrics:

$$SimGIC2(P,T) = \max_{th} \frac{S_{ic}(TP^{th})}{S_{ic}(TP^{th}) + S_{ic}(FP^{th}) + S_{ic}(FN^{th})} \quad (11)$$

Clark and Radivojac defined remaining uncertainty, $ru$, as the average gene-wise information content of the $FN$ set and misinformation, $mi$, as the average gene-wise information content of $FP$ set. Based on these, they defined $S_{min}$ metric (labeled here as $S_{min1}$) [22]:

$$ru(th) = \frac{1}{n} \sum_{i=1}^{n} S_{ic2}(FN_i^{th}) = \frac{1}{n} \sum_{i=1}^{n} \sum_{x \in FN_i^{th}} ic2(x)$$

$$mi(th) = \frac{1}{n} \sum_{i=1}^{n} S_{ic2}(FP_i^{th}) = \frac{1}{n} \sum_{i=1}^{n} \sum_{x \in FP_i^{th}} ic2(x)$$

$$S_{min1}(P,T) = \min_{th} \sqrt{ru(th)^2 + mi(th)^2} \quad (12)$$

Note that $S_{min1}$ has no gene-centric terms in it's equation. At the core of this metric is the sum of $ic(FN_i^{th})$ and $ic(FP_i^{th})$ terms across all genes. This makes $S_{min1}$ insensitive to distribution of misclassification errors across genes and is by this property similar to other *unstructured* metrics. To test how gene-centric terms would effect this metric we introduce a variation of $S_{min1}$ by changing the order of arithmetic average and Euclidean distance calculus:

$$S_{min2}(P,T) = \min_{th} \frac{1}{n} \sum_{i=1}^{n} \sqrt{S_{ic}(FN_i^{th})^2 + S_{ic}(FP_i^{th})^2} \tag{13}$$

While considering possible variations for $S_{min}$ calculus we noticed that $1/n$ terms can be moved outside the squere root. Thus, by multiplying $S_{min1}$ by $n$ (the number of genes) we can define a simpler metric that has the same variance and ADS performance as $S_{min1}$:

$$\begin{aligned} S_{min1}(P,T) &= \min_{th} \sqrt{\left(\frac{1}{n}\sum_{i=1}^{n} S_{ic}(FN_i^{th})\right)^2 + \left(\frac{1}{n}\sum_{i=1}^{n} S_{ic}(FP_i^{th})\right)^2} \\ &= \frac{1}{n} \min_{th} \sqrt{S_{ic}(FN^{th})^2 + S_{ic}(FP^{th})^2} \end{aligned}$$

$$S_{min3}(P,T) = S_{min1}(P,T) * n = \min_{th} \sqrt{\left(S_{ic}(FN^{th})^2 + S_{ic}(FP^{th})^2\right)}$$

Furthermore, the CAFA organizers have considered a weighted version of F metric [28]. This uses sums of Information Content weights instead of counts of predictions. We excluded this from our current comparisons as we had to limit the number of compared metrics.

#### 5.6 Metrics based on pairwise semantic similarities

Metrics listed to this point will only differentiate between exact matches and mismatches of predicted versus correct GO classes. However, using semantic similarity measures it is possible to define metrics that are aware of the relationships between mismatching GO classes in the ontology tree [24].

Semantic similarity metrics are generally gene-centric and have at their core pairwise comparison of predicted GO classes against correct GO classes. We implemented three core functions: *Resnik* [25], *Lin* [26] and Ancestor Jaccard index ($AJacc$). Let $x \in GO$ and $y \in GO$ be any two nodes in GO graph. And let $A(x)$ and $A(y)$ denote the sets of all ancestors in $GO$ for nodes $x$ and $y$. Then the most informative common ancestor of $x$ and $y$ is defined by $MICA(x,y)$ [24] (equivalent to *minimum subsummer* defined by Lord [30]):

$$MICA(x,y) = \operatorname*{argmax}_{z \in A(x) \cap A(y)} ic(z)$$

The three core semantic similarity metrics are then defined by equations:

$$Resnik(x,y) = ic(MICA(x,y))$$

$$Lin(x,y) = \frac{2 \times ic(MICA(x,y))}{ic(x) + ic(y)}$$

$$AJacc(x,y) = \frac{|A(x) \cap A(y)|}{|A(x) \cup A(y)|}$$

Now let $X$ denote the ordered list of predicted GO classes for gene $i$ at threshold $th$ and $Y$ the list of correct GO classes. Predicted GO classes are ordered using the classifiers prediction score. Let $sem(x,y)$ be any core semantic similarity function between an arbitrary pair of GO classes $x \in GO$ and $y \in GO$. Then pairwise semantic similarities for gene $i$ at threshold $th$ can be summarized by a similarity matrix, with rows standing for predicted and columns for correct GO classes:

$$SIM(k,l) = sem(X(k), Y(l))$$

See table 1 for an example on this.

##### 5.7 Combining semantic similarities into a single score

To devise a scalar scoring function, we need to summarize SIM matrix into a scalar value (see table 1). We propose six ways to do this: mean value of SIM (method $A$), mean value of column maxima of SIM (method $B$), mean value of row maxima of SIM (method $C$), mean value of methods B and C (method $D$), min value of methods B and C (method $E$), mean value of pooled column and row maxima of SIM (method $F$).

$$A(X,Y,sem) = \frac{1}{|X| \times |Y|} \sum_{x \in X} \sum_{y \in Y} sem(x,y)$$

$$B(X,Y,sem) = \frac{1}{|Y|} \sum_{y \in Y} \max_{x \in X} sem(x,y)$$

$$C(X,Y,sem) = \frac{1}{|X|} \sum_{x \in X} \max_{y \in Y} sem(x,y)$$

$$D(X,Y,sem) = \frac{B(X,Y,sem) + C(X,Y,sem)}{2}$$

$$E(X,Y,sem) = \min\{B(X,Y,sem), C(X,Y,sem)\}$$

$$F(X,Y,sem) = \frac{1}{|X| + |Y|} \big( \sum_{x \in X} \max_{y \in Y} sem(x,y) + \sum_{y \in Y} \max_{x \in X} sem(x,y) \big)$$

Method A treats all pairwise comparisons equally and is equivalent to the all-pair arithmetic average proposed by Lord [30]. Method B is the best-match average for correct GO-classes. As such this method is sensitive to false negative errors and insensitive to false positive errors. Method C is the best-match average for predicted classes and is thus sensitive to false positive errors and insensitive to false negative errors (see discussion in the next section). Methods B and C are asymmetrical and this is corrected with method D. Best-match average methods were previously described by several authors [24, 31–33].

As B can only monitor false negatives and C only the false positives, they are expected to represent weak performance in the analysis. We include them into the analysis as negative controls to see if ADS can detect their performance differences. We further introduce a novel methods $E$ and $F$, that both monitor false negatives and false positives.

Methods $A$ to $F$ are functions of threshold $th$ and gene $i$. To convert these into scalar values for the predicted annotation $P$ we calculate the selected method for each threshold $th$ and select the maximum value across all possible thresholds. Let $sem$ be any core semantic similarity function and $S$ any summation method for the similarity matrix. Semantic similarity metric $EM_{sem}$ is then:

$$EM_{sem}(P,T) = \max_{th} \frac{1}{n(th)} \sum_{i}^{n} S(P_i^{th}, T_i, sem)$$

We combined three $sem$ functions with the six $S$ functions to defined altogether 18 variations of semantic similarity metrics. In the following section we give equations for $EM_{sem}$ based on $Resnik$. Equations for $EM_{sem}$ based on $Lin$ and $AJacc$ are similar.

$$Resnik\ A(P,T) = \max_{th} \frac{1}{n(th)} \sum_{i}^{n} A(P_i^{th}, T_i, Resnik) \tag{14}$$

$$Resnik\ B(P,T) = \max_{th} \frac{1}{n(th)} \sum_{i}^{n} B(P_i^{th}, T_i, Resnik) \tag{15}$$

$$Resnik\ C(P,T) = \max_{th} \frac{1}{n(th)} \sum_{i}^{n} C(P_i^{th}, T_i, Resnik) \tag{16}$$

$$Resnik\ D(P,T) = \max_{th} \frac{1}{n(th)} \sum_{i}^{n} D(P_i^{th}, T_i, Resnik) \tag{17}$$

$$Resnik\ E(P,T) = \max_{th} \frac{1}{n(th)} \sum_{i}^{n} E(P_i^{th}, T_i, Resnik) \tag{18}$$

$$Resnik\ F(P,T) = \max_{th} \frac{1}{n(th)} \sum_{i}^{n} F(P_i^{th}, T_i, Resnik) \tag{19}$$

##### 5.8 Comparing semantic similarity summation methods

We demonstrate the principle of summation methods for semantic similarities with a toy dataset in table 1. We explain strengths and weaknesses of each method especially when the final score is the maximum score over threshold $th$ positions.

First, it is easy to see that the average of column maxima is expected to grow as the threshold goes lower even with a randomly sampled matrix, as each maximum is selected from larger set of numbers. This means that when the maximum score over $th$ for method B is selected, it will be always at lowest positions. So method B clearly *cannot monitor false positives.* Therefore, average of row maxima would do better job at separating random matrix from one with signal at the lower $th$ positions.

Situation is totally opposite at the highest threshold positions. When only a single row is selected from randomly sampled matrix, one strong value can alter the result for row maximums. This means that the maximum score over $th$ with method C will most likely at higher $th$ positions. However, now we are not paying attention on how many correct GO classes get annotated. Therefore method C clearly *cannot monitor false negatives.*

Method A has a different problem. If there is only one GO class in the correct set, then it is possible to obtain good score for correct prediction at high $th$ position. However, assume that a gene has ten correct GO classes and these are very dissimilar from each other in GO tree. Now if classifier predicts the very same GO classes in top-10 positions, 90% of the values in the matrix will represent dissimilarities between GO-classes. Therefore method A is clearly *affected by the number and the heterogeneity of the correct GO classes.*

Taking an average of B and C, like in method D, mixes weaker and better signal. This might not be good choice when maximum score over $th$ is selected. Indeed, our results show that D is quite weak method.

Concatenated row and column maxima, as in method F, will always emphasize the longer vector of two vectors. Therefore, it will pay more attention to column maxima at the higher threshold positions and more attention row maxima at the lower threshold positions. Selecting the minimum of B and C, as in method E, requires that both mean of row and column maxima show strong signal. This is more challenging to row maxima at lower positions and to column maxima at higher positions. Therefore we propose methods E and F as new summation methods for semantic similarities.

| | | | | Correct GO classes | | | | | |
|---|---|---|---|---|---|---|---|---|---|
| | | | | GO-1 | GO-2 | GO-3 | GO-4 | GO-5 | |
| | | classifier scores | Row Max | 0.6 | 0.2 | 0 | 0.7 | 1 | Column Max |
| Predicted GO classes | GO-6 | 0.9 | 0.8 | 0 | 0.2 | 0 | 0.6 | 0.8 | |
| | GO-7 | 0.86 | 0.6 | 0.6 | 0 | 0 | 0 | 0 | |
| | GO-5 | 0.84 | 1 | 0 | 0 | 0 | 0.7 | 1 | |
| | GO-8 | 0.8 | 0.2 | 0.1 | 0.2 | 0 | 0.2 | 0 | $thr = 0.8$ |
| | GO-9 | 0.76 | 0.8 | 0 | 0.2 | 0.8 | 0 | 0 | |
| | GO-1 | 0.74 | 1 | 1 | 0 | 0.1 | 0 | 0 | |
| | GO-10 | 0.74 | 0.2 | 0 | 0.2 | 0 | 0.2 | 0 | |
| | GO-11 | 0.7 | 0.3 | 0.1 | 0 | 0.2 | 0.1 | 0.3 | |

**(a)**

| Summation methods for Semantic Similarities | | | Score for threshold | |
|---|---|---|---|---|
| Method abbr. | Description | Equation | $thr = 0.8$ | $thr = 0.7$ |
| A | matrix mean | $\sum_j \sum_i x_{ij}/(NM)$ | 0.22 | 0.19 |
| B | mean of col.maxima | $\sum_{j=1}^{M} \max_{1 \leq i \leq N} x_{ij}/M$ | 0.5 | 0.74 |
| C | mean of row maxima | $\sum_{i=1}^{N} \max_{1 \leq j \leq M} x_{ij}/N$ | 0.575 | 0.538 |
| D | mean of B and C | $(B + C)/2$ | 0.538 | 0.639 |
| E | min of B and C | $min(B, C)$ | 0.5 | 0.538 |
| F | mean of concatenated row and col. maxima | $(\sum_{j=1}^{M} \max_{1 \leq i \leq N} x_{ij} + \sum_{i=1}^{N} \max_{1 \leq j \leq M} x_{ij})/(N + M)$ | 0.533 | 0.615 |

**(b)**

**Table 2.** Two tables demonstrate the different summation methods for GO semantic similarities, used here. First part shows small toy data, where rows represent predicted GO classes and columns represent correct GO classes. Predicted GO classes are ordered using the classifiers prediction score. Next, a semantic similarity score is calculated for each GO class pair in table and a threshold value *thr* is defined to select accepted predictions. Summary method then takes the selected upper subset of the table as an input. Lower table shows different summation methods that we tested. We show results for two threshold settings from toy data. In our analysis we processed large set of threshold values and selected the largest similarity score.
