## Supplementary Table 1 for "Novel Comparison of Evaluation Metrics for Gene Ontology Classifiers Reveals Drastic Performance Differences"

| Type | Abbreviation | Core function | Data summary | Thresh. function | IC weight | SemSim. Sum. | Popular | Neg. Ctrl | Novel | Simple EvM |
|---|---|---|---|---|---|---|---|---|---|---|
| AUC metrics | US AUC | Area Under ROC Curve | Unstructured | - | - | - | | + | | + |
| | GC AUC | Area Under ROC Curve | Gene-Centric | - | - | - | | | | |
| | TC AUC | Area Under ROC Curve | Term-Centric | - | - | - | + | | | |
| | US AUCPR | Area Under Prec-Recall Curve | Unstructured | - | - | - | + | | | + |
| | GC AUCPR | Area Under Prec-Recall Curve | Gene-Centric | - | - | - | | | | |
| | TC AUCPR | Area Under Prec-Recall Curve | Term-Centric | - | - | - | | | | |
| | Fmax | F-metric | Gene-Centric | max | - | - | + | | | |
| group metrics | US Jacc | Jaccard | Unstructured | max | - | - | | | | + |
| | GC Jacc | Jaccard | Gene-Centric | max | - | - | | | | + |
| | ic Smin1 | Euclidean distance | Unstructured | max | IC1 | - | | | | |
| | ic Smin2 | Euclidean distance | Gene-Centric | max | IC1 | - | | | + | |
| | ic Smin3 (excluded) | Euclidean distance | Unstructured | max | IC1 | - | | | + | |
| | ic SimGIC | Weighted Jaccard | Gene-Centric | max | IC1 | - | | | | |
| | ic SimGIC2 | Weighted Jaccard | Unstructured | max | IC1 | - | | | + | |
| | ic2 Smin1 | Euclidean distance | Unstructured | max | IC2 | - | + | | | |
| | ic2 Smin2 | Euclidean distance | Gene-Centric | max | IC2 | - | | | + | |
| | ic2 Smin3 (excluded) | Euclidean distance | Gene-CEntric | max | IC2 | - | | | + | |
| | ic2 SimGIC | Weighted Jaccard | Gene-Centric | max | IC2 | - | | | | |
| | ic2 SimGIC2 | Weighted Jaccard | Unstructured | max | IC2 | - | | | + | |
| Semantic Similarities | Resnik score A | Resnik semantics | Gene-Centric | max | IC1 | A | | | | |
| | Resnik score B | Resnik semantics | Gene-Centric | max | IC1 | B | | + | | |
| | Resnik score C | Resnik semantics | Gene-Centric | max | IC1 | C | | + | | |
| | Resnik score D | Resnik semantics | Gene-Centric | max | IC1 | D | | | | |
| | Resnik score E | Resnik semantics | Gene-Centric | max | IC1 | E | | | + | |
| | Resnik score F | Resnik semantics | Gene-Centric | max | IC1 | F | | | + | |
| | Lin score A | Lin semantics | Gene-Centric | max | IC1 | A | | | | |
| | Lin score B | Lin semantics | Gene-Centric | max | IC1 | B | | + | | |
| | Lin score C | Lin semantics | Gene-Centric | max | IC1 | C | | + | | |
| | Lin score D | Lin semantics | Gene-Centric | max | IC1 | D | | | | |
| | Lin score E | Lin semantics | Gene-Centric | max | IC1 | E | | | + | |
| | Lin score F | Lin semantics | Gene-Centric | max | IC1 | F | | | + | |
| | AJacc score A | Jaccard semantics | Gene-Centric | max | - | A | | | | |
| | AJacc score B | Jaccard semantics | Gene-Centric | max | - | B | | + | | |
| | AJacc score C | Jaccard semantics | Gene-Centric | max | - | C | | + | | |
| | AJacc score D | Jaccard semantics | Gene-Centric | max | - | D | | | | |
| | AJacc score E | Jaccard semantics | Gene-Centric | max | - | E | | | + | |
| | AJacc score F | Jaccard semantics | Gene-Centric | max | - | F | | | + | |
