## Supplementary Table 2 for "Novel Comparison of Evaluation Metrics for Gene Ontology Classifiers Reveals Drastic Performance Differences"

A

| | Compared parameters | | Correlation | |
|---|---|---|---|---|
| | 1st K | 2nd K | Pearson | rank |
| RC.uniprot.1000 | 2 | 3 | 0.998587 | 0.997759 |
| RC.cafa2012 | 2 | 3 | 0.997951 | 0.998599 |
| RC.mouseFunc | 2 | 3 | 0.998517 | 0.998319 |
| RC.uniprot.1000 | 2 | 4 | 0.994401 | 0.996639 |
| RC.cafa2012 | 2 | 4 | 0.994472 | 0.996919 |
| RC.mouseFunc | 2 | 4 | 0.995352 | 0.996078 |
| FPS.uniprot.1000 | 2 | 3 | 0.999746 | 1 |
| FPS.cafa2012 | 2 | 3 | 0.99971 | 0.999368 |
| FPS.mouseFunc | 2 | 3 | 0.9999 | 0.997467 |
| FPS.uniprot.1000 | 2 | 4 | 0.999633 | 0.999507 |
| FPS.cafa2012 | 2 | 4 | 0.999105 | 0.999438 |
| FPS.mouseFunc | 2 | 4 | 0.999765 | 0.993526 |
| | | Minimum | 0.994401 | 0.993526 |
| | | Maximum | 0.9999 | 1 |

Comparison of results when K parameter is varied
Comparison is done by calculating correlation between two sets of results
RC and FPS results are evaluated separately

B

| | Compared parameters | | correlation | |
|---|---|---|---|---|
| | jacc limit 1 | Jacc limit 2 | Pearson | Rank |
| RC.uniprot.1000 | 0.2 | 0.8 | 0.99985 | 0.99972 |
| RC.cafa2012 | 0.2 | 0.8 | 0.999907 | 0.998599 |
| RC.mouseFunc | 0.2 | 0.8 | 0.999198 | 0.998319 |
| FPS.uniprot.1000 | 0.2 | 0.8 | 0.999664 | 0.99698 |
| FPS.cafa2012 | 0.2 | 0.8 | 0.999696 | 0.994526 |
| FPS.mouseFunc | 0.2 | 0.8 | 0.999608 | 0.997748 |
| | | Minimum | 0.999198 | 0.994526 |
| | | Maximum | 0.999907 | 0.99972 |

Comparison of results when jacc-threshold in noise generation is varied
Comparison is done by calculating correlation between two sets of results
RC and FPS results are evaluated separately
